# The context dependent effects of host competence, competition, and the pathogen transmission mode on disease prevalence

**DOI:** 10.1101/2020.01.31.928820

**Authors:** Michael H. Cortez, Meghan A. Duffy

**Author notes:** Statement of Authorship: MHC and MAD designed the study; MHC did mathematical analysis and wrote first version of the paper; MAD contributed to writing. Data Statement: No new data generated in this study. Calculations and equations are provided in the appendices. The authors wish to be identified to the reviewers. Supplementary Material: Online Appendices S1-S4.

## Abstract

Biodiversity in communities is changing globally, including the gain and loss of host species in host-pathogen communities. Increased host diversity can cause infection prevalence in a focal host to increase (amplification) or decrease (dilution). However, it is unclear what general rules govern the context dependent effects, in part because theories for pathogens with different transmission modes have developed largely independently. Using a two-host model, we explore how the pathogen transmission mode and characteristics of a second host (disease competence and competitive ability) influence disease prevalence in a focal host. Our work shows how the theories for pathogens with environmental transmission, density-dependent direct transmission, and frequency-dependent direct transmission can be unified. Our work also identifies general rules about how host and pathogen characteristics affect amplification/dilution. For example, higher competence hosts promote amplification, unless they are strong interspecific competitors; strong interspecific competitors promote dilution, unless they are large sources of new infections; and dilution occurs under frequency-dependent direct transmission more than density-dependent direct transmission, unless interspecific host competition is sufficiently strong. Our work helps explain how the characteristics of the pathogen and a second host affect disease prevalence in a focal host.

## 1 Introduction

Biodiversity in communities is changing across the globe, with species introductions in some and extirpation in others. Altered biodiversity can influence patterns of infectious diseases because most pathogens can infect — and most communities are made up of — multiple host species (Cleaveland et al., 2001; Pedersen et al., 2005; Rigaud et al., 2010). These effects of biodiversity on disease levels result from the joint effects of how each host species interacts with the pathogen (e.g., within-species transmission) and interspecific interactions between host species (e.g., resource competition and between-species transmission). For example, increased snail species richness leads to greater interspecific host competition, which then causes decreased prevalence (i.e., the proportion of infected individuals in a population) of the trematode parasite *Ribeiroia ondatrae* (Johnson et al., 2012). Similarly, while greater rodent diversity increases transmission between individuals, the reductions in host density due to interspecific competition result in an overall decrease in the prevalence of Sin Nombre hantavirus (Luis et al., 2018). In contrast, native and invasive daphniid hosts of the fungal pathogen *Metchnikowia bicuspidata* compete for resources, but both hosts cause increased infection prevalence in the other (Searle et al., 2016). These contrasting impacts of altered species diversity highlight the need to better understand the biological mechanisms that promote higher versus lower prevalence in a focal host as host biodiversity changes.

The dilution effect argues that increased host biodiversity decreases disease prevalence (i.e., the proportion of infected individuals in a population) (Keesing et al., 2006). However, when and whether increased host biodiversity reduces disease prevalence (dilution) or increases disease prevalence (amplification) in a focal host population has been debated in the literature (e.g., Lafferty and Wood 2013; Ostfeld and Keesing 2013; Wood and Lafferty 2013 and reviewed in Rohr et al. 2020). Empirical evidence is mixed: a recent meta-analysis found general empirical support for dilution (Civitello et al., 2015), but amplification also occurs (Salkeld et al., 2013; Wood et al., 2014; Venesky et al., 2014). This implies that increased host biodiversity likely has context-dependent effects (Salkeld et al., 2013), motivating calls for theory that identifies specific biological mechanisms promoting amplification versus dilution (Buhnerkempe et al., 2015; Halsey, 2019; Rohr et al., 2020).

Current theory (Keesing et al., 2006, 2010) predicts amplification versus dilution depends on how host biodiversity affects host-pathogen encounter rates, susceptible host densities, and transmission, mortality and recovery rates. Specific mechanisms affecting these factors include host competence (i.e., the host’s ability to transmit the pathogen), intraspecific and interspecific host competition, and the pathogen transmission mechanism. For the latter, some studies have focused on pathogens with either (i) frequency-dependent direct transmission (transmission via host-host contact that depends on the frequency of infected individuals), (ii) density-dependent direct transmission (transmission via host-host contact that depends on the density of infected individuals), or (iii) environmental transmission (transmission via contact with infectious propagules, e.g., spores, excreted by infected individuals). Many studies suggest that frequency-dependent direct transmission promotes dilution whereas density-dependent direct transmission and environmental transmission promote amplification (Begon et al., 1992; Begon and Bowers, 1994; Dobson, 2004; Rudolf and Antonovics, 2005; Mihaljevic et al., 2014; Faust et al., 2017; Roberts and Heesterbeek, 2018). However, theoretical studies also show that accounting for interspecific host competition (Strauss et al., 2015; O’Regan et al., 2015; Searle et al., 2016) and host competence (Rudolf and Antonovics, 2005; O’Regan et al., 2015; Roberts and Heesterbeek, 2018) can qualitatively alter predictions. For example, introduction of a high competence host can cause dilution (i.e., *lower* prevalence) in a focal host, even for density-dependent direct transmission pathogens (O’Regan et al., 2015; Roberts and Heesterbeek, 2018) or environmental transmission (Searle et al., 2016).

This variation in predictions is, in part, due to three reasons. First, studies use different metrics to quantify levels of disease, including disease prevalence in a focal host or the community, the density of infected individuals in a focal host or the community, and the pathogen’s basic reproductive number (*ℛ*_0_). Predictions about biodiversity-disease relationships can disagree when different metrics are used (Roche et al., 2012; Roberts and Heesterbeek, 2018). Second, studies vary different quantities in order to assess how levels of disease respond to changes in host biodiversity. Three common approaches are to measure responses to (i) varying model parameters, (ii) varying the density of non-focal hosts, or (iii) the addition or removal of a host species. The approaches are mathematically related, but related work on indirect effects in predator-prey systems has shown that the three approaches can make qualitatively different predictions (Abrams and Nakajima, 2007; Abrams and Cortez, 2015). Third, studies make different assumptions about community make-up and species interactions. This includes whether there are two or more host species, whether some hosts are non-competent (e.g., a host can become infected, but cannot transmit the pathogen), whether interspecific host interactions are present or absent, and whether there is direct, environmental, or vector-borne transmission. Given this variation across studies, there is a clear need for new theory that can unify existing theory and provide general predictions about how specific mechanisms shape host biodiversity-disease relationships.

The goal of this study is to address two key limitations of the current theory. First, the theories for pathogens with different transmission modes have developed largely independently. This makes it difficult to compare predictions across models and isolate how the pathogen transmission mode influences disease prevalence. Second, it is unclear what general rules govern the context dependent effects the pathogen transmission mode and host species characteristics (e.g., competence and competitive ability) have on amplification and dilution. To begin to address these limitations, we present a two-host model with environmental transmission of a pathogen and use it to unify the theories for environmental transmission, density-dependent direct transmission and frequency-dependent direct transmission. We then explore how the pathogen transmission mode and characteristics of a second host (specifically, disease competence and interspecific and intraspecific competitive abilities) influence whether a second host increases or decreases infection prevalence in a focal host. Our work yields insight into the context dependent effects an additional host has on disease prevalence. These findings point to possible explanations for the variation in prior theoretical and empirical studies on the effects of host biodiversity on disease prevalence.

## 2 Models and Methods

### 2.1 Two-host-one-pathogen environmental transmission model

We model a system with two host species and an environmentally transmitted pathogen. Examples of environmentally transmitted pathogens include human pathogens such as *Vibrio cholera* and *Giardia lamblia* (Kaper et al., 1995; Wolfe, 1992) and wildlife diseases like whirling disease in fish (Hedrick et al., 1998; Bartholomew and Reno, 2002) and trematode parasites of snails (Johnson et al., 2012).

Our environmental transmission model describes the changes in the densities of susceptible (*S*_*i*_) and infected (*I*_*i*_) individuals in two host populations (*i* = 1, 2) and the density of infectious propagules (*P*) in the environment. Throughout, we refer to host species 1 and 2 as the ‘focal’ and ‘second’ host, respectively. New infections arise when susceptible individuals come in contact with infectious propagules that were released by infected individuals. We assume the two host species share the same habitat, excrete infectious propagules into the same pathogen pool, and are equally exposed to the pathogen pool. To simplify the presentation and analysis in the main text, we assume there is no recovery from infection (i.e., infection is always lethal). In appendix S2 we analyze a model with recovery and show all of our results hold.

The model is,

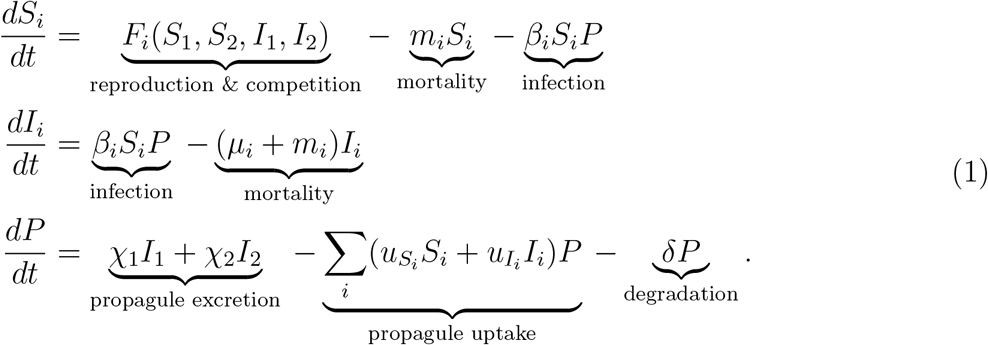

In the model, susceptible individuals increase due to reproduction (*F*_*i*_(*S*_1_, *S*_2_, *I*_1_, *I*_2_)), die due to sources other than disease (*m*_*i*_*S*_*i*_), and become infected when they come in contact with infectious propagules (*β*_*i*_*S*_*i*_*P*); infected individuals die due to disease and sources other than disease (*μ*_*i*_*I*_*i*_ and *m*_*i*_*I*_*i*_, respectively); and infectious propagules are excreted into the environment by infected individuals (*χ*_*i*_*I*_*i*_) and lost due to uptake by all hosts 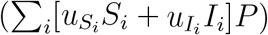 and degradation (*δP*). The transmission coefficient (*β*_*i*_) and susceptible individual uptake rate 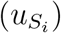 for population *i* are related because the transmission coefficient is the product of the uptake rate and the per propagule probability of infection 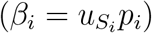. This constrains the parameters such that 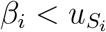. In our analyses, we vary *β*_*i*_ independent of 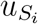 by varying the per propagule probability of infection and we vary 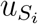 independent of *β*_*i*_ by accounting for changes in 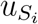 with compensatory increases or decreases in the per propagule probability of infection.

The reproduction rates (*F*_*i*_) account for reproductive output from both susceptible and infected individuals and the effects of intraspecific and interspecific host competition. We use the general functions in order to develop theory that applies across systems. In our numerical examples we use the Lotka-Volterra competition functions,

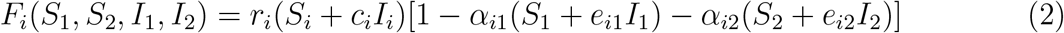

where *r*_*i*_ and *c*_*i*_*r*_*i*_ are the maximum exponential growth rates of susceptible and infected individuals of species *i, α*_*ij*_ is the per capita competitive effect of host *j* on host *i*, and *e*_*ij*_ determines if infected individuals of host *j* have weaker (*e*_*ij*_ < 1), equal (*e*_*ij*_ = 1), or stronger (*e*_*ij*_ > 1) competitive effects on host *i* than susceptible individuals of host *j*. Infected individuals might be stronger competitors when infection causes increased appetite or resource acquisition rates (Ponton et al., 2011; Shikano and Cory, 2016; Bernardo and Singer, 2017).

The total population size for each host is *N*_*i*_ = *S*_*i*_ + *I*_*i*_. We restrict our analysis to scenarios where there is a stable endemic multi-host equilibrium where both hosts coexist with the pathogen, 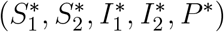, and a stable endemic single-host equilibrium where only the focal host and pathogen coexist, 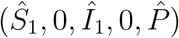. These conditions are satisfied in all of our numerical simulations, but they are not satisfied in all regions of parameter space (for example, when interspecific host competition or infection drives one host extinct).

### 2.2 High and low competence hosts, sinks, and sources

Throughout, we describe host species as having higher or lower competence and being sinks or sources of infectious propagules. Competence is the ability of an individual to transmit the pathogen to another susceptible individual. We measure host competence by 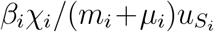, which simplifies to *p*_*i*_*χ*_*i*_*/*(*m*_*i*_ +*μ*_*i*_) after substituting 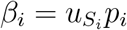. That quantity is the number of new infections produced by a single infected individual when placed in a completely susceptible population of infinite size; it is the basic reproductive number, *ℛ*_0_, when host density is arbitrarily large. One consequence of this definition is that host competence depends on the host’s environment through its mortality rate (*m*_*i*_). This means that it may be possible for a focal host to have higher competence than a second host in one environment and lower competence in another environment. Nonetheless, in any environment, higher competence hosts produce more new infections per infected individual because they have a combination of larger infection coefficients (*β*_*i*_), larger propagule release rates (*χ*_*i*_), smaller disease-induced mortality rates (*μ*_*i*_), and smaller uptake rates by susceptible individuals 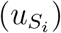.

Sink and source are defined by the per host net production of infectious propagules by infected individuals at the multi-species equilibrium 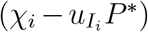. Source hosts excrete more infectious propagules than they take up 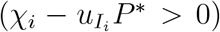 whereas sink hosts excrete less infectious propagules than they take up 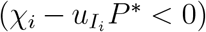. For any system, a host is a larger source (or a smaller sink) if it has a larger propagule release rate (*χ*_*i*_) and a smaller uptake rate 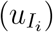. Note that because the definitions of sink and source depend on the equilibrium density of infectious propagules (*P*^*\**^), a host species can be a source in one environment (where infectious propagule density is low) and a sink in another (where infectious propagule density is high). Consequently, whether a host is a sink or source will depend on the characteristics of the pathogen (e.g., degradation rates, *δ*) and the characteristics of the other host species present in the community. For example, if the focal host has a higher excretion rate and a lower uptake rate than the second host, then the focal host will necessarily be a source and the second host will be a smaller source or a sink. In addition, it is possible for a host to switch from a sink to a source, or vice versa, when model parameters are varied over large ranges. We control for this in our numerical simulations by using parameter sets where the designation of sink and source is unchanged for all or nearly all of the range of parameter values; see specific figures for details.

Competence and sink/source are related characteristics but not identical. A high competence host can be a source if its excretion parameter (*χ*_*i*_) is large and its uptake parameters 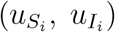 are small. Similarly, a low competence host can be a sink if its excretion (*χ*_*i*_) parameter is small and its uptake parameters 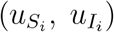 are large. In comparison, a high competence host can be a sink if its per propagule infection probability (*p*_*i*_) is close to one, its uptake rates 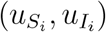 are large, and its mortality (*μ*_*i*_) and excretion (*χ*_*i*_) rates are small; this results in *p*_*i*_*χ*_*i*_*/*[*m*_*i*_ + *μ*_*i*_] being large and 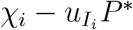 being negative. A low competence host can be a source if its per propagule infection probability (*p*_*i*_) and uptake rates 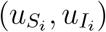 are small and its mortality (*μ*_*i*_) and excretion (*χ*_*i*_) rates are large; this results in *p*_*i*_*χ*_*i*_*/*[*m*_*i*_ +*μ*_*i*_] being small and 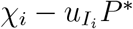 being positive.

### 2.3 Computing how prevalence depends on model parameters

Our metric of disease is the infection prevalence in the focal host at the multi-species equilibrium 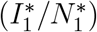. Infection prevalence depends on the characteristics of the second host that define competence (*χ*_2_, *β*_2_, *μ*_2_, and 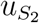), sink/source 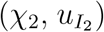, intraspecific competitive ability (e.g., *α*_22_ in a Lotka-Volterra model), and interspecific competitive ability (e.g., *α*_12_ in a Lotka-Volterra model). Our approach is to compute how infection prevalence of the focal host changes as the competence and competitive ability of the second host is varied. We compute this dependence based on how a small change in one parameter affects the infection prevalence of the focal host at the multi-species equilibrium. Mathematically, this is done using partial derivatives. For example, the effect of the second host having a higher infection coefficient (*β*_2_) is computed using the derivative 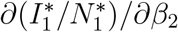; positive and negative values mean larger infection coefficients lead to higher or lower prevalence in the focal host, respectively.

The derivatives are computed using the Jacobian-based theory developed in Bender et al. (1984), Yodzis (1988), Novak et al. (2011), and Cortez and Abrams (2016); see appendix S1.2 for additional details. The Jacobian is a matrix of derivatives that accounts for all of the intraspecific and interspecific interactions of the system. The Jacobian for model (1) is

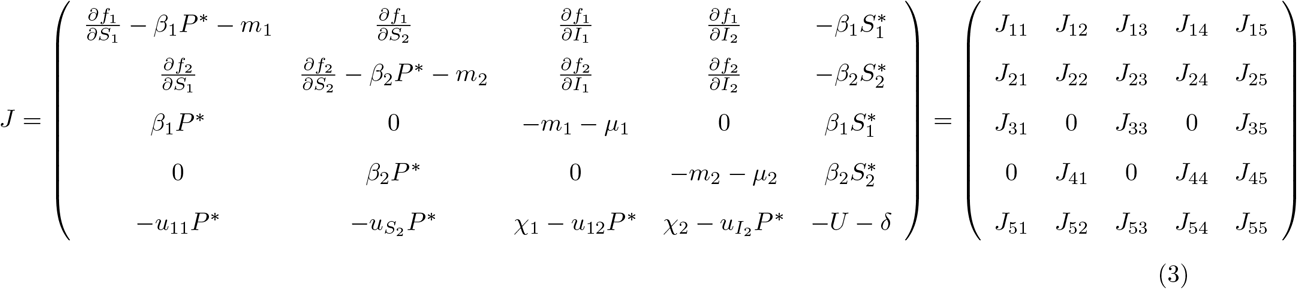

where 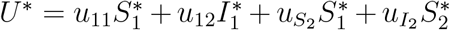 is the total per spore uptake rate at equilibrium. The entries in the first and second rows represent combinations of the effects of interspecific competition (*J*_12_, *J*_14_, *J*_21_, *J*_23_), infection (*J*_11_, *J*_15_, *J*_22_, *J*_25_), and intraspecific competition and reproduction (*J*_11_, *J*_13_, *J*_22_, *J*_24_); the entries in the third and fourth rows represent the effects of infection (*J*_31_, *J*_35_, *J*_42_, *J*_45_) and mortality (*J*_33_, *J*_44_); and the entries in the fifth row represent the negative effects due to uptake by susceptible individuals (*J*_51_, *J*_52_) and degradation (*J*_55_) and the combined effects of propagule release and uptake by infected individuals (*J*_53_, *J*_54_).

In total, our method allows us to write each derivative in terms of the Jacobian entries. (Due to their large size, all derivative equations are relegated to the appendices.) Because each Jacobian entry corresponds to a specific set of ecological processes, we can disentangle how each characteristic of the second host affects infection prevalence in the focal host and whether there are interactions between the characteristics (e.g., the effect of increased competence in the second host may depend on its interspecific competitive ability). This allows us to make predictions about the factors that promote higher or lower infection prevalence in the focal host. This in turn yields insight into the factors that promote amplification (i.e., higher prevalence in the focal host at the multi-species equilibrium than at the single-species equilibrium; 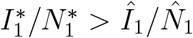) versus dilution (i.e., lower prevalence in the focal host at the multi-species equilibrium than the at single-species equilibrium; 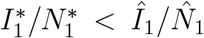), respectively.

One limitation of our approach is it is based on a linear approximation (the Jacobian). It is guaranteed to accurately predict how small changes in parameters affect infection prevalence, but it may not accurately capture the effects of large changes in parameters. Nonetheless, our approach only depends on the signs of the Jacobian entries, most of which are constant for all parameter values. For example, the signs of all entries in column 5 and all entries in rows 3 and 4 of the Jacobian do not change. Because of this, we can predict when the relationship between infection prevalence and a parameter will change signs (e.g., switch from increasing to decreasing), which in turn helps explain patterns observed when there are large changes in parameter values.

3 Results

In the following, we first show how our environmental transmission model can be unified with frequency-dependent and density-dependent direct transmission models. We then explore how the transmission mechanism and the competence and competitive ability of a second host affect infection prevalence in the focal host. In each section, we compare our results to predictions from previous theoretical studies in order to show how our results can help explain why differing predictions have been made.

### 3.1 Unifying environmental and direct transmission models

We first show that our environmental transmission model (1) turns into a density-dependent direct transmission (DDDT) model or frequency-dependent direct transmission (FDDT) model when the parameter values satisfy particular conditions. Importantly, this means that predictions made using our environmental transmission model apply to environmental transmission (ET), FDDT, and DDDT pathogens. Thus, our results are a first step towards unifying the dilution effect theory that has developed independently for direct transmission pathogens and environmental transmission pathogens. Our work builds on Lafferty et al. (2015), which showed that ET, DDDT, and FDDT models are all special cases of a general-consumer resource model. Following previous studies (Li et al., 2009; Eisenberg et al., 2013; Cortez and Weitz, 2013), we use an alternative approach wherein we identify specific conditions under which our ET model reduces to a DDDT or FDDT model.

In general, the ET model reduces to a model with direct transmission when the host excretion rates (*χ*_*i*_) are large and the infectious propagule uptake rates 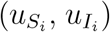 or degradation rate (*δ*) are large. If the loss of infectious propagules due to uptake by hosts is negligible compared to loss due to degradation (*U* + *δ* ≈ *δ*), then the ET model reduces to a DDDT model. Alternatively, if there is no degradation of infectious propagules (*δ* = 0), then the ET model reduces to a FDDT model. The intuition is the following. Infectious propagules persist in the environment for short periods of time when degradation or uptake rates are large. Consequently, susceptible individuals can only encounter infectious propagules immediately after the infectious propagules are excreted by an infected individual. This requires susceptible and infected individuals to be in close proximity, in effect implying infection only occurs when there are direct contacts between infected and susceptible individuals. When loss due to uptake is negligible compared to degradation (*U* + *δ* ≈ *δ*), contact rates between susceptible individuals and infectious propagules are proportional to the density of infected individuals. In this case, the dynamics of the ET pathogen are essentially identical to those of a DDDT pathogen. In contrast, when there is no degradation (*δ* = 0), contact rates between susceptible individuals and infectious propagules are proportional to the weighted frequency of infected individuals in the community, where the weights are the uptake rates of each host class. In this case, the dynamics of the ET pathogen are essentially identical to those of a FDDT pathogen.

To mathematically justify the above, we assume the changes in infectious propagule density are much faster than changes in the host densities, i.e., the host excretion rates (*χ*_*i*_) and infectious propagule uptake 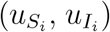 or degradation (*δ*) rates are large. Under this condition, the infectious propagule densities reach a quasi-steady state defined by *dP/dt* = 0; see appendix S1.3 for details. Solving for the quasi-steady state density and substituting into the infected host equations yields

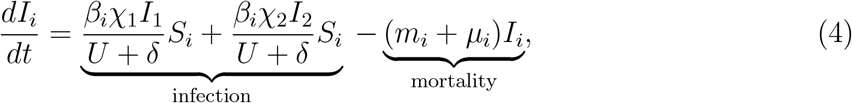

where 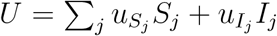 is the total uptake of infectious propagules by all host classes. When loss due to uptake is negligible relative to degradation (*U* + *δ* ≈ *δ*), the infection rates simplify to the infection rates of a DDDT model, 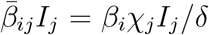. When there is no degradation (*δ* = 0), the infection rates simplify to the infection rates of a FDDT model, 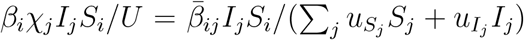, where the weights in the denominator determine if an infected individual is more likely to have contacts with individuals in the focal host population or the second host population. For example, 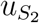 and 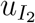 larger than 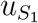 and 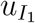 means an infected individual is more likely to have contacts with individuals in the second host population than the focal host population.

### 3.2 Interpreting the relationship between environmental and direct transmission models

There are four important consequences of the mathematical relationship between the ET, DDDT, and FDDT models. First, our results show that environmental transmission lies intermediate between the two different types of direct transmission. In particular, the three transmission modes sit in a two-dimensional space defined by the total uptake rate (*U*) and degradation rate (*δ*), with ET lying intermediate between DDDT and FDDT (see Figure 1A). The ET model reduces to a FDDT model when the degradation rate is zero (*δ* = 0; blue vertical axis) and it reduces to a DDDT model when the uptake rates are negligible (*U* = 0; red horizontal axis). When the uptake and degradation rates are both non-negligible (*δ* > 0, *U* > 0; white), ET behaves like a combination of density-dependent and frequency-dependent direct transmission.

**Figure 1:**
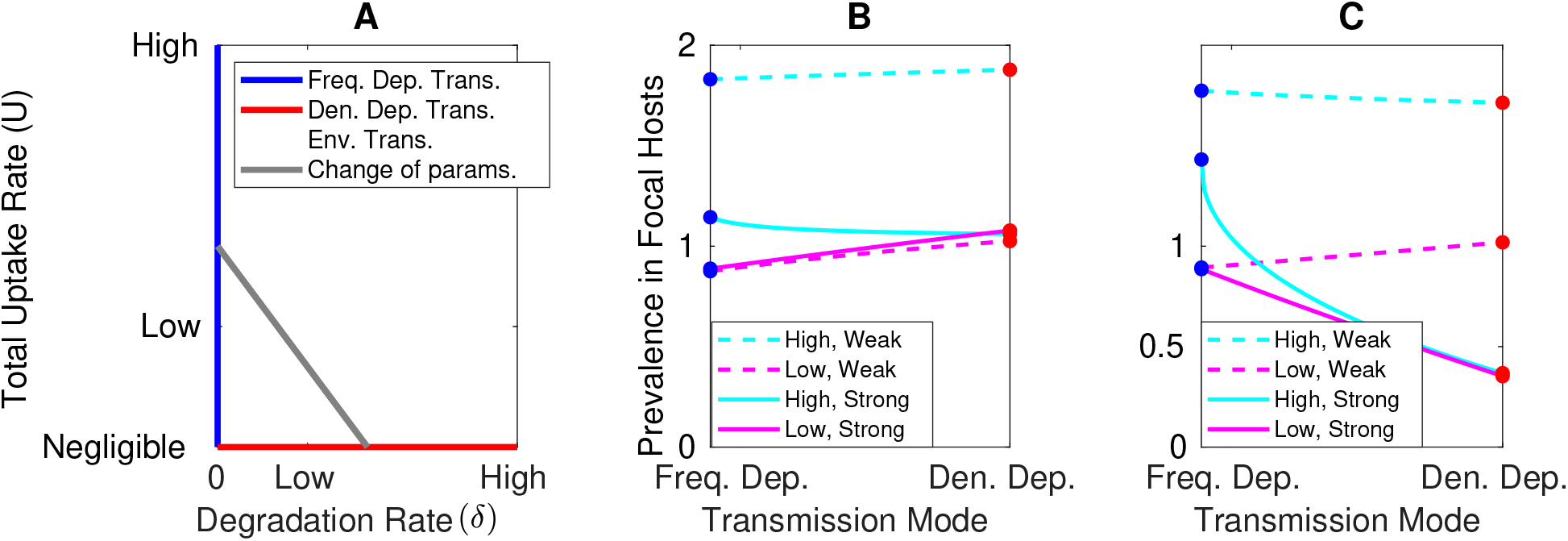
Environmental transmission models and density-dependent and frequencydependent direct transmission models can be unified, which helps identify how the transmission mechanism influences infection prevalence in a focal host. (A) Environmental transmission sits intermediate between density-dependent and frequency-dependent direct transmission. Environmental transmission models are defined by all possible uptake and degradation rates (blue, red and white) and they are identical to density-dependent direct transmission models when loss of infectious propagules due to uptake by hosts is negligible (*U* = 0, red) and identical to frequency-dependent direct transmission models when there is no infectious propagule degradation (*δ* = 0, blue). Gray line illustrates a change of parameters that transforms a particular parameterization of the environmental transmission model from a frequency-dependent form to a density-dependent form while holding the single-species equilibrium densities constant; see text for details. (B,C) Effect of transmission mode on focal host prevalence in the (B) absence and (C) presence of interspecific host competition when the second host has low (magenta lines) or high (cyan lines) competence and is a weak (dashed lines) or strong (solid lines) intraspecific competitor. The vertical axis is the ratio of the prevalence in the focal host at the multi-species equilibrium to that at the single-species equilibrium, 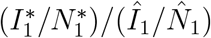; values greater or less than 1 mean that prevalence in the focal host increases or decreases, respectively, with the addition of the second host. Each curve shows the ratio as the environmental transmission model is transformed from a frequency-dependent form (blue dots) to a density-dependent form (red dots) while holding the single-species equilibrium densities constant. See appendix S4 for models and parameters.

Second, when loss due to both uptake and degradation are non-negligible and the infectious propagule dynamics are fast, the ET model acts like a direct transmission model where transmission is intermediate to DDDT and FDDT. Specifically, the direct transmission rate *β*_*i*_*χ*_*j*_*I*_*j*_*/*(*U* + *δ*) in equation (4) looks like DDDT when infected density is low and like FDDT when infected density is high. This qualitatively matches transmission rates estimated for empirical systems (Smith et al., 2009; Hopkins et al., 2020). In addition, while the assumption of fast infectious propagule dynamics was necessary for the dynamics of the environmental and direct transmission models to be identical, our results about equilibrium infection prevalence apply for any speed of the infectious propagule dynamics. This is because equilibria of the ET model are defined by *dI*_*i*_*/dt* = 0 and *dP/dt* = 0, which is identical to equation (4) being equal to zero for the direct transmission models. Consequently, our results about infection prevalence at equilibrium for ET pathogens apply to pathogens with DDDT, FDDT, or transmission rates intermediate to DDDT and FDDT.

Third, changes in the parameter values of the ET model can be interpreted in terms of changes in the parameter values of a DDDT or FDDT model. Varying the excretion rate (*χ*_*i*_) or per propagule infection probability (*p*_*i*_) in the ET models translates to varying the direct transmission rate parameters of a DDDT or FDDT model, e.g., 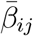 in the rates 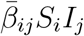 or 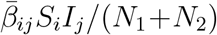. Varying an uptake parameter 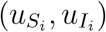 translates to increasing the weights in the direct transmission rate of a FDDT model, e.g., increasing *w*_*i*_ in 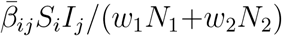; there is no corresponding change in DDDT models because *U* = 0. Changes in the other epidemiological parameters (like disease-induced mortality) are identical across models. In total, this means that higher or lower competence of a host in the ET model implies higher or lower competence of a host in a DDDT or FDDT model, and vice versa.

Fourth, the concepts of sink and source host extend to direct transmission pathogens. For DDDT, all hosts are sources. This is because DDDT corresponds to negligible uptake in the ET model, which means the per individual net production rate of infectious propagules is positive 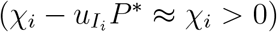. For FDDT, a host can be a sink or a source. Recall that the second host is a sink in the ET model when *χ*_2_ is small and 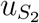 and 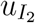 are large. When translated to a FDDT model, the second host is a sink when the interspecific transmission coefficient 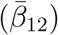 is smaller and infected individuals have an equal or larger likelihood of having contacts with focal host individuals than second host individuals (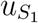 and 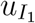 larger than 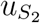 and 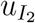). For example, previous modeling studies (Rudolf and Antonovics, 2005; Dobson, 2004; Mihaljevic et al., 2014; Roberts and Heesterbeek, 2018) use the transmission rates 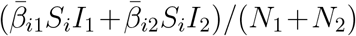, where all weights are unity 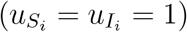 and the interspecific transmission parameters are smaller than the intraspecific transmission parameters 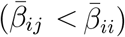. Under these conditions, the second host is always a sink.

### 3.3 The effects of the pathogen transmission mechanism on infection prevalence

We now identify how the pathogen transmission mode influences prevalence in the focal host. Our goal is to identify conditions under which infection prevalence in the focal host is higher for FDDT pathogens than DDDT pathogens, and vice versa.

Our approach is to use a change of parameters to convert the ET model from a form that behaves like a DDDT model (*U* = 0) to a form that behaves like a FDDT model (*δ* = 0) (gray line in Figure 1A). The general approach for creating changes of parameters is discussed in appendix S1.4.1. To make a fair comparison between models, our change of parameters holds all parameters constant except the uptake 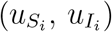 and degradation (*δ*) rates, which must necessarily differ between the models. The uptake and degradation rates are varied such that we hold constant the per capita total loss rate of infectious propagules at the single-species equilibrium 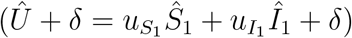. This particular change of parameters results in the single-species equilibrium densities and infection prevalence being the same across models and only the multi-species equilibrium densities differing across models. This allows us to isolate how the transmission mechanism influences disease prevalence in the focal host. The biological way to think about the transformation is that the endemic prevalence in the focal host at the single-species equilibrium is known, the pathogen transmission mechanism is unknown, and we want to predict how the effect of the addition of the second host on prevalence in the focal host depends on the transmission mechanism. Our results are unchanged if we use a change of parameters that instead holds the multi-species equilibrium densities constant and causes the single-species equilibrium densities to change; see appendix S1.4.3 for details. The biological way to think about this transformation is that the endemic prevalence in the focal host at the multi-species equilibrium is known, the pathogen transmission mechanism is unknown, and we want to predict how the effect of removal of the second host depends on the transmission mechanism.

Prevalence in the focal host under ET always sits intermediate to the prevalence under FDDT and DDDT. Whether prevalence in the focal host is higher or lower under FDDT than DDDT depends on the levels of intraspecific and interspecific host competition and the competence of the second host; see appendix S1.4.4 for details. Specifically, infection prevalence in the focal host is lower under FDDT than DDDT when (i) interspecific host competition is weak, (ii) the second host is a weak intraspecific competitor, and (iii) the second host has low competence. In comparison, infection prevalence in the focal host is lower under DDDT than FDDT when (i) interspecific host competition is strong, (ii) the second host is a strong intraspecific competitor, and (iii) the second host has high competence.

Figure 1 shows numerical examples of how the transmission mechanism affects prevalence in the focal host. In the absence of interspecific competition (Figure 1B), prevalence in the focal host is typically lower under FDDT than DDDT (blue dots are lower than red dots for solid magenta, dashed magenta, and dashed cyan curves). The opposite only occurs if the second host is a strong intraspecific competitor and a high competence host (blue dot higher than red dot for solid cyan curve). When there is interspecific competition between hosts (Figure 1C), the relationship between the transmission mode and prevalence in the focal host can reverse. For example, in the absence of interspecific competition (Figure 1B), focal host prevalence is lower under FDDT when the second host is a low competence, strong intraspecific competitor (solid magenta “Low, Strong” curve) or a high competence, weak intraspecific competitor (dashed cyan “High, Weak” curve). However, the pattern reverses when interspecific competition is sufficiently strong (solid magenta and dashed cyan curves are decreasing in Figure 1C). In our numerical simulations, transmission mode only had a modest effect on focal host prevalence in all cases where the second host was a high competence, weak intraspecific competitor and increased interspecific competition reversed the relationship between transmission mode and focal host prevalence (dashed cyan curves in Figure 1B,C have slopes of small magnitude).

In addition to predicting when one transmission mechanism leads to higher or lower prevalence in the focal host, our results identify when dilution will be more common under FDDT than DDDT, and vice versa. We predict that dilution is more common under FDDT than DDDT, unless interspecific host competition is sufficiently strong. We outline our reasoning here; see appendix S1.4.4 for details. For any level of interspecific competition, it is possible for dilution to occur under FDDT and amplification to occur under DDDT. For example, the dashed and solid magenta curves in Figure 1B and the dashed magenta curve in Figure 1C show dilution for FDDT (blue dots below 1) and amplification for DDDT (red dots above 1). In contrast, only when interspecific competition is sufficiently strong can dilution occur under DDDT and amplification occur under FDDT (blue dot above 1 and red dot below 1 for solid cyan curve in Figure 1C). Overall, this means that dilution will be more common for FDDT pathogens than DDDT pathogens, unless there is sufficiently strong interspecific host competition.

Many previous studies predict dilution occurs more under FDDT than DDDT because levels of disease often increase with biodiversity under DDDT (Begon et al., 1992; Dobson, 2004; Faust et al., 2017) and often decrease with biodiversity under FDDT (Rudolf and Antonovics, 2005; Dobson, 2004; Mihaljevic et al., 2014; Roberts and Heesterbeek, 2018). Our results suggest that this is because most previous studies assume all hosts are competent (i.e., can spread the disease) and there is weak or no interspecific competition between host species. Under these conditions, we predict prevalence in the focal host will be lowest under FDDT. However, if interspecific host competition is sufficiently strong or the second host has high competence, then prevalence in the focal host can be highest under FDDT.

### 3.4 The effects of host competence on infection prevalence

Increased competence of the second host causes higher infection prevalence in the focal host, unless the second host is a sufficiently large sink. Intuition suggests that higher competence in the second host (larger 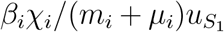) willcause greater prevalence i n the focal host. This pattern holds under many conditions. For example, increased excretion (larger *χ*_2_) or decreased uptake (smaller 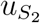 and 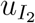) leads to greater infection prevalence in the focal host. Prevalence in the focal host can also increase with decreased mortality (*μ*_2_) of the second host (red, magenta, and cyan curves in Figure 2A) and increases in the infection coefficient (*β*_2_) for the second host (left side of all curves in Figure 2B).

**Figure 2:**
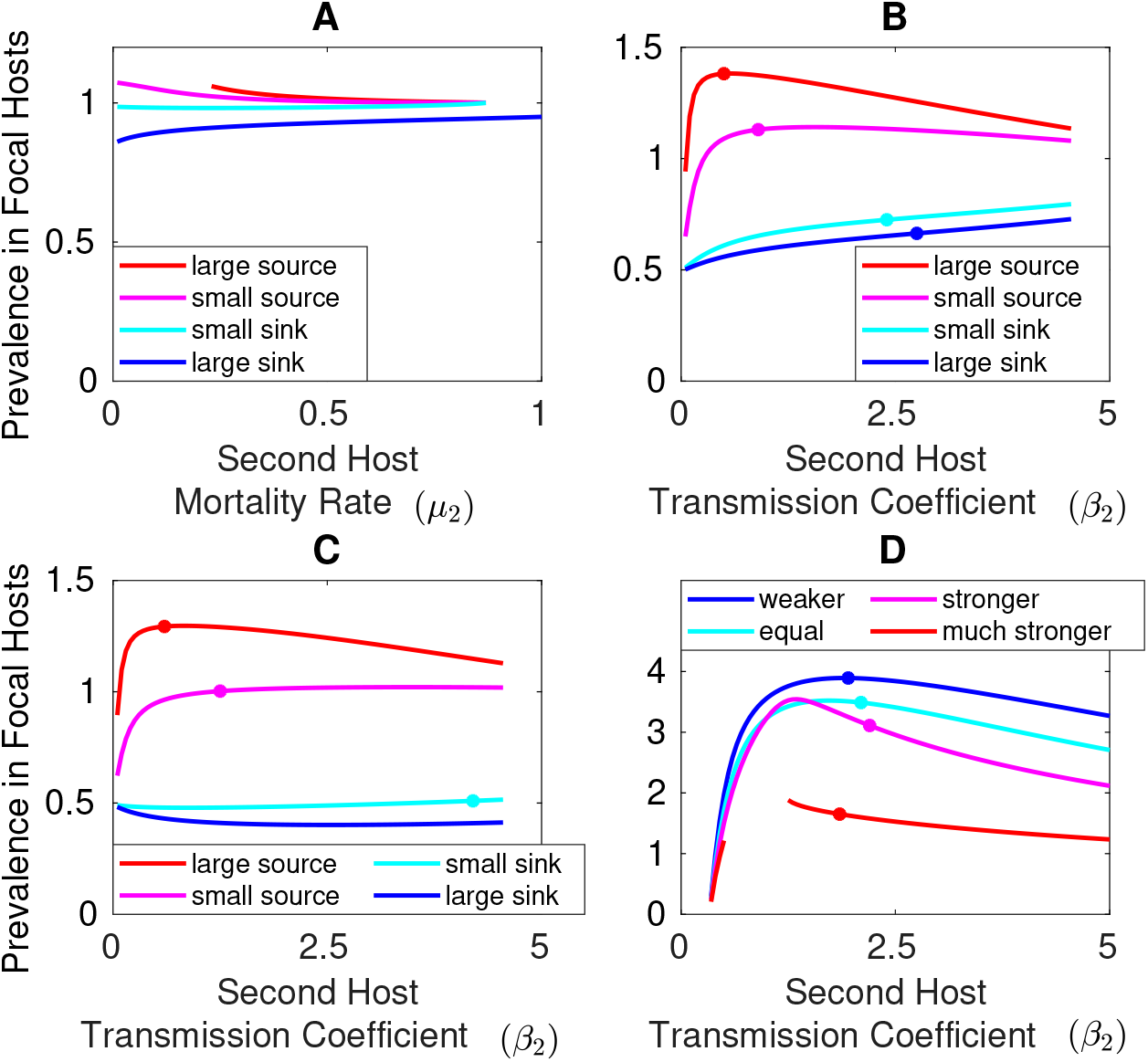
Increased competence of the second host leads to greater infection prevalence in a focal host, unless the second host is a sufficiently large sink. This prediction can be reversed if one or both hosts experience strong positive density dependence at equilibrium or infected individuals are stronger interspecific competitors than susceptible individuals. For all panels the vertical axis is the ratio of the prevalence in the focal host at the multi-species equilibrium to that at the single-species equilibrium, 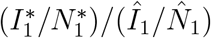; values greater or less than 1 mean that prevalence in the focal host increases or decreases, respectively, with the addition of the second host. All panels show how the ratio changes as components defining the competence of the second host are varied. Filled circles in panels B-D denote parameter values above which at least one host experiences positive density dependence. (A) Responses to increased disease-induced mortality in the second host when the second host is a (blue) large sink, *χ*_2_ = 0.2, (cyan) small sink, *χ*_2_ = 1, (magenta) small source, *χ*_2_ = 2, or (red) large source, *χ*_2_ = 6. (B,C) Responses to increases in the transmission rate of the second host when infected individuals of the second host have (B) equal 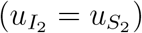 or (C) greater 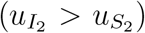 uptake rates than susceptible individuals and the second host is a (blue) large sink, *χ*_2_ = 0.1, (cyan) small sink, *χ*_2_ = 0.2, (magenta) small source, *χ*_2_ = 2, or (red) large source, *χ*_2_ = 6. (D) Responses to increases in the transmission rate of the second host when infected individuals are (blue) weaker, (cyan) equal, (magenta) stronger, or (red) much stronger interspecific competitors than susceptible individuals; the break in the red curve is due to coexistence being impossible for intermediate transmission coefficients. The designations of sink and source apply for all parameter values used to generate the figures; see appendix S4 for equations and parameters.

However, higher competence in the second host can decrease prevalence in the focal host if the second host is a sufficiently large sink. When increased competence of the second host is due to decreases in its disease-induced mortality rate (*μ*_2_), focal host disease prevalence can increase with increased disease-induced mortality of the second host if the second host is a sufficiently large sink (increasing blue curve in Figure 2A). When increased competence of the second host is due to increases in its transmission coefficient (*β*_2_), focal host infection prevalence can decrease with increases in the transmission coefficient of the second host, but only if the second host is a sufficiently large sink and infected individuals of the second host have higher uptake rates than susceptible individuals 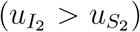. For example, when the second host is a sink, increases in the transmission coefficient of the second host cause focal host infection prevalence to increase when infected and susceptible individuals of the second host have equal uptake rates (left side of blue curve is increasing in Figure 2B) and decrease when infected individuals of the second host have sufficiently greater uptake rates than susceptible individuals (left side of blue curve is decreasing in Figure 2C).

The reason for the differing responses is that there are multiple indirect pathways through which changes in the competence of the second host propagate through the system and affect prevalence in the focal host. To provide intuition, we focus on a few particular indirect pathways and refer the reader to appendices S1.6.1-S1.6.3 for additional details about other pathways. In Figure 3, we represent the indirect pathway as a series of arrows between variables in the system and as a chain of partial derivatives where dots denote derivatives with respect to time (e.g., *İ* _1_ = *dI*_1_*/dt*).

**Figure 3:**
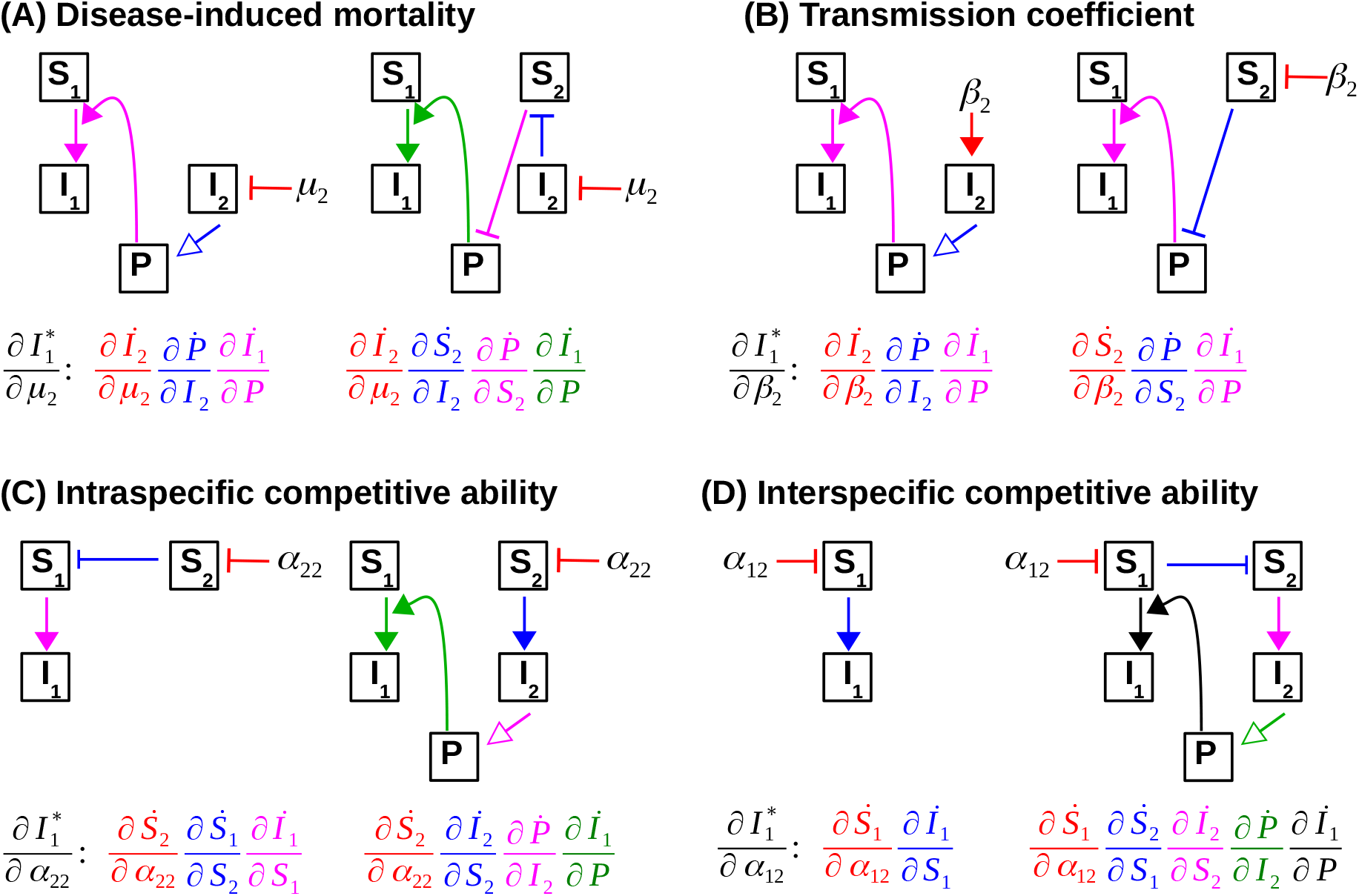
Illustration of some indirect pathways through which the competence and competitive ability of the second host affect infection prevalence in the focal host. Flat arrows and filled pointed arrows denote direct effects that are negative and positive, respectively. Open pointed arrows denote direct effects that are negative or positive depending on whether the second host is a sink or source, respectively. The color of each arrow matches the color of the corresponding term in the sequence of direct effects. See main text for a more complete explanation of the pathways. (A) Two indirect pathways affecting the response to an increase in the disease-induced mortality of the second host. The right pathway is negative and the left pathway is positive or negative if the second host is a sink or source, respectively. (B) Two indirect pathways affecting the response to an increase in the transmission coefficient of the second host. The right pathway is positive and the left pathway is positive or negative if the second host is a source or sink, respectively. (C) Two indirect pathways affecting the response to an increase in the intraspecific competitive ability of the second host. The left pathway is positive and the right pathway is negative or positive if the second host is a source or sink, respectively. (D) Two indirect pathways affecting the response to an increase in the interspecific competitive ability of the second host. The left pathway is negative and the right pathway is negative or positive if the second host is a sink or source, respectively.

The reason why increases in the disease-induced mortality of the second host (*μ*_2_) typically decrease focal host infection prevalence is the following. Increasing the disease-induced mortality of the second host causes a decrease in its infected density (red flat arrows in Figure 3A, *∂İ*_2_*/∂μ*_2_ < 0). One consequence of reduced infected density in the second host (left side of Figure 3A) is a change in the production of infectious propagules by infected individuals. If the second host is a source, this decreases the density of infectious propagules (blue arrow, 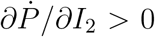), which leads to lower prevalence in the focal host (magenta arrow, *∂İ*_1_*/∂P* > 0). If the second host is a sink, then infectious propagule density increases (blue arrow, 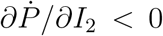), which leads to greater prevalence in the focal host (magenta arrow, *∂İ*_1_*/∂P* > 0). A second consequence of reduced infected density in the second host (right side of Figure 3A) is reduced intraspecific competition, which causes an increase in the susceptible density of the second host (blue flat arrow, 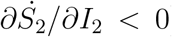). The increase in susceptible density lowers infectious propagule density (purple flat arrow, 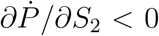), which leads to lower infection prevalence in the focal host (purple flat arrow, *∂İ*_1_*/∂P* < 0). Because the right pathway is always negative, the net effect of the two pathways is negative unless the second host is a sufficiently large sink. Overall, increases in the disease-induced mortality rate (*β*_2_) cause decreased disease prevalence in the focal host, unless the second host is a sufficiently large sink.

In contrast, increases in the transmission parameter of the second host (*β*_2_) typically increase focal host infection prevalence for the following reason. Increases in the transmission parameter of the second host cause an increase in its infected density and a decrease in its susceptible density, both of which affect infectious propagule density (left and right sides of Figure 3B, respectively). If infected individuals have equal or lower uptake rates than susceptible individuals, the net effect of the two pathways is an increase in infectious propagule density because susceptible individuals who only take up infectious propagules become infected individuals who both excrete infectious propagules and take them up at equal or lower rates. The net increase in infectious propagules results in greater infection prevalence in the focal host. In contrast, if infected individuals have greater uptake rates than susceptible individuals and infected individuals are sinks, then the net effect of the two pathways is a decrease in infectious propagule density because an individual’s uptake rate increases when it becomes infected. The net decrease in infectious propagules results in lower infection prevalence in the focal host. Overall, increases in the transmission coefficient (*β*_2_) cause increased disease prevalence in the focal host, unless the second host is a sufficiently large sink and infected individuals of the second host have larger uptake rates then susceptible individuals of the second host.

Our results can help explain why there are contrasting predictions about how host competence affects amplification and dilution for DDDT and FDDT pathogens. Many theoretical studies predict that higher competence hosts amplify DDDT pathogens (Begon et al., 1992; Joseph et al., 2013; O’Regan et al., 2015; Roberts and Heesterbeek, 2018). We predict this occurs because all hosts are sources under DDDT. In contrast, other studies predict higher competence hosts can dilute FDDT pathogens (Rudolf and Antonovics, 2005; O’Regan et al., 2015). In those studies, the transmission rates are of the form 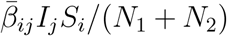, where the weights for all individuals are unity and interspecific rates of transmission are less than intraspecific rates of transmission 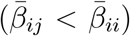. As explained in “Interpreting the relationship between environmental and direct transmission models”, these conditions correspond to the second host having a low *χ*_2_ value and being a sink, which we predict decreases prevalence in the focal host.

### 3.5 The effects of intraspecific and interspecific host competition on infection prevalence

#### Intraspecific competitive ability of the second host

Stronger intraspecific competition in the second host leads to increased focal host prevalence, unless the second host is a sufficiently large source (e.g., has very high excretion rates); see appendix S1.6.4 for details. In addition, the threshold for being a sufficiently large source increases with increased interspecific competition between the hosts. For example, in the absence of interspecific competition (Figure 4A), stronger intraspecific competition leads to greater prevalence in the focal host when the second host is a sink (blue curve) and lower prevalence when the second host is a source (cyan and red curves). However, when interspecific competition is higher (Figure 4B), stronger intraspecific competition causes lower prevalence only if the second host is a sufficiently large source (cyan curve switches from decreasing in Figure 4A to increasing in Figure 4B).

**Figure 4:**
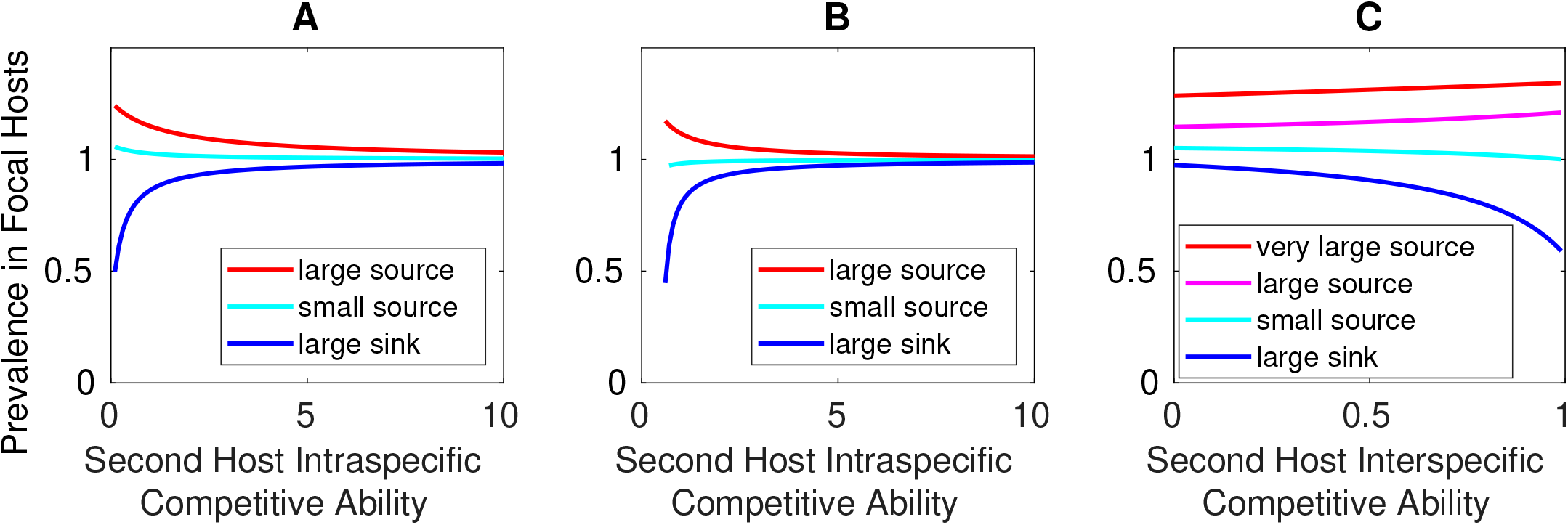
Increased intraspecific competitive ability of the second host causes greater infection prevalence in the focal host and increased interspecific competitive ability of the second host leads causes infection prevalence in a focal host, unless the second host is a sufficiently large source of infectious propagules. For all panels the vertical axis is the ratio of the prevalence in the focal host at the multi-species equilibrium to that at the single-species equilibrium, 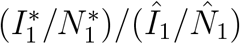; values greater or less than 1 mean that prevalence in the focal host increases or decreases, respectively, with the addition of the second host. Responses to increased intraspecific competitive ability of the second host in the (A) absence and (B) presence of interspecific competition when the second host is a (blue) large sink, *χ*_2_ = 0.5, (cyan) small source, *χ*_2_ = 1.5, or (red) large source, *χ*_2_ = 3. Note that all sink hosts become small sources when the intraspecific competitive ability of the second host is sufficiently low; see text for details. (C) Responses to increased interspecific competitive ability of the second host when the second host that is a (blue) large sink, *χ*_2_ = 1, (cyan) small source, *χ*_2_ = 2, (magenta) large source, *χ*_2_ = 4, or (red) very large source, *χ*_2_ = 12. See appendix S4 for equations and parameters.

The reason is that increased intraspecific competitive ability of the second host has two effects on the focal host transmission rate. First, increasing the intraspecific competitive ability (*α*_22_) of the second host causes a decrease in its density of susceptible individuals (red flat arrow in the left side of Figure 3C, 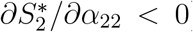). This reduces competition with the focal host, which causes an increase in susceptible focal host density (flat blue arrow, 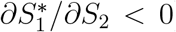). The resulting increase in the focal host transmission rate leads to higher prevalence in the focal host (purple arrow, 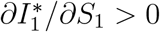). Note that the effect of this indirect pathway is stronger when interspecific competition is stronger.

Second, the decrease in susceptible density in the second host population (red flat arrow in the right side of Figure 3C, 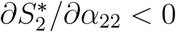) also reduces the transmission rate of the second host, resulting in a decrease in infected density in the second host (blue arrow, 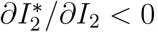). If the second host is a source, then the decrease in infected density leads to lower infectious propagule density (magenta arrow, *∂P*\*/*∂I*_2_ > 0), which results in lower prevalence in the focal host (green arrow, 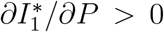). On the other hand, if the second host is a sink, then the decrease in infected density in the second host leads to greater infectious propagule density (magenta arrow, *∂P*\*/*∂I*_2_ > 0), which results in greater prevalence in the focal host (green arrow, 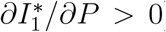). The combined effect of these two pathways is that stronger intraspecific competition in the second host leads to increased prevalence in the focal host, unless the second host is a sufficiently large source. In addition, because the effects of the first pathway increase with interspecific competition, the threshold for being a sufficiently large source increases with increased interspecific competition between the hosts.

We note that in Figure 4A&B the second host switches from a sink to a small source as the intraspecific competitive ability of the second host is decreased to a sufficiently small values (blue curves, far left side). This is a necessary outcome: Decreasing intraspecific competition in the second host causes its density to increase, which because the second host is sink, causes the infectious propagule density to decrease. Eventually infectious propagule density becomes so low that 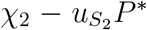 is positive and small, which implies host 2 is a small source. Consequently, the designation of sink holds for most, but not all, of the parameter range in Figure 4A&B. Nonetheless, the switch from sink to small source does not qualitatively affect our general prediction because the relationship between focal host infection prevalence and intraspecific competition in the second host is positive so long as the second host is not a large source.

#### Interspecific competitive ability of the second host

Stronger interspecific competitive ability of the second host causes a decrease in focal host prevalence (blue and cyan curves in Figure 4C), unless the second host is a large source (magenta and red curves in Figure 4C); see appendix S1.6.5 for details.

The reason is that increased interspecific competitive ability of the second host has two effects on the focal host transmission rate. First, increased interspecific competitive ability of the focal host (larger *α*_12_ in a Lotka-Volterra model) causes a decrease in the density of susceptible focal hosts (red flat arrow in the left side of Figure 3D, 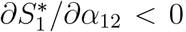). This decreases the focal host transmission rate and leads to lower prevalence in the focal host (blue arrow, 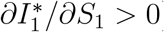).

Second, the decrease in the density of susceptible focal hosts (red flat arrow in the right side of Figure 3D, 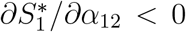) reduces competition with the focal host, which results in an increase in susceptible individuals in the second host population (flat blue arrow, 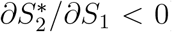). This leads to an increase in infected density in the second host population (green arrow, 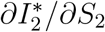). If the second host is a source, then the increase in infected density leads to greater infectious propagule density (magenta arrow, *∂P* ^*\**^*/∂I*_2_ > 0) and prevalence in the focal host (green arrow, 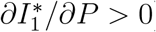). On the other hand, if the second host is a sink, then the increase in infected density leads to lower infectious propagule density (magenta arrow, 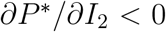) and lower prevalence in the focal host (green arrow, 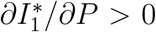). If the second host is a sufficiently large source, then the second indirect effect will be larger in magnitude than the first, resulting in stronger interspecific competition causing greater prevalence in the focal host.

Previous empirical studies have observed that interspecific competition can lead to reduced infection prevalence (Johnson et al., 2008), even when increased host diversity increases per individual transmission rates (Luis et al., 2018). This is supported by theoretical studies (Rudolf and Antonovics, 2005; Strauss et al., 2015) showing that increased interspecific competition can reduce disease prevalence. Our results suggest we should expect infection prevalence in a focal host to decrease with increased interspecific competitive ability of the second host, and that this is simply a consequence of decreased transmission due to reduced density of the focal host. However, our results also show that this pattern is not universal and may break down if the second host is a large source. For ET pathogens this means the second host has high excretion rates and for direct transmission pathogens this means the second host has high intraspecific and interspecific transmission coefficients.

## 3.6 Predictions for factors promoting amplification versus dilution

Our results focus on how prevalence in a focal host depends on the pathogen transmission mechanism and the characteristics of a second host. While this does not directly address the effects of biodiversity on disease, it does provide some insight about factors promoting amplification versus dilution in the focal host. Our predictions are summarized in Table 1.

**Table 1:** Predictions for when the pathogen transmission mode and characteristics of a second host promote amplification or dilution in a focal host

| Characteristic | Predicted effects on prevalence in a focal host |
| --- | --- |
| Transmission mode | Frequency-dependent direct transmission promotes dilution more than density-dependent direct transmission when<br>(i) interspecific host competition is weak<br>(ii) second host is a weaker intraspecific competitor<br>(iii) second host is a lower competence host<br>Density-dependent direct transmission promotes dilution more than frequency-dependent direct transmission when<br>(i) interspecific host competition is strong<br>(ii) second host is a stronger intraspecific competitor<br>(iii) second host is a higher competence host |
| Competence of second host | Higher competence promotes amplification, unless the second host is a sufficiently large sink |
| Intraspecific competitive ability of second host | Stronger intraspecific competition promotes amplification, unless the second host is a sufficiently large source of infectious propagules |
| Interspecific competitive ability of second host | Weaker interspecific competition promotes amplification, unless the second host is a sufficiently large source of infectious propagules |
| Conditions that can reverse predictions | Sufficiently strong positive density dependence in either host<br>Interspecific competition is greater than intraspecific competition<br>Infected individuals are stronger interspecific competitors than susceptible individuals |

We predict (i) higher competence of the second host promotes amplification, unless the second host is a large sink (Figure 2); (ii) stronger intraspecific competitive ability of the second host promotes amplification, unless the second host is a large source (Figure 4A,B); and (iii) stronger interspecific competitive ability of the second host promote dilution, unless the second host is a large source (Figure 4C). We note that the reversal in prediction (i) only applies when higher competence of the second host means a higher transmission coefficient and/or a lower disease-induced mortality rate. We predict that in general dilution will occur more often under frequency-dependent direct transmission (FDDT) than density-dependent direct transmission (DDDT), unless interspecific host competition is strong. Moreover, we predict that introduction of a second host will dilute disease more (or amplify disease less) under FDDT than DDDT when (i) interspecific host competition is weak, (ii) the second host has lower competence, and (iii) the second host experiences weaker intraspecific competition. There are three conditions under which some or all of our predictions can be reversed.

First, every prediction can be reversed if interspecific host competition is greater than intraspecific host competition. This can occur, for example, in systems where the hosts cannot coexist in the absence of the pathogen and host coexistence is pathogen-mediated. Second, every prediction can be reversed if one or both hosts are experiencing positive density dependence (at equilibrium). This occurs when the pathogen reduces the density of one host to the point where the growth rate of that host is an increasing function of its own density. This is analogous to positive density dependence of a prey species in a predator-prey system, which occurs when the predator reduces the prey density to levels below the hump in the predator nullcline. The filled circles in Figure 2B-D denote the minimum parameter values at which one host is experiencing positive density dependence. When the positive density dependence is sufficiently large, the curves reverse direction. Third, the predictions about host competence can be reversed if infected individuals are sufficiently stronger interspecific competitors than susceptible individuals. For example, if infected individuals are stronger interspecific competitors, then higher competence hosts can amplify disease less (decreasing portions of magenta and red curves left of the filled circles in Figure 2D). This can occur in systems where infection causes hosts to have increased appetite or resource acquisition rates (Ponton et al., 2011; Shikano and Cory, 2016; Bernardo and Singer, 2017).

## 4 Discussion

Whether increased host biodiversity leads to greater or less disease has been contested in the literature (Lafferty and Wood, 2013; Ostfeld and Keesing, 2013; Wood and Lafferty, 2013), leading to calls for theory explaining when particular mechanisms promote amplification versus dilution (Buhnerkempe et al., 2015; Halsey, 2019; Rohr et al., 2020). We used a host-pathogen model to explore how the characteristics of the pathogen and a second host affect disease prevalence in a focal host. Our work helps explain some of the variation in predictions and outcomes in previous empirical and theoretical studies on the dilution effect. One main contribution of our work is that we show models for environmental transmission (ET), density-dependent direct transmission (DDDT), and frequency-dependent direct transmission (FDDT) can be unified, which in turn yields insight into how the pathogen transmission mechanism affects disease prevalence (Figure 1). Dilution effect theory for pathogens with different transmission modes has largely developed independently, masking the effects the transmission mode has on amplification and dilution. Our work allows us to identify the conditions under which a particular transmission mode will lead to higher or lower prevalence in a focal host (Table 1). In addition, by using common metrics to classify host species (competence and sink/source), our work can help explain why many previous theoretical studies predict FDDT promotes dilution more than DDDT. Previous studies on DDDT pathogens show amplification always occurs, unless there is interspecific competition between hosts (Begon et al., 1992; Rudolf and Antonovics, 2005; Faust et al., 2017; Roberts and Heesterbeek, 2018). The reason for this is that all hosts are sources under DDDT, which implies sufficiently strong interspecific competition is necessary for dilution. For studies on FDDT pathogens, dilution always occurs unless the host has very high transmission rates (Rudolf and Antonovics, 2005; Faust et al., 2017). This is because those studies use transmission rates like 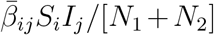] that correspond to the second host having small *χ*_2_ values in the ET model and being a sink, unless interspecific transmission is greater than intraspecific transmission. Overall, this means the prior prediction (FDDT promotes dilution more than DDDT) is partially confounded by some hosts being sources and others being sinks. Our work clarifies the prediction for systems with two hosts by showing that dilution occurs more under FDDT than DDDT in the absence of interspecific competition, but dilution can occur more under DDDT than FDDT when interspecific competition is sufficiently high.

Another main contribution of our work is it yields insight into how host characteristics can shape the context-dependent effects of host diversity on disease prevalence. Many previous studies have pointed out that host biodiversity-disease relationships are likely to be context dependent and strongly depend on which specific species are gained or lost from a community (LoGiudice et al., 2008; Randolph and Dobson, 2012; Rohr et al., 2020; Halliday et al., 2020). Our work identifies the context-dependent rules defining when the competence and competitive ability of a second host promote higher versus lower infection prevalence in a focal host (Table 1). Our use of the word ‘promote’ is important because our results do not determine if introduction of the second host is guaranteed to cause an increase or decrease in the prevalence in a focal host. Instead, our results identify which characteristics of the second host and pathogen tend to make one outcome more likely to occur. This is useful because host characteristics can have opposing effects on disease prevalence (Luis et al., 2018). In addition, these results show that some conclusions reported in previous studies are incomplete because they are missing important context-dependent qualifiers. For example, previous studies predict (Rudolf and Antonovics, 2005; Strauss et al., 2015) increased interspecific competition causes decreased prevalence, which is true provided the second host is not a large source (Figure 4).

Our context-dependent rules also help explain differing results from previous empirical studies on the environmentally transmitted fungal pathogen *Metschnikowia bicuspidata*. In one study (Searle et al., 2016), the higher competence host *Daphnia lumholtzi* and the lower competence host *Daphnia dentifera* amplified disease in each other, despite asymmetric interspecific competition. Our results suggest that while interspecific competition between the host species promoted dilution, both hosts were sufficiently large sources that each amplified disease in the other. In contrast, even though there was strong interspecific competition between the host species, higher competence clones of *D. dentifera* amplified disease in the much lower competence clones of *Ceriodaphnia sp*. and *Ceriodaphnia sp*. diluted disease in*D. dentifera* (Case 1 in Strauss et al. (2015)). Our results suggest that this outcome was driven by the high competence *D. dentifera* clones being large sources and the much lower competence *Ceriodaphnia sp*. clones being large sinks. Our results also generally agree with the pattern in Strauss et al. (2015) that the diluting effects of a second host are strongest when the second host has lower competence and is a stronger interspecific competitor than the focal host (compare Cases 1-3 in Strauss et al. (2015)).

While our work helps unify and explain some results from previous studies, particularly those for systems with two host species, our work cannot fully explain all of the variation in empirical results and theoretical predictions observed across the literature. This is partially because studies use different metrics to quantify levels of disease and vary different quantities to assess the effect of host biodiversity on disease. Previous studies have shown that these differences can result in quantitatively and qualitatively different predictions about host biodiversity-disease relationships (Roche et al., 2012; Wood et al., 2014, 2016; Roberts and Heesterbeek, 2018). For example, while increased competence of the second host can cause lower prevalence in the focal host when the second host is a large sink (Figure 2A,C), our numerical calculations in Figure S1 show that for those parameter values the pathogen’s basic reproductive number (*ℛ*_0_) always increases with increased competence of the second host. Differences between metrics also arise in empirical studies. For example, infection prevalence and density changed in the same direction for each host in Strauss et al. (2015), but infection prevalence and density changed in opposite directions for *D. dentifera* in Searle et al. (2016). Identifying why the metrics differ is important because it would highlight the potentially different ways host biodiversity affects disease dynamics. In addition, because calculations for *ℛ*_0_-based metrics are often simpler than calculations for the other metrics, it would identify when predictions from the simpler method will suffice or be insufficient.

Altogether, this points to the need for new theory that explains how and why predictions differ when different approaches and metrics of disease are used. Importantly, to be able to explain how host diversity affects disease, that theory will need to address and build upon the theory for communities with more than two host species (Roche et al., 2012; Faust et al., 2017; Dobson, 2004; Joseph et al., 2013; Mihaljevic et al., 2014). This will require accounting for correlations between species traits. For example, propagule uptake and resource consumption rates of *Daphnia* species are affected by the host filtering rate (Hall et al., 2007; Dallas et al., 2016). Similar correlations may be present in insects (Evans and Entwistle, 1987; Naug, 2014), snails (Lafferty, 1993; Miura et al., 2006), and grazing mammals (Williams and Barker, 2008; Wobeser, 2013) that consume their environmentally transmitted pathogens or encounter them while foraging (Hall et al., 2007). In addition, new theory will need to move beyond equilibrium-based analyzes (like this study or *ℛ*_0_-based studies) and help explain how and why the effects of host biodiversity on disease may fluctuate in time (Dallas et al., 2016).

Developing the new theory will not be trivial, but we think the approaches in this study point to possible ways forward. First, our results show that pathogens with ET, DDDT, and FDDT pathogens can be studied simultaneously with a single model. Doing so will yield understanding about how the pathogen transmission mode affects host biodiversity-disease relationships. In addition, extending our approach or applying the framework in Lafferty et al. (2015) may provide a way to unify dilution effect theory for environmental, direct, and vector-borne transmission. Second, our Jacobian-based method can be adapted for other metrics and approaches that vary different quantities; examples can be found in both predator-prey studies (Abrams and Nakajima, 2007; Abrams and Cortez, 2015) and host-pathogen studies (Cortez and Duffy, 2020; Clay et al., submitted). A key challenge will be dealing with the increased dimensionality of communities with more than two species. We have shown that responses in disease prevalence can be interpreted in terms of indirect pathways (Figure 3), and this may provide simple interpretations to complex mathematical expressions. In total, while there is much work to be done, this study helps us move towards dilution effect theory that identifies the general rules governing how the characteristics of host and pathogens shape host biodiversity-disease relationships.

## 5 Acknowledgments

We thank Patrick Clay, the FSU Ecology Reading Group, and three anonymous reviewers for helpful comments on previous versions of the paper. MAD and MHC were supported by the National Science Foundation under Awards DEB-1748729 & DEB-2015280.

