## Appendices for "The context dependent effects of host competence, competition, and the pathogen transmission mode on disease prevalence"

### Supplementary Material for 'Toward the unification of dilution effect theory for environmental and direct transmission pathogens'

Michael H. Cortez & Meghan A. Duffy

S1.4.3 Change of parameters that holds the multi-species equilibrium constant 9
S1.4.4 Change of parameters that holds the single-species equilibrium constant 11
S1.5 Response of the single-species equilibrium prevalence to parameter variation 14
S1.6 Response of the multi-species equilibrium prevalence to parameter variation . 15
S1.6.1 Response to increased excretion or removal of infectious propagules . 15
S1.6.2 Response to increased mortality of the second host . . . 17
S1.6.3 Response to increased infection rates of the second host . . . 18
S1.6.4 Response to increased intraspecific competition in the second host . . 19
S1.6.5 Response to increased interspecific competitive ability of the second host 20

**S2 Analysis of SIRS Environmental Transmission Model 22**
S2.4 Response of the multi-species equilibrium prevalence to parameter variation . 29

S2.4.1 Response to increased excretion or removal of infectious propagules . 29
S2.4.2 Response to increased disease-induced mortality of the second host . 31
S2.4.3 Response to increased intraspecific competition in the second host . . 32
S2.4.4 Response to the focal host experiencing increased competition: . . . . 33
S2.4.5 Comparison of responses in SI an SIRS models . . . . . . . . . . . . . 34

**S3 Supplementary Figure** **35**

**S4 Figure equations and parameters** **36**

### S1 Analysis of SI Environmental Transmission Model

#### S1.1 Model assumptions and notation

To simplify the presentation of our results, we absorb the natural mortality rate for each host ($m_i$) into the host reproduction rates ($F_i$) and the disease-induced mortality rates ($\mu_i$). This means that in the equations, $F_i$ should be interpreted as $F_i - m_i$ and $\mu_i$ should be interpreted as $m_i + \mu_i$. We assume the growth rates for each host can be written as

$$F_i(\cdot) = S_i f_{S_i}(\cdot) + I_i f_{I_i}(\cdot) \tag{S1}$$

where $(\cdot)$ is shorthand for $(S_1, S_2, I_1, I_2)$, $f_{I_i}$ is the per capita reproductive rate of infected individuals, and $f_{S_i}$ is the sum of the natural mortality rate and the per capita reproductive rate of susceptible individuals. We assume $\partial f_{X_i}/\partial Y_j < 0$ for all $i, j$ and any host classes $X, Y \in \{S, I\}$ because of intraspecific and interspecific competition.

We assume the multi-species model (1) has a unique stable endemic coexistence equilibrium, $p^* = (S_1^*, S_2^*, I_1^*, I_2^*, P^*)$. Removing the $dS_2/dt$ and $dI_2/dt$ equations and setting $S_2 = I_2 = 0$ in model (1) yields a single-species model where only the focal host is present. We assume the single-species model has a unique stable endemic equilibrium, $\hat{p} = (\hat{S}_1, \hat{I}_1, \hat{P})$. Total host density is denoted $N_i = S_i + I_i$ and the per spore total infectious propagule uptake rate is denoted $U = \sum_i u_{S_i} S_i + u_{I_i} I_i$. The equilibrium values are denoted $N_i^*$, $\hat{N}_i$, $U^*$ and $\hat{U}$.

As described in the main text, we measure host competence by $\mathcal{R}_{0,i}(\infty) = \beta_i \chi_i / \mu_i u_{S_i}$. That quantity is the pathogen's basic reproductive number for the single-species system, $\mathcal{R}_{0,i} = \beta_i \chi_i \bar{S}_i / \mu_i(\delta + u_{S_i} \bar{S}_i)$, in the limit where the susceptible density at the single-species disease free equilibrium ($\bar{S}_1$) is infinite. We assume $\mathcal{R}_{0,i} > 1$, which implies $\mathcal{R}_{0,i}(\infty) > 1$. In general, this is not a necessary condition for the existence of the single-species endemic equilibrium, $\hat{p}$, however in many common epidemiological models, $\mathcal{R}_{0,i} > 1$ is a necessary condition.

Our analysis primarily focuses on systems where both host species experience negative density dependence at the single-species and multi-species equilibria. Mathematically, we assume the growth rate of host $i$ at equilibrium decreases with an increase in the density of any host class, i.e., $\partial F_i/\partial S_i < 0$ and $\partial F_i/\partial I_i < 0$ for $i = 1, 2$ when evaluated at $\hat{p}$ or $p^*$. We then discuss how our results can differ if one or both hosts experiences positive density dependence. Mathematically, positive density dependence is defined by $\partial F_i/\partial X_i > 0$ for some host class $X_i$. Positive density dependence can arise if the pathogen or competition suppresses host densities to very low values.

##### S1.1.1 Single-species equilibrium and Jacobian

The Jacobian evaluated at the single-species equilibrium, $\hat{p}$, is

$$\hat{J} = \left.\begin{pmatrix} \frac{\partial f_1}{\partial S_1} - \beta_1 P & \frac{\partial f_1}{\partial I_1} & -\beta_1 S_1 \\ \beta_1 P & -\mu_1 & \beta_1 S_1 \\ -u_{S_1} P & \chi_1 - u_{I_1} P & -U - \delta \end{pmatrix}\right|_{\hat{p}} \tag{S2}$$

with sign structure

$$\begin{pmatrix} - & \pm & - \\ + & - & + \\ - & + & - \end{pmatrix} \tag{S3}$$

where $\pm$ means the entry can have a positive or negative sign. Stability of $\hat{p}$ implies $|\hat{J}| < 0$. The entry $\hat{J}_{11}$ is negative because

$$\hat{J}_{11} = f_{S_1} - \beta_1 P + S_1 \frac{\partial f_{S_1}}{\partial S_1} + I_1 \frac{\partial f_{I_1}}{\partial S_1}\bigg|_{\hat{p}} = -\frac{I_1 f_{I_1}(\cdot)}{S_1} + S_1 \frac{\partial f_{S_1}}{\partial S_1} + I_1 \frac{\partial f_{I_1}}{\partial S_1}\bigg|_{\hat{p}} < 0 \tag{S4}$$

where the last equality follows from the fact that at equilibrium $dS_1/dt = 0$, which means $f_{S_1}(\cdot) - \beta_1 \hat{P} = -\hat{I}_1 f_{I_1}(\cdot)/\hat{S}_1$. While $\hat{J}_{11}$ is negative, entry $\hat{J}_{12}$ can have either sign because $\partial(I_1 f_{I_1})/\partial I_1$ can be positive or negative depending on the densities of the host classes. Whenever we assume negative density dependence, $\hat{J}_{12}$ will be negative.

##### S1.1.2 Multi-species equilibrium and Jacobian

The Jacobian evaluated at the multi-species equilibrium, $p^*$, is

$$J = \begin{pmatrix} \frac{\partial f_1}{\partial S_1} - \beta_1 P & \frac{\partial f_1}{\partial S_2} & \frac{\partial f_1}{\partial I_1} & \frac{\partial f_1}{\partial I_2} & -\beta_1 S_1 \\ \frac{\partial f_2}{\partial S_1} & \frac{\partial f_2}{\partial S_2} - \beta_2 P & \frac{\partial f_2}{\partial I_1} & \frac{\partial f_2}{\partial I_2} & -\beta_2 S_2 \\ \beta_1 P & 0 & -\mu_1 & 0 & \beta_1 S_1 \\ 0 & \beta_2 P & 0 & -\mu_2 & \beta_2 S_2 \\ -u_{S_1} P & -u_{S_2} P & \chi_1 - u_{I_1} P & \chi_2 - u_{I_2} P & -U - \delta \end{pmatrix}\Bigg|_{p^*} \tag{S5}$$

with sign structure

$$\begin{pmatrix} - & - & \pm & - & - \\ - & - & - & \pm & - \\ + & 0 & - & 0 & + \\ 0 & + & 0 & - & + \\ - & - & \pm & \pm & - \end{pmatrix} \tag{S6}$$

where $\pm$ means the entry can have a positive or negative sign. Stability of $p^*$ implies $|J| < 0$. Entries $J_{11}$ and $J_{22}$ are negative for the same reasons as in the single-species model. Whenever we assume negative density dependence, $J_{13}$ and $J_{24}$ will be negative.

#### S1.2 Method for computing equilibrium responses to parameter variation

We use the Jacobian-based theory in Bender et al. (1984), Yodzis (1988), Novak et al. (2011) and Cortez and Abrams (2016) to compute how changes in a parameter affect equilibrium densities. Let $x_i$ $(i = 1, ..., n)$ be the variables of the model, $dx_i/dt = F_i(\cdot)$. Let $J$ be the Jacobian of the model evaluated at an equilibrium point $(x_1^*, ..., x_n^*)$.

If $a$ is a parameter of the model that only affects the equation for $x_j$, i.e., only $F_j(\cdot)$ depends on $a$, then the change in $x_i^*$ with a small change in the parameter $a$ is defined by the derivative,

$$\frac{\partial x_i^*}{\partial a} = -\frac{\partial F_j}{\partial a}(J^{-1})_{ji} = -\frac{\partial F_j}{\partial a}\frac{(-1)^{i+j}M_{ji}}{|J|} \tag{S7}$$

where $M_{ji}$ is the $j, i$ minor of the Jacobian (i.e., the determinant of the submatrix of $J$ where row $j$ and column $i$ are removed). When we are only interested in the sign of the derivative, we write

$$\frac{\partial x_i^*}{\partial a} \propto (-1)^k M_{ji}/|J| \tag{S8}$$

where "$\propto$" means "proportional to", $(-1)^k = \text{sgn}(-\frac{\partial F_j}{\partial a})(-1)^{1+i+j}$, and sgn$(\cdot)$ is the sign function.

If $a$ is a parameter that affects the equations for a set $Q$ of the variables, i.e., $F_j(\cdot)$ depends on $a$ if $j \in Q$, then the change in $X_i^*$ with a small change in the parameter $a$ is defined by the sum of derivatives,

$$\frac{\partial x_i^*}{\partial a} = \sum_{j \in Q} -\frac{\partial F_j}{\partial a}(J^{-1})_{ji} = \sum_{j \in Q} -\frac{\partial F_j}{\partial a}\frac{(-1)^{i+j}M_{ji}}{|J|}. \tag{S9}$$

#### S1.3 Relationships between environmental and direct transmission models

To show the ET model (1) can reduce to a DDDT model, we assume the excretion and degradation rates are much larger than the uptake rates and sufficiently large that there is a separation of time scales between the dynamics of the infectious propagules and the dynamics of the host classes. Mathematically, we assume the infectious propagule equation can be written as

$$\frac{dP}{dt} = \sum_i \frac{\chi_i}{\epsilon} I_i - \sum_i (u_{S_i} S_i + u_{I_i} I_i) P - \frac{\delta}{\epsilon} P \tag{S10}$$

where $\chi_i$ and $\delta$ are order one and $\epsilon$ is a positive parameter much smaller than one in magnitude (i.e., $0 < \epsilon \ll 1$). Under these assumptions, the dynamics of the infectious propagules reach a quasi-steady state equilibrium density, $P = \sum_i \chi_i I_i / \delta$, which is found by multiplying both sides of the previous equation by $\epsilon$, setting $\epsilon$ equal to zero, and solving for $P$. Substitution into the equations for the host dynamics yields a DDDT model

$$\begin{aligned} \frac{dS_i}{dt} &= F_i(S_1, S_2, I_1, I_2) - m_i S_i - \sum_j \bar{\beta}_{ij} S_i I_j \\ \frac{dI_i}{dt} &= \sum_j \bar{\beta}_{ij} S_i I_j - (m_i + \mu_i) I_i \end{aligned} \tag{S11}$$

where the direct transmission coefficients are $\bar{\beta}_{ji} = \beta_i \chi_j / \delta$. Note that changes in the values the $u_{S_i}$ and $u_{I_i}$ do not affect the dynamics of model (S11) because the model does not depend on $u_{Ii)}$ and any changes in $u_{S_i}$ are counteracted by changes in $p_i$.

To show ET model (1) reduces to a FDDT model, we assume the excretion and host uptake rates are much larger than the degradation rate and sufficiently large that there is a separation of time scales between the dynamics of the infectious propagules and the dynamics of the host classes. Mathematically, we assume the infectious propagule equation can be written as

$$\frac{dP}{dt} = \sum_i \frac{\chi_i}{\epsilon} I_i - \sum_i \left( \frac{u_{S_i}}{\epsilon} S_i + \frac{u_{I_i}}{\epsilon} I_i \right) P - \delta P \tag{S12}$$

where $\chi_i$, $u_{S_i}$, and $u_{I_i}$ are order one and $\epsilon$ is a positive parameter much smaller than one in magnitude (i.e., $0 < \epsilon \ll 1$). Under these assumptions, the dynamics of the infectious propagules reach a quasi-steady state equilibrium density, $P = (\sum_i \chi_i I_i)/(\sum_i u_{S_i} S_i + u_{I_i} I_i)$, which is found by multiplying both sides of the previous equation by $\epsilon$, setting $\epsilon$ equal to zero, and solving for $P$. Substitution into the equations for the host dynamics yields a FDDT model

$$\begin{aligned} \frac{dS_i}{dt} &= F_i(S_1, S_2, I_1, I_2) - m_i S_i - \frac{\sum_j \bar{\beta}_{ij} S_i I_j}{\sum_j u_{S_j} S_j + u_{I_j} I_j} \\ \frac{dI_i}{dt} &= \frac{\sum_j \bar{\beta}_{ij} S_i I_j}{\sum_j u_{S_j} S_j + u_{I_j} I_j} - (m_i + \mu_i) I_i \end{aligned} \tag{S13}$$

where $\bar{\beta}_{ij} = \beta_i \chi_j$ and the densities of the host classes are weighted by their uptake rates.

#### S1.4 Response of the equilibrium prevalence to changes in the transmission mode

The first subsection shows how to construct the change of parameter functions. The second subsection provides interpretation for an important quantity. The third section computes how equilibrium prevalence responds to changes in the transmission mode.

##### S1.4.1 Constructing changes of parameter functions

Here we show how to determine the change of parameters that converts the environmental transmission model from a form that behaves like a frequency-dependent direct transmission (FDDT) model to a form that behaves like a density-dependent direct transmission (DDDT) model, while holding some specific quantity fixed.

The first step is to choose a quantity to hold fixed. Possible quantities include $\mathcal{R}_{0,i}$, the single-species equilibrium ($\hat{p}$), or the multi-species equilibrium ($p^*$). The second step is to write the infectious propagule equation as

$$\frac{dP}{dt} = \chi_1 I_1 + \chi_2 I_2 - f(q)(u_{S_1} S_1 + u_{I_1} I_1 + u_{S_2} S_2 + u_{I_2} I_2)P - q\delta P. \tag{S14}$$

The function $f(q)$ is to be determined. It will be constructed such that $f(q) > 0$ for $0 \leq q < q_1$ and $f(q_1) = 0$ for some $q_1 > 0$. When $f(q) = 0$, the ET model has a form that behaves like a DDDT model. When $q = 0$, the ET model has a form that behaves like a FDDT model. The third step is to compute the chosen quantity using the new infectious propagule equation (S14), set it equal to the fixed value, and solve for $f(q)$. For example, if $\mathcal{R}_{0,i}$ is held constant, then $f(q)$ must satisfy

$$\frac{\beta_i \chi_i \hat{S}_i}{\mu_i (f(q) u_{S_i} \hat{S}_i + q\delta_i)} = \frac{\beta_i \chi_i \hat{S}_i}{\mu_i (u_{S_i} \hat{S}_i + \delta_i)}. \tag{S15}$$

Solving yields $f(q) = 1 + (\delta_i / u_{S_i} \hat{S}_i)(1 - q)$ for $0 \leq q \leq 1 + u_{S_i} \hat{S}_i / \delta_i$.

Our analysis focuses on holding the single-species equilibrium ($\hat{p}$) or the multi-species equilibrium ($p^*$) fixed. Holding the single-species equilibrium ($\hat{p}$) fixed, only requires holding constant the per spore loss rate at the single-species equilibrium ($\hat{U} - \delta$). This means $f(q)$ satisfies

$$f(q)(u_{S_1} \hat{S}_1 + u_{I_1} \hat{I}_1) + q\delta = (u_{S_1} \hat{S}_1 + u_{I_1} \hat{I}_1) + \delta \tag{S16}$$

which implies $f(q) = 1 + \frac{\delta}{\hat{U}} - \frac{\delta}{\hat{U}} q$ for $0 \leq q \leq 1 + U^*/\delta$. Similarly, holding the multi-species equilibrium ($\hat{p}$) fixed, only requires holding constant the per spore loss rate at the multi-species equilibrium ($U^* - \delta$). This means $f(q)$ satisfies

$$f(q) \left( \sum_i u_{S_i} S_i^* + u_{I_i} I_i^* \right) + q\delta = \sum_i u_{S_1} S_i^* + u_{I_1} I_i^* + \delta \tag{S17}$$

which implies $f(q) = 1 + \frac{\delta}{U^*} - \frac{\delta}{U^*} q$ for $0 \leq q \leq 1 + U^*/\delta$.

##### S1.4.2 Factors affecting the sign of $U^* - \hat{U}$

Here, we explore the factors that affect the sign of $U^* - \hat{U}$. Using the equilibrium conditions $dP/dt = 0$ and $dI_1/dt = 0$ for the single-species and multi-species models, we can rewrite $U^* - \hat{U}$ as

$$U^* - \hat{U} = \frac{\chi_1 I_1^*}{P^*} + \frac{\chi_2 I_2^*}{P^*} - \frac{\chi_1 \hat{I}_1}{\hat{P}} \tag{S18}$$

$$= \frac{\chi_1 \beta_1}{\mu_1}(S_1^* - \hat{S}_1) + \frac{\chi_2 \beta_2}{\mu_2} S_2^*. \tag{S19}$$

Before giving general conditions, we start with a few special cases.

<u>Special case 1 - No interspecific host competition:</u> In this case, the magnitudes of $S_1^*$ and $S_2^*$ are determined by the competence and intraspecific competitive ability of each host. $U^* - \hat{U}$ is more likely to be positive when the second host has lower competence and lower intraspecific competitive ability, because both of these make $S_1^*$ and $S_2^*$ larger. $U^* - \hat{U}$ is more likely to be negative when the second host has higher competence and higher intraspecific competitive ability, because both of these make $S_1^*$ and $S_2^*$ smaller.

<u>Special case 2 - Symmetric uptake:</u> Assuming $u_{S_i} = u_{I_i} = u_i$ results in

$$U^* - \hat{U} = u_1(N_1^* - \hat{N}_1) + u_2 N_2^* \tag{S20}$$

where $N_i = S_i + I_i$. $N_i^*$ is lower with increased interspecific competition, increased intraspecific competitive ability of host $i$, and increased density of infected individuals in host $i$. This means $U^* - \hat{U}$ is more likely to be positive when interspecific competition is weak, the second host is a weaker intraspecific competitor, and the second host is a lower competence host. $U^* - \hat{U}$ is more likely to be negative when interspecific competition is strong, and the second host is a strong intraspecific competitor and a higher competence host.

<u>Special case 3 - No uptake by infected individuals:</u> Assuming $u_{I_i} = 0$ yields

$$u_{S_1}(S_1^* - \hat{S}_1) + u_{S_2} S_2 = U^* - \hat{U} = \frac{\chi_1 \beta_1}{\mu_1}(S_1^* - \hat{S}_1) + \frac{\chi_2 \beta_2}{\mu_2} S_2^*. \tag{S21}$$

Setting the left and right hand sides equal to each other, solving for $S_1^* - \hat{S}_1$, substituting into equation (S19), and some algebraic manipulation yields

$$U^* - \hat{U} = u_{S_2} S_2^* \left(1 - \frac{\frac{\chi_2 \beta_2}{\mu_2 u_{S_2}} - 1}{\frac{\chi_1 \beta_1}{\mu_1 u_{S_1}} - 1}\right) = u_{S_2} S_2^* \left(1 - \frac{\mathcal{R}_{0,2}(\infty) - 1}{\mathcal{R}_{0,1}(\infty) - 1}\right), \tag{S22}$$

where $\mathcal{R}_{0,i}(\infty) = \chi_i \beta_i / \mu_i u_{I_i}$ is our measure of host competence. Recall that we assume $\mathcal{R}_{0,1}(\infty) > 1$.

Under the assumption that $\mathcal{R}_{0,1}(\infty) > 1$, equation (S22) is positive whenever $\mathcal{R}_{0,1}(\infty) > \mathcal{R}_{0,2}(\infty)$ and equation (S22) is negative whenever $\mathcal{R}_{0,1}(\infty) < \mathcal{R}_{0,2}(\infty)$. Thus, $U^* - \hat{U}$ is positive when the second host is a lower competence host than the focal host and $U^* - \hat{U}$ is negative when the second host is a higher competence host than the focal host .

General Case: In general, $S_i^*$ will be smaller in magnitude when host $j$ is a stronger interspecific competitor, host $i$ is a stronger intraspecific competitor, and host $j$ ($j \neq i$) is a higher competence host. Thus, $U^* - \hat{U}$ is more likely to be positive when (i) interspecific competition is weaker, (ii) the second host is a weaker intraspecific competitor, and (iii) the second host is a lower competence host. In contrast, $U^* - \hat{U}$ is more likely to be negative when (i) interspecific competition is stronger, (ii) the second host is a stronger intraspecific competitor, and (iii) the second host is a higher competence host.

##### S1.4.3 Change of parameters that holds the multi-species equilibrium constant

We use a change of parameters to convert the ET model from a form that behaves like a FDDT model to a form that behaves like a DDDT model, while leaving the host and infectious propagule densities at the multi-species equilibrium unchanged. The specific change of parameters is $f(q) = 1 + \frac{\delta}{U^*} - \frac{\delta}{U^*}q$ for $0 \leq q \leq 1 + U^*/\delta$; see section S1.4.1 for details. With this change of parameters, the densities at the multi-species equilibrium ($p^*$) are independent of the value of $q$ and the densities at the single-species equilibrium ($\hat{p}$) change as $q$ is varied.

**Effect on loss rate at single-species equilibrium:** To see how varying $q$ affects the total per spore loss rate of infectious propagules in the single-species model, we can compute the difference in the loss rates at equilibrium for $q = 1$ and $q = 1 + U^*/\delta$,

$$[\text{loss rate for q=}1 + U^*/\delta] - [\text{loss rate for q=1}] = U^* + \delta - (\hat{U} + \delta) = U^* - \hat{U}. \quad \text{(S23)}$$

Similarly, we can compute the difference in the loss rates at equilibrium for $q = 0$ and $q = 1$,

$$[\text{loss rate for q=1}] - [\text{loss rate for q=0}] = \hat{U} + \delta - \left(1 + \frac{\delta}{U^*}\right)\hat{U} = \frac{\delta}{U^*}(U^* - \hat{U}). \quad \text{(S24)}$$

In both cases, the effect of increasing $q$ on the total per capita loss rate of infectious propagules depends on the sign of $U^* - \hat{U}$.

Another way to see how varying $q$ affects the per capita infectious propagule loss rate is to compute the partial derivative,

$$\frac{\partial}{\partial q}\frac{dP}{dt} = \frac{\partial}{\partial q}\left[\chi_1 I_1 - f(q)(u_{S_1}S_1 + u_{I_1}I_1) - q\delta P\right] \quad \text{(S25)}$$

$$= \frac{\delta}{U^*}UP - \delta P = -P\delta(U^* - U)/U^*. \quad \text{(S26)}$$

When evaluated at the single-species equilibrium, $\hat{p}$, the above shows that the effect of increasing $q$ depends on the sign of $U^* - \hat{U}$.

In total, if $U^* - \hat{U}$ is positive, then the total per capita loss rate will increase as the environmental transmission model is changed from a form that behaves like a FDDT model ($q = 0$) into a form that behaves like a DDDT model ($q = 1 + \hat{U}/\delta$). If $U^* - \hat{U}$ is negative, then the opposite is true.

**Effect on single-species equilibrium densities:** The effects of varying $q$ on the densities at the single-species equilibrium, $\hat{p}$, are

$$\frac{\partial \hat{I}_1}{\partial q} = -\frac{\partial}{\partial q}\left(\frac{dP}{dt}\right)\frac{(-1)^{2+3}M_{32}}{|\hat{J}|} \propto -(U^* - \hat{U})\frac{M_{32}}{|J|} = -(U^* - \hat{U})\frac{\beta_1 S_1}{|\hat{J}|}\frac{\partial f_1}{\partial S_1}, \tag{S27}$$

$$\frac{\partial \hat{S}_1}{\partial q} = -\frac{\partial}{\partial q}\left(\frac{dP}{dt}\right)\frac{(-1)^{1+3}M_{31}}{|\hat{J}|} \propto (U^* - \hat{U})\frac{M_{31}}{|\hat{J}|} = (U^* - \hat{U})\frac{\beta_1 S_1}{|\hat{J}|}\frac{\partial f_1}{\partial S_1} \tag{S28}$$

and

$$\frac{\partial}{\partial q}\left(\frac{\hat{I}_1}{\hat{N}_1}\right) = \frac{1}{\hat{N}_1^2}\left[\hat{S}_1\frac{\partial \hat{I}_1}{\partial q} - \hat{I}_1\frac{\partial \hat{S}_1}{\partial q}\right] \propto -(U^* - \hat{U})\frac{1}{|\hat{J}|}\left[\hat{S}_1\frac{\partial f_1}{\partial S_1} + \hat{I}_1\frac{\partial f_1}{\partial I_1}\right] \tag{S29}$$

where $|\hat{J}| < 0$, $M_{31}$ and $M_{32}$ are minors of matrix (S2), and all terms are evaluated at $\hat{p}$.

Negative density dependence of the focal host at equilibrium: We assume that the focal host has negative density dependence at the single-species equilibrium, i.e., $\partial f_1/\partial X_1 < 0$ for $X \in \{S, I\}$.

If $U^* - \hat{U} > 0$, then equation (S29) is negative. This implies that focal host infection prevalence is higher at the single-species equilibrium under FDDT than DDDT, which means $(I_1^*/N_1^*) - (\hat{I}_1/\hat{N}_1)$ is more negative under FDDT than DDDT. From this we predict that the introduction of the second host dilutes disease more or amplifies disease less in the focal host under FDDT than DDDT when:

- (D1) Interspecific competition is weaker, (D2) the second host is a weaker intraspecific competitor, and (D3) the second host a lower competence host.

If $U^* - \hat{U} < 0$, then equation (S29) is positive. This implies focal host infection prevalence is lower at the single-species equilibrium under FDDT than DDDT, which means $(I_1^*/N_1^*) - (\hat{I}_1/\hat{N}_1)$ is more positive under FDDT than DDDT. From this we predict that the introduction of the second host dilutes disease more or amplifies disease less in the focal host under DDDT than FDDT when:

- (D4) Interspecific competition is stronger, (D5) the second host is a stronger intraspecific competitor, and (D6) the second host a higher competence host.

Positive density dependence of the focal host at equilibrium: Sufficiently strong positive density dependence in the focal host can reverse the above predictions.

##### S1.4.4 Change of parameters that holds the single-species equilibrium constant

Now, we use a change of parameters to convert the ET model from a form that behaves like a FDDT model to a form that behaves like a DDDT model, while holding fixed the densities at the single-species equilibrium. The specific change of parameters is $f(q) = 1 + \frac{\delta}{\hat{U}} - \frac{\delta}{\hat{U}}q$ for $0 \leq q \leq 1 + \hat{U}/\delta$; see section S1.4.1 for details. With this change of parameters, the densities at the single-species equilibrium ($\hat{p}$) are independent of the value of $q$ and the densities at the multie-species equilibrium ($p^*$) change as $q$ is varied.

**Effect on loss rate at multi-species equilibrium:** To see how varying $q$ affects the total per spore rate of loss of infectious propagules in the multi-species models, we can compute the difference in the loss rates at equilibrium when $q = 1$ and when $q = 1 + \hat{U}/\delta$. This yields

$$[\text{loss rate for q=}1 + \hat{U}/\delta] - [\text{loss rate for q=1}] = \hat{U} + \delta - (U^* + \delta) = \hat{U} - U^*. \tag{S30}$$

Similarly, we can compute the difference in the loss rates at equilibrium for $q = 0$ and $q = 1$. This yields

$$[\text{loss rate for q=1}] - [\text{loss rate for q=0}] = U^* + \delta - \left(1 + \frac{\delta}{\hat{U}}\right) U^* = \frac{\delta}{\hat{U}}(\hat{U} - U^*). \tag{S31}$$

In both cases, the effect of increasing $q$ on the total per capita loss rate of infectious propagules depends on the sign of $\hat{U} - U^*$.

Another way to see how varying $q$ affects the per capita infectious propagule loss rate is to compute the partial derivative,

$$\begin{aligned} \frac{\partial}{\partial q}\frac{dP}{dt} &= \frac{\partial}{\partial q}\left[\chi_1 I_1 + \chi_2 I_2 - f(q)(u_{S_1}S_1 + u_{I_1}I_1 + u_{S_2}S_2 + u_{I_2}I_2) - q\delta P\right] && \text{(S32)} \\ &= \frac{\delta}{\hat{U}}UP - \delta P = P\delta\hat{U}(U - \hat{U}). && \text{(S33)} \end{aligned}$$

When evaluated at the multi-species equilibrium, $p^*$, the above shows that the effects of increasing $q$ depends on the sign of $U^* - \hat{U} < 0$.

In total, if $U^* - \hat{U}$ is negative, then the total per capita loss rate will increase as the ET model is continuously changed from a form that behaves like a FDDT model ($q = 0$) into a form that behaves like a DDDT model ($q = 1 + \hat{U}/\delta$). If $U^* - \hat{U}$ is positive, then the opposite is true.

**Effect on multi-species equilibrium densities:** The effects of varying $q$ on the densities at the multi-species equilibrium are

$$\begin{aligned} \frac{\partial I_1^*}{\partial q} &= -\frac{\partial}{\partial q}\left(\frac{dP}{dt}\right)\frac{(-1)^{3+5}M_{53}}{|J|} \\ &\propto -(U^* - \hat{U})\left[\frac{\beta_1 S_1}{|J|}\left(\mu_2\frac{\partial f_1}{\partial S_1}\frac{\partial f_2}{\partial S_2} + \beta_2 P\frac{\partial f_1}{\partial S_1}\frac{\partial f_2}{\partial I_2} - \mu_2\beta_2 P\frac{\partial f_1}{\partial S_1}\right)\right. \\ &\quad \left. + \frac{\beta_1\beta_2 P}{|J|}\left(\frac{\partial f_2}{\partial I_2}S_2\frac{\partial f_1}{\partial S_2} - \frac{\partial f_1}{\partial I_2}\left(S_1\frac{\partial f_2}{\partial S_1} + S_2\frac{\partial f_2}{\partial S_2}\right)\right) - \frac{\beta_1 S_1\mu_2}{|J|}\frac{\partial f_1}{\partial S_2}\frac{\partial f_2}{\partial S_1}\right], \end{aligned} \tag{S34}$$

$$\frac{\partial S_1^*}{\partial q} = -\frac{\partial}{\partial q}\left(\frac{dP}{dt}\right)\frac{(-1)^{1+5}M_{51}}{|J|} \propto -(U^* - \hat{U})\frac{M_{51}}{|J|}, \tag{S35}$$

and

$$\begin{aligned}\frac{\partial}{\partial q}\left(\frac{I_1^*}{N_1^*}\right) \propto &-(U^* - \hat{U})\left[\frac{P\beta_1^2 S_1^2}{\mu_1|J|}\frac{\partial f_1}{\partial I_1}\left(P\beta_2\frac{\partial f_2}{\partial I_2} - P\beta_2\mu_2 + \mu_2\frac{\partial f_2}{\partial I_2}\right)\right.\\ &+ \frac{\beta_1\beta_2 S_1^2 P}{|J|}\frac{\partial f_2}{\partial I_2}\left(\frac{\partial f_1}{\partial I_1} - \beta_2\right) + \frac{\beta_1\mu_2 S_1^2}{|J|}\left(\frac{\partial f_1}{\partial S_1} - \beta_2 P\right)\left(\frac{\partial f_2}{\partial S_2} - \beta_2 P\right)\\ &\left.- \frac{\beta_1 S_1^2}{\mu_1|J|}\left(P\beta_2\frac{\partial f_1}{\partial I_2} + \mu_2\frac{\partial f_1}{\partial S_2}\right)\left(P\beta_2\frac{\partial f_2}{\partial I_1} + \mu_1\frac{\partial f_2}{\partial S_1}\right)\right]\end{aligned} \tag{S36}$$

where all terms are evaluated at $p^*$ and $M_{51}/|J|$ is given in equation (S43).

Negative density dependence at equilibrium: We assume both hosts have negative density dependence at equilibrium, i.e., $\partial F_i/\partial X_i < 0$ for $X \in \{S, I, R\}$. Recall from subsection S1.4.1 that the terms in brackets in equations (S36) are negative when interspecific host competition is sufficiently low and positive when (1) interspecific competition is greater than intraspecific competition or (2) competition between infected and susceptible individuals is stronger than competition between susceptible individuals. Also recall from subsection S1.5.2 that positive values of $U^* - \hat{U}$ are promoted when (a) interspecific competition is weak, and the second host is (b) a low competence host and (c) a weak intraspecific competitor.

Equation (S36) is positive when either (i) $U^* - \hat{U} > 0$ and the terms in brackets are negative or (ii) $U^* - \hat{U} < 0$ and the terms in brackets are positive. In these cases, focal host infection prevalence at the multi-species equilibrium is lower under FDDT than DDDT, which means $I_1^*/N_1^* - \hat{I}_1/\hat{N}_1$ is more negative under FDDT than DDDT. Equation (S36) is negative when either (i) $U^* - \hat{U} < 0$ and the terms in brackets are negative or (ii) $U^* - \hat{U} > 0$ and terms in brackets are positive. In these cases, focal host infection prevalence at the multi-species equilibrium is higher under FDDT than DDDT, which means $I_1^*/N_1^* - \hat{I}_1/\hat{N}_1$ is more positive under FDDT than DDDT.

Combining the above yields the following predictions:

- Assume interspecific competition between hosts is less than intraspecific competition and infected individuals are weaker competitors than susceptible individuals
  - Focal host infection prevalence at the multi-species equilibrium is more likely to be lower under FDDT than DDDT when (D1) interspecific competition is weaker, (D2) the second host is a weaker intraspecific competitor, and (D3) the second host is a lower competence host.
  - Focal host infection prevalence at the multi-species equilibrium is more likely to be lower under DDDT than FDDT when (D4) interspecific competition is stronger, (D5) the second host is a stronger intraspecific competitor, and (D6) the second host is a higher competence host.

- Assume interspecific competition between hosts is greater than intraspecific competition or infected individuals are stronger competitors than susceptible individuals
  - Focal host infection prevalence at the multi-species equilibrium is more likely to be lower under FDDT than DDDT when when (F1) the second host is a stronger intraspecific competitor and (F2) the second host is a higher competence host.
  - Focal host infection prevalence at the multi-species equilibrium is more likely to be lower under DDDT than FDDT when when (F3) the second host is a weaker intraspecific competitor and (F4) the second host is a lower competence host.

We note three things about the above. First, conditions D1-D6 in this section are identical to the conditions with the same labels in section S1.5.3. This agreement is expected because in both cases increased removal of infectious propagules causes the proportion of infected individuals in the focal host to decrease (i.e, $\partial(\hat{I}_1/\hat{N}_1)/\partial\delta$ and $\partial(I_1^*/N_1^*)/\partial\delta$ are both negative). Second, conditions F1-F4 differ from conditions D1-D6 because increased removal of infectious propagules causes focal host infection prevalence at the multi-species equilibrium to increase (i.e, $\partial I_1^*/\partial\delta$ and $\partial(I_1^*/N_1^*)/\partial\delta$ positive). This disagreement is also expected because interspecific host competition does not influence the response to increased removal of infectious propagules in the single-species model; $\partial\hat{I}_1/\partial\delta$ is independent of interspecific competition because there is only one host species in the single-species model.

Third, if interspecific competition is absent and there is negative density dependence in the focal host, it is not possible for the focal host to experience amplification at $q = 0$ and dilution at $q = 1 + \hat{U}/\delta$. The proof by contradiction is the following. Under the assumed negative density dependence, amplification at $q = 0$ and dilution at $q = 1 + \hat{U}/\delta$ is only possible if $U^* - \hat{U} < 0$, which implies $S_1^* - \hat{S}_1 < 0$ and sufficiently large for $0 \leq q \leq 1 + \hat{U}/\delta$. Because interspecific competition is absent, the equilibrium host densities satisfy $0 = f_1(S_1^*, I_1^*) - \mu_1 I_1^*$. The dependence of $S_1^*$ on $I_1^*$ can be computed using implicit differentiation,

$$0 = \frac{\partial f_1}{\partial S_1}\frac{\partial S_1^*}{\partial I_1^*} + \frac{\partial f_1}{\partial I_1} - \mu_1\bigg|_{p^*} \quad \Rightarrow \quad \frac{\partial S_1^*}{\partial I_1^*} = -\left(\frac{\partial f_1}{\partial I_1} - \mu_1\right)\bigg/\frac{\partial f_1}{\partial S_1}\bigg|_{p^*} \tag{S37}$$

The assumed negative density dependence implies $\frac{\partial S_1^*}{\partial I_1^*}$ is negative. Thus, increases in $I_1^*$ imply decreases in $S_1^*$ and vice versa. Combining $\frac{\partial S_1^*}{\partial I_1^*} < 0$ and $S_1^* - \hat{S}_1 < 0$ yields $S_1^* - \hat{S}_1 < 0$ and $I_1^* - \hat{I}_1 > 0$ for all $0 \leq q \leq 1 + \hat{U}/\delta$, which implies $I_1^*/(S_1^* + I_1^*) > \hat{I}_1/(\hat{S}_1 + \hat{I}_1)$ for $0 \leq q \leq 1 + \hat{U}/\delta$. However, this means that the proportion of infected individuals at the multi-species equilibrium is higher than the proportion of infected individuals at the single-species equilibrium for all values of $q$, which contradicts our assumption that dilution occurs at $q = 1 + \hat{U}/\delta$. Thus, in the absence of interspecific host competition, it is not possible for amplification to occur under FDDT ($q = 0$) and dilution to occur under DDDT ($q = 1+\hat{U}/\delta$).

Positive density dependence at equilibrium: If either host has positive density dependence at equilibrium and the positive density dependence is sufficiently large, then the terms in brackets in equations (S34) and (S36) change signs and the sign of $U^* - \hat{U}$ is unchanged. In this case, the above predictions are reversed.

#### S1.5 Response of the single-species equilibrium prevalence to parameter variation

All terms and derivatives in this section are evaluated at $\hat{p}$.

The responses to increased infectious propagule degradation are

$$\frac{\partial \hat{S}_1}{\partial \delta} = (-1)^{1+3}\hat{P}\frac{M_{31}}{|\hat{J}|} = \frac{\hat{P}}{|\hat{J}|}\begin{vmatrix} \frac{\partial f_1}{\partial I_1} & -\beta_1\hat{S}_1 \\ -\mu_1 & \beta_1 S_1 \end{vmatrix} = \frac{\beta_1\hat{S}_1\hat{P}}{|\hat{J}|}\left(\frac{\partial f_1}{\partial I_1} - \mu_1\right) \tag{S38}$$

and

$$\frac{\partial \hat{I}_1}{\partial \delta} = (-1)^{2+3}\hat{P}\frac{M_{32}}{|\hat{J}|} = \frac{-\hat{P}}{|\hat{J}|}\begin{vmatrix} \frac{\partial f_1}{\partial S_1} - \beta_1\hat{P} & -\beta_1\hat{S}_1 \\ \beta_1\hat{P} & \beta_1\hat{S}_1 \end{vmatrix} = \frac{-\beta_1\hat{S}_1\hat{P}}{|\hat{J}|}\frac{\partial f_1}{\partial S_1}. \tag{S39}$$

The response in the proportion of infected individuals is determined using the chain rule,

$$\frac{\partial}{\partial \delta}\left(\frac{\hat{I}_1}{\hat{N}_1}\right) = \frac{1}{\hat{N}_1^2}\left[\hat{S}_1\frac{\partial \hat{I}_1}{\partial \delta} - \hat{I}_1\frac{\partial \hat{S}_1}{\partial \delta}\right] = -\frac{\beta_1\hat{S}_1^2\hat{P}}{\mu_1\hat{N}_1^2|\hat{J}|}\left[\mu_1\left(\frac{\partial f_1}{\partial S_1} - \beta_1\hat{P}\right) + \hat{P}\beta_1\frac{\partial f_1}{\partial I_1}\right], \tag{S40}$$

where we use the identity $\hat{I} = \beta_1\hat{S}_1\hat{P}/\mu_1$ defined by the single-species equilibrium.

The sign of $\partial(\hat{I}_1/\hat{N}_1)/\partial\delta$ is negative under the assumption of negative density dependence. Sufficiently large positive density dependence in the infected class can cause the derivative to be positive.

#### S1.6 Response of the multi-species equilibrium prevalence to parameter variation

In each subsection, we present the derivatives for $\partial S_1^*/\partial a$, $\partial I_1^*/\partial a$, and $\partial(I_1^*/N_1^*)/\partial a$ where $a$ is a parameter. The derivative $\partial(I_1^*/N_1^*)/\partial a$ is computed using the chain rule,

$$\begin{aligned}\frac{\partial}{\partial a}\left(\frac{I_1^*}{N_1^*}\right) &= \frac{1}{(N_1^*)^2}\left[(S_1^*+I_1^*)\frac{\partial I_1^*}{\partial a} - I_1^*\left(\frac{\partial S_1^*}{\partial a}+\frac{\partial I_1^*}{\partial a}\right)\right] = \frac{1}{(N_1^*)^2}\left[S_1^*\frac{\partial I_1^*}{\partial a} - I_1^*\frac{\partial S_1^*}{\partial a}\right] \\ &\propto \left[\frac{\partial I_1^*}{\partial a} - \frac{\beta_1 P^*}{\mu_1}\frac{\partial S_1^*}{\partial a}\right]\end{aligned} \tag{S41}$$

where the last line uses the equilibrium condition $0 = dI_i/dt|_{p^*} = \beta_1 S_1^* P^* - \mu_1 I_1^*$ to substitute for $I_1^*$.

The terms in the equations for $\partial S_1^*/\partial a$ and $\partial I_1^*/\partial a$ are labeled with the corresponding indirect pathway. Indirect pathways are listed as a string of variables and arrows that start and end with different variables (e.g., $P \to S_1 \to I_1$). Indirect pathways that only involve a subset of the variables are multiplied by loops, which are represented by a string of variables and arrows that start and end with the same stage (e.g., $S_2 \to I_2 \to S_2$). For each term, the corresponding indirect pathways and loops are combined using ampersands (e.g., $P \to S_1 \to I_1 \,\&\, S_2 \to I_2 \to S_2$). When terms define multiple sets of indirect effects and loops, each set is separated by a comma. For example, $P \to I_1 \,\&\, S_1 \to S_1 \,\&\, S_2 \to S_2 \,\&\, I_2 \to I_2$, $P \to I_1 \,\&\, S_1 \to S_1 \,\&\, S_2 \to I_2 \to S_2$ in the first line of equation (S42) means that the first term corresponds to $P \to I_1 \& S_1 \to S_1 \& S_2 \to S_2 \& I_2 \to I_2$ and the second term corresponds to $P \to I_1 \,\&\, S_1 \to S_1 \,\&\, S_2 \to I_2 \to S_2$. We do not label all of the terms in the equations for $\partial(I_1^*/N_1^*)/\partial a$ and instead we collect terms based on their signs. This facilitates determining when the equations are positive or negative and adding labeling makes the equations difficult to read.

When determining the signs of the terms in the following equations, it is useful to keep in mind that $U^* + \delta - u_{S_1}S_1^* - u_{S_2}S_2^* > 0$, the effects of interspecific competition show up through the terms $\frac{\partial F_i}{\partial X_j}$ for $X \in \{S, I\}$ and $i \neq j$, and the absence of interspecific competition implies $\partial F_i/\partial X_j = 0$ for all host classes $X$ and $i \neq j$. All terms and derivatives in the following subsections are evaluated at $p^*$.

##### S1.6.1 Response to increased excretion or removal of infectious propagules

**Response to increased degradation rate:** The responses to increases in the infectious propagule degradation rate are

$$\frac{\partial I_1^*}{\partial \delta} \propto (-1)^{3+5}\frac{M_{53}}{|J|} = \overbrace{\frac{\beta_1 S_1}{|J|}\left(\mu_2\frac{\partial f_1}{\partial S_1}\left[\frac{\partial f_2}{\partial S_2}-\beta_2 P\right]+P\beta_2\frac{\partial f_1}{\partial S_1}\frac{\partial f_2}{\partial I_2}\right)}^{P\to I_1 \,\&\, S_1\to S_1 \,\&\, S_2\to S_2 \,\&\, I_2\to I_2,\; P\to I_1 \,\&\, S_1\to S_1 \,\&\, S_2\to I_2\to S_2}$$
$$+ \overbrace{\frac{\beta_1\beta_2 P}{|J|}\left[\frac{\partial f_2}{\partial I_2}S_2\frac{\partial f_1}{\partial S_2}-\frac{\partial f_1}{\partial I_2}\left(S_1\frac{\partial f_2}{\partial S_1}+S_2\frac{\partial f_2}{\partial S_2}\right)\right]}^{P\to I_2\to S_2\to S_1\to I_1,\; P\to I_1 \,\&\, S_2\to I_2\to S_1\to S_2,\; P\to I_2\to S_1\to I_1 \,\&\, S_2\to S_2} - \overbrace{\frac{\partial f_1}{\partial S_2}\frac{\partial f_2}{\partial S_1}\frac{\beta_1 S_1\mu_2}{|J|}}^{P\to I_1 \,\&\, I_2\to I_2 \,\&\, S_2\to I_2\to S_2}, \tag{S42}$$

$$\frac{\partial S_1^*}{\partial \delta} \propto (-1)^{1+5}\frac{M_{13}}{|J|} = \overbrace{\frac{\mu_2}{|J|}\left(\frac{\partial f_2}{\partial S_2}-\beta_2 P\right)\left(\mu_1-\beta_1 S_1\frac{\partial f_1}{\partial I_1}\right)}^{P\to I_1\to S_1 \,\&\, S_2\to S_2 \,\&\, I_2\to I_2} - \overbrace{\frac{\beta_1 S_1\beta_2 P}{|J|}\left(\frac{\partial f_1}{\partial I_1}-\mu_1\right)\frac{\partial f_2}{\partial I_2}}^{P\to I_1\to S_1 \,\&\, S_2\to I_2\to S_2}$$
$$+\overbrace{\frac{\mu_1\beta_2 S_2}{|J|}\left(\frac{\partial f_2}{\partial S_2}-\beta_2 P\right)\frac{\partial f_1}{\partial I_2}+\frac{\beta_2 P}{|J|}\frac{\partial f_1}{\partial I_2}\left(\frac{\partial f_2}{\partial I_1}\beta_1 S_1-\mu_1\beta_2 S_2\right)}^{P\to I_2\to S_1 \,\&\, S_2\to S_2 \,\&\, I_1\to I_1,\; P\to I_1\to S_2\to I_2\to S_1}$$
$$+\overbrace{\frac{1}{|J|}\frac{\partial f_1}{\partial S_2}\left[\frac{\partial f_2}{\partial I_1}\mu_2\beta_1 S_1+\beta_2 S_2\mu_1\left(\frac{\partial f_2}{\partial I_2}-\mu_2\right)\right]}^{P\to I_1\to S_2\to S_1 \,\&\, I_2\to I_2,\; P\to I_2\to S_2\to S_1 \,\&\, I_1\to I_1}, \tag{S43}$$

and

$$\frac{\partial}{\partial \delta}\left(\frac{I_1^*}{N_1^*}\right) \propto \overbrace{\frac{P\beta_1^2 S_1^2}{\mu_1|J|}\frac{\partial f_1}{\partial I_1}}^{P\to I_1\to S_1}\overbrace{\left[P\beta_2\frac{\partial f_2}{\partial I_2}+\mu_2\left(\frac{\partial f_2}{\partial S_2}-P\beta_2\right)\right]}^{I_2\to S\to S_2,\; I_2\to I_2 \,\&\, S_2\to S_2}$$
$$+\overbrace{\frac{\beta_1\beta_2 S_1^2 P}{|J|}\frac{\partial f_2}{\partial I_2}\left(\frac{\partial f_1}{\partial S_1}-\beta_1 P\right)}^{P\to I_1 \,\&\, S_2\to I_2\to S_2 \,\&\, S_1\to S_1}+\overbrace{\frac{\beta_1\mu_2 S_1^2}{|J|}\left(\frac{\partial f_1}{\partial S_1}-\beta_1 P\right)\left(\frac{\partial f_2}{\partial S_2}-\beta_2 P\right)}^{P\to I_1 \,\&\, I_2\to I_2 \,\&\, S_2\to S_2 \,\&\, S_1\to S_1}$$
$$-\frac{\beta_1 S_1^2}{\mu_1|J|}\left(P\beta_2\frac{\partial f_1}{\partial I_2}+\mu_2\frac{\partial f_1}{\partial S_2}\right)\left(P\beta_1\frac{\partial f_2}{\partial I_1}+\mu_1\frac{\partial f_2}{\partial S_1}\right). \tag{S44}$$

Negative density dependence: In the absence of interspecific competition equation (S44) is negative. Equation (S44) becomes more positive with increased interspecific competition. Equation (S44) is expected to be negative for most systems. Equation (S44) can be positive under two scenarios: (1) interspecific competition is greater than intraspecific competition ($\partial F_i/\partial X_j$ much larger than $\partial F_i/\partial X_i$ for all host classes $X$ and $i \neq j$) and (2) interspecific competition between infected and susceptible individuals is greater than both intraspecific competition between infected and susceptible individuals ($\partial F_i/\partial I_j$ larger than $\partial F_i/\partial I_i$ for $i \neq j$) and interspecific competition between susceptible individuals ($\partial F_i/\partial I_j$ larger than $\partial F_i/\partial S_j$ for $i \neq j$).

Positive density dependence: All of the above predictions can be reversed if the infected classes for one or both hosts have sufficiently strong positive density dependence at equilibrium. Sufficiently strong positive density dependence requires $\partial F_i/\partial I_i$ is positive and large in magnitude.

**Response to increased uptake:** Because $\partial(dP/dt)/\partial\delta$ and $\partial(dP/dt)/\partial u_{ij}$ have the same sign, the signs of $\partial(I_1^*/N_1^*)/\partial u_{ij}$ and $\partial(I_1^*/N_1^*)/\partial\delta$ are the same.

**Response to increased excretion rates:** Because $\partial(dP/dt)/\partial\delta$ and $\partial(dP/dt)/\partial\chi_i$ have opposite signs, $\partial(I_1^*/N_1^*)/\partial\chi_i$ and $\partial(I_1^*/N_1^*)/\partial\delta$ have opposite signs.

##### S1.6.2 Response to increased mortality of the second host

The responses to increases in the mortality rate of $I_2$ are

$$\begin{aligned}\frac{\partial I_1^*}{\partial\mu_2} \propto (-1)^{4+3}\frac{M_{43}}{|J|} &= \overbrace{(\chi_2 - u_{I_2}P)\frac{\beta_1 S_1}{|J|}\frac{\partial f_1}{\partial S_1}\left(\frac{\partial f_2}{\partial S_2} - \beta_2 P\right)}^{I_2\to P\to I_1\ \&\ S_1\to S_1\ \&\ S_2\to S_2} + \overbrace{\frac{\beta_1 u_{I_2} S_1 P}{|J|}\frac{\partial f_1}{\partial S_1}\frac{\partial f_2}{\partial I_2}}^{I_2\to S_2\to P\to I_1\ \&\ S_1\to S_1} \\ &\overbrace{-(\chi_2 - u_{I_2}P)\frac{\beta_1}{|J|}\frac{\partial f_1}{\partial S_2}\left(PS_2\beta_2 + S_1\frac{\partial f_2}{\partial S_1}\right)}^{I_2\to P\to S_2\to S_1\to I_1,\ I_2\to P\to I_1\ \&\ S_1\to S_2\to S_1} - \overbrace{\frac{\beta_1 u_{S_2} S_1 P}{|J|}\frac{\partial f_1}{\partial I_2}\frac{\partial f_2}{\partial S_1}}^{I_2\to S_1\to S_2\to P\to I_1} \\ &+ \overbrace{\frac{\beta_1 P}{|J|}\frac{\partial f_1}{\partial S_2}\frac{\partial f_2}{\partial I_2}(U^* + \delta - S_1 u_{S_1})}^{I_2\to S_2\to S_1\to I_1\ \&\ P\to P} \\ &+ \overbrace{\frac{\beta_1\beta_2 P^2}{|J|}\frac{\partial f_1}{\partial I_2}(U^* + \delta - S_1 u_{S_1} - S_2 u_{I_2}) - \frac{\beta_1 P}{|J|}\frac{\partial f_1}{\partial I_2}\frac{\partial f_2}{\partial S_2}(U^* + \delta - S_1 u_{S_1})}^{I_2\to S_1\to I_1\ \&\ S_2\to S_2\ \&\ P\to P},\end{aligned} \tag{S45}$$

$$\begin{aligned}\frac{\partial S_1^*}{\partial\mu_2} \propto (-1)^{4+1}\frac{M_{41}}{|J|} &= \overbrace{(\chi_2 - u_{I_2}P)\frac{\beta_1 S_1}{|J|}\left(\frac{\partial f_2}{\partial S_2} - \beta_2 P\right)\left(\mu_1 - \frac{\partial f_1}{\partial S_1}\right)}^{I_2\to P\to I_1\to S_1\ \&\ S_2\to S_2} + \overbrace{\frac{u_{S_2} P\beta_1 S_1}{|J|}\frac{\partial f_2}{\partial I_2}\left(\mu_1 - \frac{\partial f_1}{\partial I_1}\right)}^{I_2\to S_2\to P\to I_1\to S_1} \\ &+ \overbrace{\frac{(\chi_2 - u_{I_2}P)}{|J|}\frac{\partial f_1}{\partial S_2}\left(\beta_1 S_1\frac{\partial f_2}{\partial I_1} - \mu_1\beta_2 S_2\right)}^{I_2\to P\to I_1\to S_2\to S_1,\ I_2\to P\to S_2\to S_1\ \&\ I_1\to I_1} - \overbrace{\frac{u_{S_2} P}{|J|}\frac{\partial f_1}{\partial I_2}\left(\mu_1\beta_2 S_2 - \beta_1 S_1\frac{\partial f_2}{\partial I_1}\right)}^{I_2\to S_1\ \&\ I_1\to I_1\ \&\ S_2\to P\to S_2,\ I_2\to S_1\ \&\ S_2\to P\to I_1\to S_2} \\ &+ \overbrace{\frac{\mu_1(U^* + \delta) - (\chi_1 - u_{I_1}P)\beta_1 S_1}{|J|}\left[\frac{\partial f_1}{\partial S_2}\frac{\partial f_2}{\partial I_2} - \left(\frac{\partial f_2}{\partial I_2} - \beta_2 P\right)\frac{\partial f_1}{\partial I_2}\right]}^{I_2\to S_2\to S_1\ \&\ I_1\to I_1\ \&\ P\to P,\ I_2\to S_1\ \&\ S_2\to S_2\ \&\ I_1\to I_1\ \&\ P\to P},\end{aligned} \tag{S46}$$

and

$$
\begin{aligned}
\frac{\partial}{\partial \mu_2}\left(\frac{I_1^*}{N_1^*}\right) \propto\ & \overbrace{\frac{\beta_1 S_1^2}{\mu_1|J|}(\chi_2 - u_{S_2}P)}^{I_2\to P\to I_1}\overbrace{\left(\frac{\partial f_2}{\partial S_2} - \beta_2 P\right)}^{S_2\to S_2}\overbrace{\left[P\beta_1\frac{\partial f_1}{\partial I_1} + \mu_1\left(\frac{\partial f_1}{\partial S_1} - \beta_1 P\right)\right]}^{S_1\to I_1\to S_1,\ S_1\to S_1\ \&\ I_1\to I_2} \\
& + \overbrace{\frac{\beta_1 u_{S_2} S_1^2 P}{\mu_1|J|}\frac{\partial f_2}{\partial I_2}}^{I_2\to S_2\to P\to I_1}\overbrace{\left[P\beta_1\frac{\partial f_1}{\partial I_1} + \mu_1\left(\frac{\partial f_1}{\partial S_1} - \beta_1 P\right)\right]}^{S_1\to I_1\to S_1,\ S_1\to S_1\ \&\ I_1\to I_2} \\
& - \frac{\beta_1 S_1^2}{\mu_1|J|}(\chi_2 - u_{S_2}P)\frac{\partial f_1}{\partial S_2}\left(\beta_1 P\frac{\partial f_2}{\partial I_1} + \mu_1\frac{\partial f_2}{\partial S_1}\right) - \frac{\beta_1 u_{S_2} S_1^2 P}{\mu_1|J|}\frac{\partial f_1}{\partial I_2}\left(\beta_1 P\frac{\partial f_2}{\partial I_1} + \mu_1\frac{\partial f_2}{\partial S_1}\right) \\
& + \frac{S_1 P}{|J|}\left[\frac{\partial f_1}{\partial I_2}\left(\beta_2 P - \frac{\partial f_2}{\partial S_2}\right) + \frac{\partial f_1}{\partial S_2}\frac{\partial f_2}{\partial I_2}\right](\chi_1 I_1 - u_{I_1} I_1 P - u_{S_1} S_1 P)\,.
\end{aligned}
\tag{S47}
$$

<u>Negative density dependence:</u> In the absence of interspecific competition, only the first two lines of equation (S47) are nonzero. The sign of the first line is $-(\chi_2 - u_{I_2}P)$ and the second line is negative. This means increases in $\mu_2$ cause $I_1/N_1$ to decrease unless infected individuals in population 2 are large sinks, i.e., $\chi_2 - u_{I_2}P$ negative and large in magnitude. The terms on line three of equation (S47) have the opposite sign of those on lines 1 and 2. This means that increased interspecific host competition can cause the sign of equation (S47) to switch.

Equation (S47) can have the opposite signs under two scenarios: (1) interspecific competition is greater than intraspecific competition ($\partial F_i/\partial X_j$ much larger than $\partial F_i/\partial X_i$ for all host classes $X$ and $i \neq j$) and (2) interspecific competition between infected and susceptible individuals is greater than both intraspecific competition between infected and susceptible individuals ($\partial F_i/\partial I_j$ larger than $\partial F_i/\partial I_i$ for $i \neq j$) and interspecific competition between susceptible individuals ($\partial F_i/\partial I_j$ larger than $\partial F_i/\partial S_j$ for $i \neq j$).

<u>Positive density dependence:</u> All of the above predictions can be reversed if the infected classes for one or both hosts have sufficiently strong positive density dependence at equilibrium.

##### S1.6.3 Response to increased infection rates of the second host

The responses to increases in the infection rate of the second host are defined by

$$
\begin{aligned}
\frac{\partial I_1^*}{\partial \beta_2} =\ & \overbrace{\frac{-\beta_1 u_{S_2} S_1 P}{|J|}\frac{\partial f_1}{\partial S_1}\left(\frac{\partial f_2}{\partial I_2} - \mu_2\right)}^{I_2\to S_2\to P\to I_1\ \&\ S_1\to S_1} - \overbrace{(\chi_2 - u_{I_2}P)\frac{\beta_1 S_1}{|J|}\left(\frac{\partial f_1}{\partial S_1}\frac{\partial f_2}{\partial S_2} - \frac{\partial f_1}{\partial S_2}\frac{\partial f_2}{\partial S_1}\right)}^{I_2\to P\to I_1\ \&\ S_1\to S_1\ \&\ S_2\to S_2,\ I_2\to P\to I_1\ \&\ S_1\to S_2\to S_1} \\
& + \overbrace{\frac{\beta_1 P}{|J|}(U^* + \delta - u_{S_1}S_1)\left(\frac{\partial f_1}{\partial I_2}\frac{\partial f_2}{\partial S_2} - \frac{\partial f_1}{\partial S_2}\frac{\partial f_2}{\partial I_2} + \mu_2\frac{\partial f_1}{\partial S_2}\right)}^{I_2\to S_1\to I_1\ \&\ P\to P\ \&\ S_2\to S_2,\ I_2\to S_2\to S_1\to I_1\ \&\ P\to P} + \overbrace{\frac{\beta_1 u_{I_2} S_1 P}{|J|}\frac{\partial f_1}{\partial I_2}\frac{\partial f_2}{\partial S_1}}^{I_2\to S_1\to S_2\to P\to I_1},
\end{aligned}
\tag{S48}
$$

$$
\frac{\partial S_1^*}{\partial \beta_2} = -\frac{\partial}{\partial \beta_2}\left(\frac{dS_2}{dt}\right)\frac{(-1)^{2+1}M_{21}}{|J|} - \frac{\partial}{\partial \beta_2}\left(\frac{dI_2}{dt}\right)\frac{(-1)^{4+1}M_{41}}{|J|} = \frac{S_2 P}{|J|}(M_{41} - M_{21}), \tag{S49}
$$

where $M_{ij}$ is the $i,j$ minor of $J$, and

$$
\begin{aligned}
\frac{\partial}{\partial\beta_2}\left(\frac{I_1^*}{N_1^*}\right) \propto &- \overbrace{\frac{(\chi_2 - u_{I_2}P)}{|J|}\left[\beta_1 P\frac{\partial f_1}{\partial I_1} + \mu_1\left(\frac{\partial f_1}{\partial S_1} - \beta_1 P\right)\right]}^{I_2\to P\to I_1\to S_1,\, I_2\to P\to I_1}\overbrace{\frac{\partial f_2}{\partial S_2}}^{S_2\to S_2} \\
&- \overbrace{\frac{u_{S_2}P}{|J|}\left(\frac{\partial f_2}{\partial I_2} - \mu_2\right)}^{I_2\to S_2\to P}\overbrace{\left[\beta_1 P\frac{\partial f_1}{\partial I_1} + \mu_1\left(\frac{\partial f_1}{\partial S_1} - \beta_1 P\right)\right]}^{P\to I_1\to S_1,\, P\to S_1,\, P\to I_1} \\
&+ \frac{(\chi_2 - u_{I_2}P)}{|J|}\frac{\partial f_1}{\partial S_2}\left(\beta_1 P\frac{\partial f_2}{\partial I_1} + \mu_2\frac{\partial f_2}{\partial S_1}\right) - (\chi_1 - u_{I_1}P)\frac{\beta_1 P}{|J|}\frac{\partial f_1}{\partial S_2}\left(\frac{\partial f_2}{\partial I_2} - \mu_2\right) \\
&+ \frac{\mu_1 u_{S_1}P}{|J|}\frac{\partial f_1}{\partial S_2}\left(\frac{\partial f_2}{\partial I_2} - \mu_2\right) + \frac{u_{S_2}P}{|J|}\frac{\partial f_1}{\partial I_2}\left(\beta_1 P\frac{\partial f_2}{\partial I_1} + \mu_1\frac{\partial f_2}{\partial S_1}\right) \\
&+ (\chi_1 I_1 - u_{I_1}I_1 P - u_{S_1}S_1 P)\frac{\mu_1 P}{\beta_1 S_1 |J|}\frac{\partial f_1}{\partial I_2}\frac{\partial f_2}{\partial S_2}.
\end{aligned}
\tag{S50}
$$

Negative density dependence: In the absence of interspecific competition, only the first and second lines of equation (S50) are nonzero. The second line is positive and the sign of the first line is $\chi_2 - u_{I_2}P$. The term on the first line is negative and larger in magnitude than the term on the second line when $\chi_2 - u_{I_2}P < 0$ and either $u_{I_2}$ is sufficiently greater than $u_{S_2}$. This means that if there is no interspecific competition, then equation (S50) is positive unless the second host is a large sink ($\chi_2 - u_{I_2}P < 0$) and infected individuals of the second host have larger uptake rates than susceptible individuals ($u_{I_2} > u_{S_2}$).

The first term on line three and the first line four of equation (S50) have the opposite signs of the first and second lines of equation (S50). The remaining terms can make equation (S50) more positive or negative depending on the sign of $\chi_1 - u_{I_1}P$. This means that increased interspecific competition can change the sign of equation (S50). We expect increased interspecific competition to cause equation (S50) to change sign when (1) interspecific competition is much greater than intraspecific competition ($\partial F_i/\partial X_j$ much larger than $\partial F_i/\partial X_i$ for all host classes $X$ and $i \neq j$), (2) interspecific competition between infected and susceptible individuals is greater than both intraspecific competition between infected and susceptible individuals ($\partial F_i/\partial I_j$ larger than $\partial F_i/\partial I_i$ for $i \neq j$) and interspecific competition between susceptible individuals ($\partial F_i/\partial I_j$ larger than $\partial F_i/\partial S_j$ for $i \neq j$), or (3) the second host is a large source and the focal host is a large sink ($\chi_1 - u_{S_1}P$ negative and large in magnitude).

Positive density dependence: All of the above predictions can be reversed if the infected classes for one or both hosts have sufficiently strong positive density dependence at equilibrium.

##### S1.6.4 Response to increased intraspecific competition in the second host

Let $\alpha_{22}$ be any parameter that negatively affects the reproduction rate of the second host. Mathematically, we assume $\partial f_2/\partial\alpha_{22} < 0$, which implies $\partial(dS_2/dt)/\partial\alpha_{22} < 0$. For example, a model with Lotka-Volterra competition, $f_2 = r_2(S_2 + c_2 I_2)[1 - \alpha_{22}(S_1 + e_{21}I_1) - \alpha_{22}(S_2 + e_{22}I_2)]$, satisfies these conditions. Note that our results apply to changes in any parameter that negatively affects the growth rate of the second host. For the Lotka-Volterra example, our results apply to changes in $\alpha_{22}$ as well as changes in $\alpha_{21}$, $e_{21}$, and $e_{22}$.

The responses to the second host experiencing increased intraspecific competition are

$$\begin{aligned}\frac{\partial I_1^*}{\partial \alpha_{22}} \propto \frac{(-1)^{2+3}M_{23}}{|J|} = & \overbrace{-\frac{\beta_1\mu_2}{|J|S_2}\left(S_1\frac{\partial f_1}{\partial S_1}+S_2\frac{\partial f_1}{\partial S_2}\right)(\chi_2 I_2 - u_{S_2}S_2P - u_{I_2}I_2P)}^{S_2\to I_2\to P\to I_1\ \&\ S_1\to S_1,\ S_2\to P\to I_1\ \&\ I_2\to P\to I_2} \\ & + \overbrace{\frac{\beta_1 P}{|J|}\left[\mu_2\frac{\partial f_1}{\partial S_2}+\frac{\partial f_1}{\partial I_2}\beta_2 P\right](U^*+\delta-u_{S_1}S_1-u_{S_2}S_2)}^{S_2\to S_1\to I_1\ \&\ P\to P\ \&\ I_2\to I_2,\ S_2\to I_2\to S_1\to I_1\ \&\ P\to P}\end{aligned} \quad \text{(S51)}$$

$$\begin{aligned}\frac{\partial S_1^*}{\partial \alpha_{22}} \propto \frac{(-1)^{2+1}M_{21}}{|J|} = & \overbrace{(\chi_2-u_{I_2}P)\frac{\beta_1 S_1\beta_2 P}{|J|}\left(\frac{\partial f_1}{\partial I_1}-\mu_1\right)}^{S_2\to I_2\to P\to I_1\to S_1} - \overbrace{\frac{u_{S_2}P\mu_2\beta_1 S_1}{|J|}\left(\frac{\partial f_1}{\partial I_1}-\mu_1\right)}^{S_2\to P\to I_1\to S_1\ \&\ I_2\to I_2} \\ & + \overbrace{\frac{\mu_1\beta_2 P}{|J|}\frac{\partial f_1}{\partial I_2}(U^*+\delta-u_{S_2}S_2)}^{S_2\to I_2\to S_1\ \&\ I_1\to I_1\ \&\ P\to P} + \overbrace{\frac{\mu_1\mu_2}{|J|}\frac{\partial f_1}{\partial S_2}(U^*+\delta)}^{S_2\to S_1\ \&\ I_1\to I_1\ \&\ I_2\to I_2\ \&\ P\to P} \\ & \overbrace{-\frac{\partial f_1}{\partial S_2}\left[(\chi_1-u_{I_1}P)\mu_2\beta_1 S_1+(\chi_2-u_{I_2}P)\mu_1\beta_2 S_2\right]}^{S_2\to S_1\ \&\ I_2\to I_2\ \&\ I_1\to P\to I_1,\ S_2\to S_1\ \&\ I_1\to I_1\ \&\ I_2\to P\to I_2},\end{aligned} \quad \text{(S52)}$$

and

$$\begin{aligned}\frac{\partial}{\partial \alpha_{22}}\left(\frac{I_1^*}{N_1^*}\right) \propto & -\frac{\beta_1\mu_2 S_1^2 P}{\beta_2 S_2\mu_1|J|}\left[\chi_2 I_2 - u_{I_2}I_2P - u_{S_2}S_2P\right]\left[\beta_1 P\frac{\partial f_1}{\partial I_1}+\mu_1\left(\frac{\partial f_1}{\partial S_1}-\beta_1 P\right)\right] \\ & + \frac{S_1 P}{|J|}\left[\chi_1 I_1 - u_{I_1}I_1P - u_{S_1}S_1P\right]\left(\beta_2 P\frac{\partial f_1}{\partial I_2}+\mu_2\frac{\partial f_1}{\partial S_2}\right).\end{aligned} \quad \text{(S53)}$$

Negative density dependence: The sign of the first line of equation (S53) is $-(\chi_2 I_2 - u_{I_2}I_2P - u_{S_2}S_2P)$ and the sign of the second line is $\chi_1 I_1 - u_{I_1}I_1P - u_{S_1}S_1P$. Equation (S53) is negative when $\chi_1 - u_{I_1}P$ is negative and sufficiently large or $\chi_2 - u_{I_2}P$ is positive and sufficiently large. At equilibrium, one of $\chi_1 - u_{I_1}P^*$ or $\chi_2 - u_{I_2}P^*$ must be positive. This means equation (S53) will be positive unless the second host is a sufficiently large source.

Positive density dependence: All of the above predictions can be reversed if the infected class for the focal host has sufficiently strong positive density dependence at equilibrium.

###### S1.6.5 Response to increased interspecific competitive ability of the second host

Let $\alpha_{12}$ be any parameter that negatively affects the reproduction rate of the focal host . Mathematically, we assume $\partial f_1/\partial\alpha_{12} < 0$, which implies $\partial(dS_1/dt)/\partial\alpha_{12} < 0$. For example, a model with Lotka-Volterra competition, i.e., $f_1 = r_1(S_1+b_1I_1)[1-\alpha_{11}(S_1+c_1I_1)-\alpha_{12}(S_2+c_2I_2)]$, satisfies these conditions. Note that our results apply to any parameter that negatively affects the growth rate of the focal host . For the Lotka-Volterra example, our results apply to changes in $\alpha_{12}$ as well as changes in $\alpha_{11}$, $c_1$, and $c_2$.

The responses to the focal host experiencing increased interspecific competition are defined by

$$\begin{aligned}\frac{\partial I_1^*}{\partial \alpha_{12}} \propto \frac{(-1)^{1+3}M_{13}}{|J|} &= \overbrace{(\chi_2 I_2 - u_{S_2}S_2P - u_{I_2}I_2P)\frac{\beta_1\mu_2P}{\beta_2S_2|J|}\left(S_2\frac{\partial f_2}{\partial S_2}+S_1\frac{\partial f_2}{\partial S_1}\right)}^{S_1\to I_1\ \&\ S_2\to I_2\to P\to S_2,\ S_1\to S_2\to I_2\to P\to I_1} \\ &\overbrace{-\frac{\beta_1P}{|J|}\left[\left(\frac{\partial f_2}{\partial S_2}-\beta_2P\right)\mu_2+\frac{\partial f_2}{\partial I_2}\beta_2P\right](U^*+\delta-u_{S_1}S_1-u_{S_2}S_2)}^{S_1\to I_1\ \&\ S_2\to S_2\ \&\ I_2\to I_2\ \&\ P\to P,\ S_1\to I_1\ \&\ S_2\to I_2\to S_2\ \&\ P\to P},\end{aligned} \tag{S54}$$

$$\begin{aligned}\frac{\partial S_1^*}{\partial \alpha_{12}} \propto \frac{(-1)^{1+1}M_{11}}{|J|} &= \overbrace{(\chi_2-u_{I_2}P)\frac{\mu_1\beta_2S_2}{|J|}\frac{\partial f_2}{\partial S_2}}^{S_2\to S_2\ \&\ I_2\to P\to I_2} + \overbrace{(\chi_1-u_{I_1}P)\frac{\beta_1S_1}{|J|}\left(\mu_2\frac{\partial f_2}{\partial S_2}+\beta_2S_2\frac{\partial f_2}{\partial I_2}-\beta_2\mu_2P\right)}^{I_1\to P\to I_1\ \&\ S_2\to I_2\to S_2} \\ &- \overbrace{\frac{\mu_1}{|J|}(U^*+\delta)\left(\mu_2\frac{\partial f_2}{\partial S_2}+\beta_2P\frac{\partial f_2}{\partial I_2}\right)}^{P\to P\ \&\ I_1\to I_1\ \&\ S_2\to I_2\to S_2} + \overbrace{\frac{\mu_1\mu_2}{|J|}(U^*+\delta-u_{S_2}S_2)}^{I_1\to I_1\ \&\ I_2\to I_2\ \&\ P\to P} \\ &- \overbrace{(\chi_2-u_{I_2}P)\frac{\beta_1S_1\beta_2P}{|J|}\frac{\partial f_2}{\partial I_1}}^{I_1\to S_2\to I_2\to P\to I_1} + \overbrace{\frac{\mu_2u_{S_2}P\beta_1S_1}{|J|}\frac{\partial f_2}{\partial I_1}}^{I_1\to S_2\to P\to I_1\ \&\ I_2\to I_2},\end{aligned} \tag{S55}$$

and

$$\begin{aligned}\frac{\partial}{\partial \alpha_{12}}\left(\frac{I_1^*}{N_1^*}\right) \propto &\frac{-S_1P}{|J|}\left[\chi_1I_1-u_{I_1}I_1P-u_{S_1}S_1P\right]\left[\beta_2P\frac{\partial f_2}{\partial I_2}+\mu_2\left(\frac{\partial f_2}{\partial S_2}-\beta_2P\right)\right] \\ &+\frac{\beta_1\mu_2S_1^2P}{\beta_2\mu_1S_2|J|}\left[\chi_2I_2-u_{I_2}I_2P-u_{S_2}S_2P\right]\left(\beta_1P\frac{\partial f_2}{\partial I_1}+\mu_1\frac{\partial f_2}{\partial S_1}\right).\end{aligned} \tag{S56}$$

Negative density dependence: The sign of the first line of equation (S56) is determined by $-(\chi_1I_1 - u_{I_1}I_1P - u_{S_1}S_1P)$ and the sign of the second line of equation (S56) is determined by $\chi_2I_2 - u_{I_2}I_2P - u_{S_2}S_2P$. Equation (S56) is negative when $\chi_1I_1 - u_{I_1}I_1P - u_{S_1}S_1P$ is positive and sufficiently large or $\chi_2I_2 - u_{I_2}I_2P - u_{S_2}S_2P$ is negative and sufficiently large. At equilibrium, one of $\chi_1 - u_{I_1}P^*$ or $\chi_2 - u_{I_2}P^*$ must be positive. This means equation (S56) will be negative unless the the second host is a sufficiently large source.

Positive density dependence: All of the above predictions can be reversed if the infected class of the second host has sufficiently strong positive density dependence at equilibrium.

### S2 Analysis of SIRS Environmental Transmission Model

#### S2.1 SIRS Model

To allow for recovery from infection and loss of immunity, we extend the SI model to include a recovered class ($R$). The multi-species model is

$$\begin{aligned}
\frac{dS_i}{dt} &= \underbrace{F_i(\cdot)}_{\text{growth \& competition}} - \underbrace{\beta_i S_i P}_{\text{infection}} + \underbrace{\gamma_i R_i}_{\text{loss of immunity}} \\
\frac{dI_i}{dt} &= \underbrace{\beta_i S_i P}_{\text{infection}} - \underbrace{\mu_i I_i}_{\text{mortality}} - \underbrace{\nu_i I_i}_{\text{recovery}} \\
\frac{dR}{dt} &= \underbrace{\nu_i I_i}_{\text{recovery}} - \underbrace{\gamma_i R_i}_{\text{loss of immunity}} - \underbrace{m_i R_i}_{\text{mortality}} \\
\frac{dP}{dt} &= \underbrace{\sum_i \chi_{i1} I_i}_{\text{propagule excretion}} \underbrace{- \sum_i (u_{I_1} S_i + u_{I_2} I_i + u_{R_i} R_i) P}_{\text{propagule uptake}} \underbrace{-\delta P}_{\text{degradation}} .
\end{aligned} \tag{S57}$$

where $(\cdot)$ is shorthand for $(S_1, S_2, I_1, I_2, R_1, R_2)$ and for host $i$, $\nu_i$ is the recovery rate; $\gamma_i$ is the rate at which recovered individuals lose immunity; $u_{R_i}$ is loss rates of infectious propagules due to uptake by recovered individuals; and all other terms are defined as in model (1) from the main text. We assume the reproductive rates can be written as $F_i(\cdot) = S_i f_{S_i}(\cdot) + I_i f_{I_i}(\cdot) + R_i f_{R_i}(\cdot)$ where $X_i f_{X_i}$ is the reproductive rate of individuals in class $X_i$. We assume $\partial f_{X_i}/\partial Y_j < 0$ for all $i, j$ and any host classes $X$ and $Y_1$ because of intraspecific and interspecific competition. We assume the mortality rate for recovered individuals is the same as susceptible individuals ($m_i$). As with the SI model, to simplify the notation we absorb the natural mortality rates into the reproduction term for susceptible individuals ($f_{S_i}$) and mortality rate for the infected individuals ($\mu_i$).

We assume the multi-species model (S57) has a unique stable endemic equilibrium coexistence equilibrium, $p^* = (S_1^*, S_2^*, I_1^*, I_2^*, R_1^*, R_2^*, P^*)$. Removing the $dS_2/dt$, $dI_2/dt$ and $dR_2/dt$ equations from model (S57) and setting $S_2 = I_2 = R_2 = 0$ yields a single-species model where only the focal host is present. We assume the single-species model has a unique stable endemic equilibrium $\hat{p} = (\hat{S}_1, \hat{I}_1, \hat{R}_1, \hat{P})$. Total density is denoted $N_i = S_i + I_i + R_i$ and the total per spore infectious propagule uptake rate is denoted $U = \sum_i (u_{I_1} S_i + u_{I_2} I_i + u_{R_i} R_i)P$. The equilibrium values are denoted $N_i^*$, $\hat{N}_i$, $U^*$, and $\hat{U}$. Our definitions for higher and lower competence and sinks and sources are unchanged; see section 1.1.

Imposing particular conditions on model (S57) allows us to recover special cases of interest. Setting $\gamma_i = 0$ accounts for pathogens where immunity is life-long. In the limit where $m_i = 0$ and $\gamma \to \infty$, model (S57) reduces to an SIS model where there is recovery from infection but no immunity. Setting $\nu_i = 0$ turns the SIRS model into the SI model (1) from the main text. The SIRS model reduces to a FDDT or a DDDT system under the same conditions as the SI model. In general, the SIRS model reduces to a direct transmission model when there is a separation of time scales between the infectious propagule dynamics and the dynamics of the host classes.

Negative density dependence in the SIRS model (S57) implies that the rate of change of the susceptible population decreases with an increase in the density of any host class. Negative density dependence occurs when $\partial F_i/\partial S_i < 0$, $\partial F_i/\partial I_i < 0$, and $\gamma_i + \partial F_i/\partial R_i < 0$. Positive density dependence is defined by the reverse inequalities. We expect negative density dependence to be present in most systems. However, positive density dependence can arise if (i) the pathogen suppresses host densities to very low values or (ii) recovered individuals are weaker intraspecific competitors than susceptible individuals ($\partial F_i/\partial R_i$ smaller in magnitude than $\partial F_i/\partial S_i$) and loss of immunity rates are large ($\gamma_i$ large).

##### S2.1.1 single-species equilibrium

The Jacobian evaluated at single-species equilibrium, $\hat{p}$, is

$$\hat{J} = \left.\begin{pmatrix} \frac{\partial f_1}{\partial S_1} - \beta_1 P & \frac{\partial f_1}{\partial I_1} & \frac{\partial f_1}{\partial R_1} + \gamma_1 & -\beta_1 S_1 \\ \beta_1 P & -\mu_1 0 - \nu_1 & 0 & \beta_1 S_1 \\ 0 & \nu_1 & -\gamma_1 - m_1 & 0 \\ -u_{S_1} P & \chi_1 - u_{I_1} P & -u_{R_1} P & -\hat{U} - \delta \end{pmatrix}\right|_{\hat{p}} \tag{S58}$$

with sign structure

$$\begin{pmatrix} - & \pm & \pm & - \\ + & - & 0 & + \\ 0 & + & 0 & - \\ - & + & - & - \end{pmatrix} \tag{S59}$$

where $\pm$ means the entry can have a positive or negative sign. The entries in the first row represent the combined effects of competition and reproduction of each host class ($\hat{J}_{11}$, $\hat{J}_{12}$, $\hat{J}_{13}$), infection ($\hat{J}_{11}$, $\hat{J}_{14}$), and recovery ($\hat{J}_{13}$); the entries in the second row represent the effects of infection ($\hat{J}_{21}$, $\hat{J}_{24}$) and mortality and recovery ($\hat{J}_{22}$); the entries in the third row represent the effects of recovery ($\hat{J}_{32}$) and the negative effects of loss of immunity and mortality of recovered individuals ($\hat{J}_{33}$); and the entries in the fourth row represent the negative effects due to uptake by susceptible ($\hat{J}_{31}$) and recovered ($\hat{J}_{33}$) hosts and degradation ($\hat{J}_{33}$) and the combined effect of propagule release and uptake by infected individuals ($\hat{J}_{32}$). Stability of $\hat{p}$ implies $|\hat{J}| > 0$.

Entry $\hat{J}_{11}$ is assumed to be negative for the following reason. Computing the $\hat{J}_{11}$ entry yields

$$\begin{aligned} \hat{J}_{11} &= f_{1S} - \beta_1 P + S_1 \frac{\partial f_{1S}}{\partial S_1} + I_1 \frac{\partial f_{1I}}{\partial S_1}\bigg|_{\hat{p}} \\ &= -\left(\frac{I_1 f_{1I}(\cdot)}{S_1} + \frac{R_1 f_{1R}(\cdot)}{S_1} + \frac{\gamma_1 R_1}{S_1}\right) + S_1 \frac{\partial f_{1S}}{\partial S_1} + I_1 \frac{\partial f_{1I}}{\partial S_1}\bigg|_{\hat{p}} < 0 \end{aligned} \tag{S60}$$

where the last equality follows from the fact that at equilibrium $dS_1/dt = 0$, which means $f_{1S}(\cdot) - \beta_1 \hat{P} = -[\hat{I}_1 f_{1I}(\cdot) + \hat{R}_1 f_{1R}(\cdot) + \gamma_1 R_1]/\hat{S}_1$. While $\hat{J}_{11}$ is negative, entries $\hat{J}_{12}$ and $\hat{J}_{13}$ can have either sign because $\partial(X_1 f_{1X})/\partial X_1$ can be positive or negative depending on the densities of the host classes. Whenever we assume negative density dependence, all $\pm$ entries of the Jacobian are negative. Positive density dependence allows for one or more of those entries to be positive.

##### S2.1.2 multi-species equilibrium

The Jacobian evaluated at the multi-species equilibrium, $p^*$, is

$$J = \begin{pmatrix} \frac{\partial f_1}{\partial S_1} - \beta_1 P & \frac{\partial f_1}{\partial S_2} & \frac{\partial f_1}{\partial I_1} & \frac{\partial f_1}{\partial I_2} & \frac{\partial f_1}{\partial R_1} + \gamma_1 & \frac{\partial f_1}{\partial R_2} & -\beta_1 S_1 \\ \frac{\partial f_2}{\partial S_1} & \frac{\partial f_2}{\partial S_2} - \beta_2 P & \frac{\partial f_2}{\partial I_1} & \frac{\partial f_2}{\partial I_2} & \frac{\partial f_2}{\partial R_1} & \frac{\partial f_2}{\partial R_2} + \gamma_2 & -\beta_2 S_2 \\ \beta_1 P & 0 & -\mu_1 - \nu_1 & 0 & 0 & 0 & \beta_1 S_1 \\ 0 & \beta_2 P & 0 & -\mu_2 - \nu_2 & 0 & 0 & \beta_2 S_2 \\ 0 & 0 & \nu_1 & 0 & -\gamma_1 - m_1 & 0 & 0 \\ 0 & 0 & 0 & \nu_2 & 0 & -\gamma_2 - m_2 & 0 \\ -u_{S_1} P & -u_{S_2} P & \chi_1 - u_{I_1} P & \chi_2 - u_{I_2} P & -u_{R_1} P & -u_{R_2} P & -U^* - \delta \end{pmatrix} \tag{S61}$$

with sign structure

$$\begin{pmatrix} - & - & \pm & - & \pm & - & - \\ - & \pm & - & \pm & - & \pm & - \\ + & 0 & - & 0 & 0 & 0 & + \\ 0 & + & 0 & - & 0 & 0 & + \\ 0 & 0 & + & 0 & - & 0 & 0 \\ 0 & 0 & 0 & + & 0 & - & 0 \\ - & - & \pm & \pm & - & - & - \end{pmatrix} \tag{S62}$$

where $\pm$ means the entry can have a positive or negative sign. The entries in the first and second rows represent the combined effects of intraspecific competition and reproduction of each host class ($J_{11}, J_{13}, J_{15}, J_{22}, J_{24}, J_{16}$), interspecific competition ($J_{12}, J_{14}, J_{16}, J_{21}, J_{23}, J_{25}$), infection ($J_{11}, J_{17}, J_{22}, J_{27}$), and loss of immunity ($J_{15}, J_{26}$); the entries in the third and fourth rows represent the effects of infection ($J_{31}$, $J_{37}$, $J_{42}$, $J_{47}$) and mortality and recovery ($J_{33}$, $J_{44}$); the entries in the fifth and sixth rows represent the effects of recovery ($J_{53}$, $J_{64}$) and mortality and loss of immunity ($J_{55}$, $J_{66}$); and the entries in the fifth row represent the negative effects due to uptake by susceptible and recovered individuals ($J_{71}$, $J_{72}$, $J_{75}$, $J_{76}$) and degradation ($J_{77}$) and the combined effects of propagule release and uptake by infected individuals ($J_{73}$, $J_{74}$). Stability of $p^*$ implies $|J| < 0$.

The reason why entries $J_{11}$ and $J_{22}$ are negative and entries $J_{13}$, $J_{15}$, $J_{24}$, and $J_{26}$ can be either sign is the same as for the single-species model. Whenever we assume negative density dependence, all $\pm$ entries in the Jacobian are negative.

#### S2.2 Response of the equilibrium prevalence to changes in the transmission mode

The following identifies how converting the environmental transmission (ET) model (S57) from a form that behaves like a frequency-dependent direct transmission (FDDT) model to a form that behaves like a density-dependent direct transmission (DDDT) model affects disease prevalence in the focal host, using a change of parameters that holds the either the multi-species equilibrium densities ($p^*$) constant or single-species equilibrium densities ($\hat{p}$) constant. The discussion about changes of parameters (section section S1.6.1) and factors affecting the sign of $U^* - \hat{U}$ (section 1.6.2) for the SI model apply directly to the SIRS model. The only minor difference is that $U$ includes uptake by recovered individuals ($u_{R_i} R_i$).

##### S2.2.1 Parameter transformation that holds the multi-species equilibrium constant

We use a change of parameters that holds the multi-species equilibrium ($p^*$) of the SIRS model (S57) constant. The change of parameters is $f(q) = 1 + \frac{\delta}{U^*} - \frac{\delta}{U^*} q$ for $0 \leq q \leq 1 + U^*/\delta$; see section S1.6.1 for details.

**Effect on single-species equilibrium densities:** Following the steps in Section S1.6.3, the effects of varying $q$ on the density and proportion of infected individuals at the single-species equilibrium are

$$\frac{\partial \hat{I}_1}{\partial q} = -\frac{\partial}{\partial q}\left(\frac{dP}{dt}\right)\frac{(-1)^{2+4} M_{42}}{|\hat{J}|} \propto (U^* - \hat{U})\frac{M_{42}}{|\hat{J}|} = (U^* - \hat{U})\frac{(\gamma_1 + m_1)\beta_1 \hat{S}_1}{|\hat{J}|}\frac{\partial f_1}{\partial S_1} \quad \text{(S63)}$$

and

$$\frac{\partial}{\partial q}\left(\frac{\hat{I}_1}{\hat{S}_1 + \hat{I}_1 + \hat{R}_1}\right) \propto (U^* - \hat{U})\frac{\partial}{\partial \delta}\left(\frac{\hat{I}_1}{\hat{S}_1 + \hat{I}_1 + \hat{R}_1}\right). \quad \text{(S64)}$$

where $|\hat{J}| > 0$, $M_{42}$ is a minor of matrix (S58), and all terms are evaluated at $\hat{p}$.

Negative density dependence: The sign of equation (S64) is determined by $-(U^* - \hat{U})$. If $U^* - \hat{U} > 0$, then $I_1^*/N_1^* - \hat{I}_1/\hat{N}_1$ is more negative under FDDT than DDDT. This means the proportion of infected individuals in the focal host decreases more (or increases less) under FDDT than DDDT when a second host is added. In contrast, if $U^* - \hat{U} < 0$, then $I_1^*/N_1^* - \hat{I}_1/\hat{N}_1$ is more positive under FDDT than DDDT. This means the proportion of infected individuals in the focal host decreases more (or increases less) under DDDT than FDDT when a second host is added.

*Biological Interpretation:* The above predictions are identical to those for the SI model. In particular, when a second host is added, the focal host infection prevalence decreases more (or increases less) under FDDT than DDDT when (D1) interspecific competition is weaker, (D2) the second host is a weaker intraspecific competitor, and (D3) the second host a lower competence host. Focal host infection prevalence decreases more (or increases less) under DDDT than FDDT when (D4) interspecific competition is stronger, (D5) the second host is a stronger intraspecific competitor, and (D6) the second host is a higher competence host.

Positive density dependence of the focal host at equilibrium: If the focal host has positive density dependence at the single-species equilibrium, then the above predictions are reversed.

###### S2.2.2 Parameter transformation that holds the single-species equilibrium constant

We use a change of parameters that holds the single-species equilibrium ($\hat{p}$) of the SIRS model (S57) constant. The change of parameters is $f(q) = 1 + \frac{\delta}{\hat{U}} - \frac{\delta}{\hat{U}} q$ for $0 \leq q \leq 1 + \hat{U}/\delta$; see section S1.6.1 for details.

**Effect on multi-species equilibrium densities:** Following the steps in Section S1.5.4, the effects of varying $q$ on the density and proportion of infected individuals at the multi-species equilibrium is

$$\frac{\partial I_1^*}{\partial q} = -\frac{\partial}{\partial q}\left(\frac{dP}{dt}\right)\frac{(-1)^{3+7}M_{73}}{|J|} \propto -(U^* - \hat{U})\frac{M_{73}}{|J|} \tag{S65}$$

and

$$\frac{\partial}{\partial q}\left(\frac{I_1^*}{N_1^*}\right) = \frac{\partial}{\partial q}\left(\frac{I_1^*}{S_1^* + I_1^* + R_1^*}\right) \propto -(U^* - \hat{U})\frac{\partial}{\partial \delta}\left(\frac{I_1^*}{S_1^* + I_1^* + R_1^*}\right) \tag{S66}$$

where $M_{73}$ is given in equation (S75).

Negative density dependence: The sign of equation (S66) is the same as its counterpart for the SI model (see section S1.6.4). Specifically, if interspecific competition is sufficiently low, then the sign of equation (S66) is $U^* - \hat{U}$. In the limit of very strong interspecific competition, equation (S66) has the same sign as $-(U^* - \hat{U})$. In general, increased interspecific competition can cause the sign of equation (S66) to change.

The biological interpretation of the above is the same as in the SI model:

- Assume interspecific competition between hosts is less than intraspecific competition and infected individuals are weaker competitors than susceptible individuals
  - Focal host infection prevalence at the multi-species equilibrium is more likely to be lower under FDDT than DDDT when (D1) interspecific competition is weaker, (D2) the second host is a weaker intraspecific competitor, and (D3) the second host is a lower competence host.
  - Focal host infection prevalence at the multi-species equilibrium is more likely to be lower under DDDT than FDDT when (D4) interspecific competition is stronger, (D5) the second host is a stronger intraspecific competitor, and (D6) the second host is a higher competence host.

- Assume interspecific competition between hosts is greater than intraspecific competition or infected individuals are stronger competitors than susceptible individuals
  - Focal host infection prevalence at the multi-species equilibrium is more likely to be lower under FDDT than DDDT when when (F1) the second host is a stronger intraspecific competitor and (F2) the second host is a higher competence host.
  - Focal host infection prevalence at the multi-species equilibrium is more likely to be lower under DDDT than FDDT when when (F3) the second host is a weaker intraspecific competitor and (F4) the second host is a lower competence host.

As noted in S1.5.4, conditions D1-D6 are the same for the transformation that holds the single-species equilibrium constant (this section) and the transformation that holds the multi-species equilibrium constant (previous section). In addition, conditions F1-F4 differ because $\partial(I_1^*/N_1^*)/\partial\delta$ is positive, i.e., increased removal of infectious propagules causes focal host infection prevalence to increase.

Finally, if interspecific competition is absent and there is negative density dependence in the focal host, it is not possible for the focal host to experience amplification at $q = 0$ and dilution at $q = 1 + \hat{U}/\delta$. The proof is similar to that for the SI model. To prove it by contradiction, we assume negative density dependence. Amplification at $q = 0$ and dilution at $q = 1 + \hat{U}/\delta$ is only possible if $U^* - \hat{U} < 0$, which implies $S_1^* - \hat{S}_1 < 0$ and sufficiently large for $0 \leq q \leq 1 + \hat{U}/\delta$. We now compute the dependencies of $S_1^*$ and $R_1^*$ on $I_1^*$. From $0 = dR/dt$ we have $\partial R_1^*/\partial I_1^* = \nu_1/(\gamma_1 + m_1) > 0$. Because interspecific competition is absent, the equilibrium host densities satisfy $0 = f_1(S_1^*, I_1^*, R_1^*) - \mu_1 I_1^* - m_1 R_1^*$. The dependence of $S_1^*$ on $I_1^*$ can be computed by implicitly differentiating that equation to get,

$$0 = \frac{\partial f_1}{\partial S_1}\frac{\partial S_1^*}{\partial I_1^*} + \frac{\partial f_1}{\partial R_1}\frac{\partial R_1^*}{\partial I_1^*} + \frac{\partial f_1}{\partial I_1} - \mu_1 - m_1\frac{\partial R_1^*}{\partial I_1^*}\bigg|_{p^*} \tag{S67}$$

$$\Rightarrow \quad \frac{\partial S_1^*}{\partial I_1^*} = -\left(\frac{\partial f_1}{\partial I_1} + \frac{\partial f_1}{\partial R_1}\frac{\nu_1}{\gamma_1 + m_1} - \mu_1 - m_1\frac{\nu_1}{\gamma_1 + m_1}\right)\bigg/\frac{\partial f_1}{\partial S_1}\bigg|_{p^*} \tag{S68}$$

The assumed negative density dependence implies $\frac{\partial S_1^*}{\partial I_1^*}$ is negative. Thus, increases in $I_1^*$ imply decreases in $S_1^*$ and vice versa. Combining $\frac{\partial S_1^*}{\partial I_1^*} < 0$, $\frac{\partial R_1^*}{\partial I_1^*} > 0$ and $S_1^* - \hat{S}_1 < 0$ yields $S_1^* - \hat{S}_1 < 0$, $I_1^* - \hat{I}_1 > 0$, and $R_1^* - \hat{R}_1 > 0$ for all $0 \leq q \leq 1 + \hat{U}/\delta$, which implies $I_1^*/(S_1^* + I_1^* + R_1^*) > \hat{I}_1/(\hat{S}_1 + \hat{I}_1 + \hat{R}_1)$ for $0 \leq q \leq 1 + \hat{U}/\delta$. However, this means that the proportion of infected individuals at the multi-species equilibrium is higher than the proportion of infected individuals at the single-species equilibrium for all values of $q$, which contradicts our assumption that dilution occurs at $q = 1 + \hat{U}/\delta$. Thus, in the absence of interspecific host competition, it is not possible for amplification to occur under frequency-dependent direct transmission ($q = 0$) and dilution to occur under density-dependent direct transmission ($q = 1 + \hat{U}/\delta$).

Positive density dependence or low loss of immunity rates: If either host has positive density dependence at equilibrium and the positive density dependence is sufficiently large, then the above conditions can be reversed.

#### S2.3 Response of single-species equilibrium prevalence to parameter variation

All terms and derivatives in this section are evaluated at $\hat{p}$. Recall that $|\hat{J}| > 0$.

The responses to increased infectious propagule degradation are

$$\frac{\partial \hat{S}_1}{\partial \delta} \propto (-1)^{1+4}\frac{M_{41}}{|\hat{J}|} = \frac{\beta_1 \hat{S}_1}{|\hat{J}|}\left[(\gamma_1 + m_1)\left(\frac{\partial f_1}{\partial I_1} - (\mu_1 + \nu_1)\right) + \nu_1\left(\frac{\partial f_1}{\partial R_1} + \gamma_1\right)\right] \tag{S69}$$

$$\frac{\partial \hat{I}_1}{\partial \delta} \propto (-1)^{2+4}\frac{M_{42}}{|\hat{J}|} = \frac{(\gamma_1 + m_1)\beta_1 \hat{S}_1}{|\hat{J}|}\frac{\partial f_1}{\partial S_1}, \tag{S70}$$

$$\frac{\partial \hat{R}_1}{\partial \delta} \propto (-1)^{3+4}\frac{M_{43}}{|\hat{J}|} = \frac{\nu_1 \beta_1 \hat{S}_1}{|\hat{J}|}\frac{\partial f_1}{\partial S_1}, \tag{S71}$$

and

$$\begin{aligned}\frac{\partial(\hat{I}_1/\hat{N}_1)}{\partial \delta} \propto \frac{\hat{S}_1^2 \beta_1}{(\mu_1 + \nu_1)|\hat{J}|\hat{N}_1^2}&\left[(\gamma_1 + m_1)(\mu_1 + \nu_1)\left(\frac{\partial f_1}{\partial S_1} - \beta_2 \hat{P}\right)\right.\\ &\left. + \hat{P}\beta_1(\gamma_1 + m_1)\frac{\partial f_1}{\partial I_1} + \hat{P}\beta_1 \nu_1\left(\frac{\partial f_1}{\partial R_1} + \gamma_1\right)\right].\end{aligned} \tag{S72}$$

The sign of $\partial(\hat{I}_1/\hat{N}_1)/\partial\delta$ is negative under the assumption of negative density dependence. Sufficiently large positive density dependence in the infected or recovered classes can cause the derivative to be positive.

#### S2.4 Response of the multi-species equilibrium prevalence to parameter variation

All terms and derivatives in the following subsections are evaluated at $p^*$. In nearly all cases, the results are identical to those for the SI model. Specific differences are pointed out in each section and summarized in section S2.4.5. We do not present any results about responses to changes in the transmission coefficient for the second host ($\beta_2$) because the equations were too complex to interpret, but our numerical simulations suggest that there are no qualitative differences between results for the SI and SIR models.

To compute how the proportion of infected individuals ($I_1/N_1 = I_1/[S_1 + I_1]$) responds to a small change in parameter $a$, we use the chain rule,

$$\begin{aligned}\frac{\partial}{\partial a}\left(\frac{I_1^*}{N_1^*}\right) &= \frac{1}{(N_1^*)^2}\left[(S_1^* + I_1^* + R_1^*)\frac{\partial I_1^*}{\partial a} - I_1^*\left(\frac{\partial S_1^*}{\partial a} + \frac{\partial I_1^*}{\partial a} + \frac{\partial R_1^*}{\partial a}\right)\right]\\ &= \frac{1}{(N_1^*)^2}\left[(S_1^* + R_1^*)\frac{\partial I_1^*}{\partial a} - I_1^*\left(\frac{\partial S_1^*}{\partial a} + \frac{\partial R_1^*}{\partial a}\right)\right]\\ &= \frac{1}{(N_1^*)^2}\left[\left(S_1^* + \frac{\nu_1\beta_1 S_1^* P^*}{(\mu_1+\nu_1)(\gamma_1+m_1)}\right)\frac{\partial I_1^*}{\partial a} - \frac{\beta_1 P^*}{\mu_1+\nu_1}\left(\frac{\partial S_1^*}{\partial a} + \frac{\partial R_1^*}{\partial a}\right)\right]\end{aligned} \tag{S73}$$

where the last line uses the equilibrium conditions $0 = dI_i/dt|_{p^*}$ and $0 = dR_i/dt|_{p^*}$ to substitute for $I_1^*$ and $R_1^*$.

The forms of all derivatives are

$$\frac{\partial I_1^*}{\partial a} = (\gamma_1 + m_1)(\gamma_2 + m_2)(\text{“terms in SI model derivative”}) + \text{“new terms due to recovery”} \tag{S74}$$

where $a$ is a model parameter, "terms in SI model derivative" are the terms that show up in the analogous equation in appendix S1, and "new terms due to recovery" are additional terms that arise due to recovery from infection. In the equations, the terms from the SI model are denoted "RHS(eqn (SX))", where SX is the equation number in appendix S1. To conserve space, we only present the equations for the responses in the density and proportion of infected individuals. The equations for the responses in the densities of susceptible and recovered individuals can be computed in the accompanying Maple Worksheets.

When interpreting the equations it is useful to keep in mind that $|J| < 0$, $U^*$ is the total per capita infectious propagule uptake rate at the multi-species equilibrium, $U^* + \delta - u_{S_1}S_1 - u_{S_2}S_2 > 0$, $\chi_i - u_{I_2}P$ determines whether infected individuals in population $i$ are net sources (positive values) or net sinks (negative values), the effects of intraspecific competition show up through $\partial F_i/\partial X_i$ for $X \in \{S, I, R\}$, and the effects of interspecific competition show up through $\partial F_i/\partial X_j$ for $i \neq j$ and $X \in \{S, I, R\}$.

##### S2.4.1 Response to increased excretion or removal of infectious propagules

The responses to increased degradation rate of the infectious propagules are

$$
\begin{aligned}
\frac{\partial I_1^*}{\partial \delta} \propto & (-1)^{3+7}\frac{M_{73}}{|J|} = (\gamma_1+m_1)(\gamma_2+m_2)\,RHS(\text{eqn (S42)}) \\
& + (\gamma_1+m_1)\Bigg[\frac{PS_1\beta_1\beta_2\nu_2}{|J|}\frac{\partial f_1}{\partial S_1}\left(\frac{\partial f_2}{\partial R_2}+\gamma_2\right) \\
& - \frac{P\beta_1\beta_2\nu_2}{|J|}\frac{\partial f_1}{\partial R_2}\left(S_1\frac{\partial f_2}{\partial S_1}+S_2\frac{\partial f_2}{\partial S_2}\right) + \frac{PS_2\beta_1\beta_2\nu_2}{|J|}\frac{\partial f_1}{\partial S_2}\left(\frac{\partial f_2}{\partial R_2}+\gamma_2\right)\Bigg]
\end{aligned}
\tag{S75}
$$

and

$$
\begin{aligned}
\frac{\partial(I_1^*/N_1^*)}{\partial \delta} \propto & \,(\gamma_1+m_1)(\gamma_2+m_2)\,RHS(\text{eqn (S44)}) \\
& + (\gamma_1+m_1)\frac{S_1^2\beta_1P\beta_2\nu_2}{(\mu_1+\nu_1)|J|N_1^2}\left(\frac{\partial f_2}{\partial R_2}+\gamma_2\right)\left[P\beta_1\frac{\partial f_1}{\partial I_1}-(\mu_1+\nu_1)\left(P\beta_1-\frac{\partial f_1}{\partial S_1}\right)\right] \\
& (\gamma_2+m_2)\frac{S_1^2\beta_1^2P\nu_1}{(\mu_1+\nu_1)|J|N_1^2}\left(\frac{\partial f_1}{\partial R_1}+\gamma_1\right)\left[P\beta_2\frac{\partial f_2}{\partial I_2}-(\mu_2+\nu_2)\left(P\beta_2-\frac{\partial f_2}{\partial S_2}\right)\right] \\
& + \frac{S_1^2\beta_1^2P^2\beta_2\nu_1\nu_2}{(\mu_1+\nu_1)|J|N_1^2}\left(\frac{\partial f_2}{\partial R_2}+\gamma_2\right)\left(\frac{\partial f_1}{\partial R_1}+\gamma_1\right) \\
& - (\gamma_1+m_1)\frac{S_1^2\beta_1P\beta_2\nu_2}{(\mu_1+\nu_1)|J|N_1^2}\frac{\partial f_1}{\partial R_2}\left[P\beta_1\frac{\partial f_2}{\partial I_1}+\frac{\partial f_2}{\partial S_1}(\mu_1+\nu_1)\right] \\
& - (\gamma_2+m_2)\frac{S_1^2\beta_1^2P\nu_1}{(\mu_1+\nu_1)|J|N_1^2}\frac{\partial f_2}{\partial R_1}\left[P\beta_2\frac{\partial f_1}{\partial I_2}+\frac{\partial f_1}{\partial S_2}(\mu_2+\nu_2)\right] \\
& - \frac{S_1^2\beta_1^2P^2\beta_2\nu_1\nu_2}{(\mu_1+\nu_1)|J|N_1^2}\frac{\partial f_1}{\partial R_2}\frac{\partial f_2}{\partial R_1}
\end{aligned}
\tag{S76}
$$

<u>Negative density dependence and low loss of immunity rates:</u> The biological interpretation of equation (S76) is the same as the equation for the SI model. The intraspecific competition terms in equation (S76) are negative and the interspecific competition terms are positive. Thus, increased degradation causes the proportion of infected individuals to decrease unless (1) interspecific competition is greater than intraspecific competition for all host classes, (2) interspecific competition between infected and susceptible individuals is greater than both intraspecific competition between infected and susceptible individuals and interspecific competition between susceptible individuals, or (3) interspecific competition between infected and recovered individuals is greater than both intraspecific competition between infected and susceptible individuals and interspecific competition between susceptible individuals.

<u>Positive density dependence:</u> If one or both hosts are experiencing positive density dependence and the positive density dependence is sufficiently weak, then the above results apply. If the positive density dependence is sufficiently strong, then any of the above predictions can be reversed.

**Response to increased uptake of infectious propagules:** Because $\partial(dP/dt)/\partial\delta$ and $\partial(dP/dt)/\partial u_{ij}$ have the same sign, the signs of $\partial(I_1^*/N_2^*)/\partial u_{ij}$ and $\partial(I_1^*/N_2^*)/\partial\delta$ are the same.

**Response to increased release of infectious propagules:** Because $\partial(dP/dt)/\partial\delta$ and $\partial(dP/dt)/\partial\chi_i$ have opposite signs, $\partial(I_1^*/N_2^*)/\partial\chi_i$ and $\partial(I_1^*/N_2^*)/\partial\delta$ have opposite signs.

##### S2.4.2 Response to increased disease-induced mortality of the second host

The responses to increases in the mortality rate of $I_2$ are

$$\begin{aligned}\frac{\partial I_1^*}{\partial\mu_2} &\propto (-1)^{4+3}\frac{M_{43}}{|J|} = (\gamma_1+m_1)(\gamma_2+m_2)\,RHS(\text{eqn (S45)}) + (\gamma_1+m_1)\Bigg[\\ &\frac{\beta_1\nu_2 u_{R_2} S_1 P}{|J|}\frac{\partial f_1}{\partial S_1}\left(P\beta_2-\frac{\partial f_2}{\partial S_2}\right)+\frac{\beta_1\nu_2 u_{S_2} S_1 P}{|J|}\frac{\partial f_1}{\partial S_1}\left(\frac{\partial f_2}{\partial R_2}+\gamma_2\right)\\ &-\frac{\beta_1\nu_2 S_1 P}{|J|}\frac{\partial f_2}{\partial S_1}\left(\frac{\partial f_1}{\partial R_2}u_{S_2}-\frac{\partial f_1}{\partial S_2}u_{R_2}\right)-\frac{\beta_1\nu_2 P}{|J|}\frac{\partial f_1}{\partial R_2}\frac{\partial f_2}{\partial S_2}\left(-S_1u_{S_1}+U+\delta\right)\\ &+\frac{\beta_1\beta_2\nu_2P^2}{|J|}\frac{\partial f_1}{\partial R_2}\left(-S_1u_{S_1}-S_2u_{S_2}+U+\delta\right)+\frac{\beta_1\nu_2 P}{|J|}\frac{\partial f_1}{\partial S_2}\frac{\partial f_2}{\partial R_2}\left(-S_1u_{S_1}+U+\delta\right)\\ &+\frac{\beta_1\nu_2 P}{|J|}\frac{\partial f_1}{\partial S_2}\left(I_2[\mu_2+\gamma_2]u_{R_2}-S_1\gamma_2u_{S_1}+U+\delta\gamma_2\right)\Bigg]\end{aligned}\tag{S77}$$

and

$$\begin{aligned}&\frac{\partial(I_1^*/N_1^*)}{\partial\mu_2}\propto(\gamma_1+m_1)(\gamma_2+m_2)\,RHS(\text{eqn (S47)})\\ &+(\gamma_1+m_1)\frac{S_1^2\beta_1P\nu_2}{(\mu_1+\nu_1)|J|N_1^2}\left[u_{R_2}\left(P\beta_2-\frac{\partial f_2}{\partial S_2}\right)+u_{S_2}\left(\frac{\partial f_2}{\partial R_2}+\gamma_2\right)\right]\left[\beta_1P\frac{\partial f_1}{\partial I_1}-(\mu_1+\nu_1)\left(P\beta_1-\frac{\partial f_1}{\partial S_1}\right)\right]\\ &-(\gamma_2+m_2)\frac{S_1^2\beta_1^2P\nu_1}{(\mu_1+\nu_1)|J|N_1^2}\left(\frac{\partial f_1}{\partial R_1}+\gamma_1\right)\left[(\chi_2-u_{I_2}P)\left(P\beta_2-\frac{\partial f_2}{\partial S_2}\right)-P\frac{\partial f_2}{\partial I_2}u_{S_2}\right]\\ &+\frac{S_1^2\beta_1^2P^2\nu_1\nu_2}{(\mu_1+\nu_1)|J|N_1^2}\left(\frac{\partial f_1}{\partial R_1}+\gamma_1\right)\left[u_{R_2}\left(P\beta_2-\frac{\partial f_2}{\partial S_2}\right)+u_{S_2}\left(\frac{\partial f_2}{\partial R_2}+\gamma_2\right)\right]\\ &+(\gamma_1+m_1)\frac{S_1^2\beta_1P\nu_2}{(\mu_1+\nu_1)|J|N_1^2}\Bigg[\left(\frac{\partial f_1}{\partial R_2}\left[P\beta_2-\frac{\partial f_2}{\partial S_2}\right]+\frac{\partial f_1}{\partial S_2}\left[\frac{\partial f_2}{\partial R_2}+\gamma_2\right]\right)\left([\chi_1-u_{I_1}P]\beta_1-[\mu_1+\nu_1]\,u_{S_1}\right)\\ &+\left(-\frac{\partial f_1}{\partial R_2}u_{S_2}+\frac{\partial f_1}{\partial S_2}u_{R_2}\right)\left(P\beta_1\frac{\partial f_2}{\partial I_1}+\frac{\partial f_2}{\partial S_1}[\mu_1+\nu_1]\right)\Bigg]\\ &-(\gamma_2+m_2)\frac{S_1^2\beta_1^2P\nu_1}{(\mu_1+\nu_1)|J|N_1^2}\left(P\frac{\partial f_1}{\partial I_2}u_{R_1}\left(P\beta_2-\frac{\partial f_2}{\partial S_2}\right)+P\frac{\partial f_1}{\partial I_2}\frac{\partial f_2}{\partial R_1}u_{S_2}+P\frac{\partial f_1}{\partial S_2}\frac{\partial f_2}{\partial I_2}u_{R_1}+(\chi_2-u_{I_2}P)\frac{\partial f_1}{\partial S_2}\frac{\partial f_2}{\partial R_1}\right)\\ &+\frac{S_1^2\beta_1^2P^2\nu_1\nu_2}{(\mu_1+\nu_1)|J|N_1^2}\left(\frac{\partial f_1}{\partial R_2}u_{R_1}\left(P\beta_2-\frac{\partial f_2}{\partial S_2}\right)+\frac{\partial f_2}{\partial R_1}\left(\frac{\partial f_1}{\partial R_2}u_{S_2}-\frac{\partial f_1}{\partial S_2}u_{R_2}\right)+\frac{\partial f_1}{\partial S_2}u_{R_1}\left(\frac{\partial f_2}{\partial R_2}+\gamma_2\right)\right)\end{aligned}\tag{S78}$$

Negative density dependence and low loss of immunity rates: The biological interpretations of equation (S78) is the same as that for the SI model. The second and third lines of equation (S78) are composed of negative terms and positive terms that can be factored with the terms in equation (S44) that start with $\chi_2 - u_{I_2}P$. Overall, this means that, in the absence of interspecific competition, prevalence in the focal host decreases with increased

$\mu_2$ unless the second host is a sufficiently large sink. The terms involving interspecific competition have mixed signs. Increased interspecific competition can cause equation (S78) to be positive if interspecific competition is greater than intraspecific competition for all host classes, or if infected or recovered individuals are stronger interspecific competitors than susceptible individuals.

Positive density dependence: Sufficiently strong positive density dependence can reverse the above predictions.

##### S2.4.3 Response to increased intraspecific competition in the second host

Let $\alpha_{22}$ be any parameter that negatively affects the reproduction rate of the second host. Mathematically, we assume $\partial f_2/\partial\alpha_{22} < 0$, which implies $\partial(dS_2/dt)/\partial\alpha_{22} < 0$. We interpret increases in $\alpha_{22}$ to mean an increase in the intraspecific competitive ability of the second host. However, our results apply to any change in parameters that causes a decrease in the growth rate of $S_2$ without affecting any other equation in the model.

The responses to the second host experiencing increased increased intraspecific competition are

$$\begin{aligned}\frac{\partial I_1^*}{\partial\alpha_{22}} \propto (-1)^{4+3}\frac{M_{43}}{|J|} = (\gamma_1+m_1)(\gamma_2+m_2)\,RHS(\text{eqn (S51)}) + (\gamma_1+m_1)\frac{\beta_1\beta_2\nu_2 u_{R_2} S_1 P^2}{|J|}\frac{\partial f_1}{\partial S_1}\\ + (\gamma_1+m_1)\frac{P^2\beta_1\beta_2\nu_2}{|J|}\left[\frac{\partial f_1}{\partial R_2}(U+\delta-S_1u_{S_1}-S_2u_{S_2}) + S_2\frac{\partial f_1}{\partial S_2}u_{R_2}\right]\end{aligned} \tag{S79}$$

and

$$\begin{aligned}\frac{\partial(I_1^*/N_1^*)}{\partial\alpha_{22}} &\propto (\gamma_1+m_1)(\gamma_2+m_2)\,RHS(\text{eqn (S53)}) + \frac{S_1^2\beta_1^2P^3\beta_2\nu_1\nu_2u_{R_2}}{(\mu_1+\nu_1)|J|N_1^2}\left(\frac{\partial f_1}{\partial R_1}+\gamma_1\right)\\ &+ (\gamma_1+m_1)\frac{S_1^2\beta_1P^2\beta_2\nu_2u_{R_2}}{(\mu_1+\nu_1)|J|N_1^2}\left[P\beta_1\frac{\partial f_1}{\partial I_1} + (\mu_1+\nu_1)\left(\frac{\partial f_1}{\partial S_1}-\beta_1P\right)\right]\\ &- (\gamma_2+m_2)\frac{S_1^2\beta_1^2P^2\nu_1\beta_2}{(\mu_1+\nu_1)|J|N_1^2}\left(\frac{\partial f_1}{\partial R_1}+\gamma_1\right)(\chi_2I_2-u_{I_2}I_2P-u_{S_2}PS_2)\\ &+ (\gamma_1+m_1)\frac{P^2S_1^2\beta_1^2\beta_2\nu_2}{(\mu_1+\nu_1)|J|N_1^2}\frac{\partial f_1}{\partial R_2}(\chi_1I_1-u_{S_2}PI_1-u_{S_1}S_1P)\\ &- (\gamma_2+m_2)\frac{S_1^2\beta_1^2P^2\nu_1u_{R_1}}{(\mu_1+\nu_1)|J|N_1^2}\left[P\beta_2\frac{\partial f_1}{\partial I_2}+\frac{\partial f_1}{\partial S_2}(\mu_2+\nu_2)\right] - \frac{S_1^2\beta_1^2P^3\beta_2\nu_1\nu_2u_{R_1}}{(\mu_1+\nu_1)|J|N_1^2}\frac{\partial f_1}{\partial R_2}\end{aligned} \tag{S80}$$

Negative density dependence: The additional terms that show up in equation (S80) have a slightly different biological interpretation as the corresponding equations for the SI model. The additional terms on the first, second and third lines of equation (S82) only involve the effects of intraspecific competition. The second line has the same sign as $-(\chi_2I_2 - u_{I_2}PI_2 - u_{S_2}S_2P)$, which is the same as the corresponding terms in equation (S53) for the SI model. The additional terms on the first and third lines are positive; there are no analogous terms in equation (S53) for the SI model. The fourth and fifth lines of equation (S82) include the effects of interspecific competition. Those terms are either negative or have the same sign as $(\chi_1 I_1 - u_{I_1} P I_1 - u_{S_1} S_1 P)$.

Overall, it is still the case that increased intraspecific competition in population 2 causes an increase in infection prevalence in population 1 unless the second host is a large source. In the absence of interspecific competition, competition with recovered individuals increases the threshold for the second host to be a sufficiently large source. However, increased interspecific competition causes that threshold to decrease.

Positive density dependence: Sufficiently strong positive density dependence can reverse the above predictions.

##### S2.4.4 Response to the focal host experiencing increased competition:

Let $\alpha_{12}$ be any parameter that negatively affects the reproduction rate of the focal host . Mathematically, we assume $\partial f_1/\partial\alpha_{12} < 0$, which implies $\partial(dS_1/dt)/\partial\alpha_{12} < 0$. We interpret increases in $\alpha_{12}$ to mean an increase in the interspecific competitive ability of the second host. However, our results apply to any change in parameters that causes a decrease in the growth rate of $S_1$ without affecting any other equation in the model.

The responses to the focal host experiencing increased competition are

$$\begin{aligned}\frac{\partial I_1^*}{\partial\alpha_{12}} \propto (-1)^{4+3}\frac{M_{43}}{|J|} &= (\gamma_1+m_1)(\gamma_2+m_2)\,RHS(\text{eqn (S54)})+\\ &-(\gamma_1+m_1)\frac{P^2\beta_1\beta_2\nu_2}{|J|}\left(\frac{\partial f_2}{\partial R_2}+\gamma_2\right)(U+\delta-S_1u_{S_1}-S_2u_{S_2})\\ &-(\gamma_1+m_1)\frac{\beta_1\beta_2\nu_2u_{R_2}S_1P^2}{|J|}\frac{\partial f_2}{\partial S_1}+-(\gamma_1+m_1)\frac{\beta_1\beta_2\nu_2u_{R_2}S_2P^2}{|J|}\frac{\partial f_2}{\partial S_2}\end{aligned} \tag{S81}$$

and

$$\begin{aligned}\frac{\partial(I_1^*/N_1^*)}{\partial\alpha_{12}} &\propto (\gamma_1+m_1)(\gamma_2+m_2)\,RHS(\text{eqn (S56)})\\ &-(\gamma_1+m_1)\frac{S_1^2\beta_1^2\beta_2\nu_2P^2}{I_1(\mu_1+\nu_1)|J|N_1^2}\left(\frac{\partial f_2}{\partial R_2}+\gamma_2\right)(\chi_1I_1-u_{I_1}PI_1-u_{S_1}S_1P)\\ &+(\gamma_2+m_2)\frac{S_1^2\beta_1^2P^2\nu_1u_{R_1}}{(\mu_1+\nu_1)|J|N_1^2}\left(P\beta_2\frac{\partial f_2}{\partial I_2}+[\mu_2+\nu_2]\left[\frac{\partial f_2}{\partial S_2}-P\beta_2\right]\right)+\frac{S_1^2\beta_1^2P^3\beta_2\nu_1\nu_2u_{R_1}}{(\mu_1+\nu_1)|J|N_1^2}\left(\frac{\partial f_2}{\partial R_2}+\gamma_2\right)\\ &-(\gamma_1+m_1)\frac{S_1^2\beta_1P^2\beta_2\nu_2u_{R_2}}{(\mu_1+\nu_1)|J|N_1^2}\left(P\beta_1\frac{\partial f_2}{\partial I_1}+\frac{\partial f_2}{\partial S_1}[\mu_1+\nu_1]\right)\\ &+(\gamma_2+m_2)\frac{S_1^2\beta_1^2\beta_2P^2\nu_1}{I_2(\mu_1+\nu_1)|J|N_1^2}\frac{\partial f_2}{\partial R_1}(\chi_2I_2-u_{I_2}PI_2-u_{S_2}S_2P)-\frac{S_1^2\beta_1^2P^3\beta_2\nu_1\nu_2}{(\mu_1+\nu_1)|J|N_1^2}\frac{\partial f_2}{\partial R_1}u_{R_2}\end{aligned} \tag{S82}$$

Negative density dependence: The additional terms that show up in equation (S82) have a slightly different biological interpretations as the corresponding equations for the SI model. The second and third lines of equation (S82) only involve the effects of intraspecific competition. The second line has the same sign as $-(\chi_1 I_1 - u_{I_1} P I_1 - u_{S_1} S_1 P)$, which is the same as the corresponding terms in equation (S56) for the SI model. The third line is positive, which is the opposite of the corresponding lines in equation (S56) for the SI model. This means that intraspecific competition with recovered individuals tends to promote increased infection prevalence with increased interspecific competition. The remaining lines involve the effects of interspecific host competition; all of these terms have the same signs as the corresponding terms in equation (S56) for the SI model.

In total, it is still that case that increased interspecific competitive ability of the second host causes infection prevalence in the focal host to decrease, unless the second host is a sufficiently large source. However, intraspecific competition with recovered individuals causes the threshold for being a sufficiently large source to decrease.

Positive density dependence: Sufficiently strong positive density dependence can reverse the above predictions.

##### S2.4.5 Comparison of responses in SI an SIRS models

The only qualitative difference between the SI and SIRS models is that positive density dependence can arise in the SIRS model when recovered individuals are weak intraspecific competitors and loss of immunity rates are large. Otherwise, our conclusions about how the characteristics of the second host affect the disease prevalence the focal host are qualitatively similar for the SI and SIRS models.

Including recovery and loss of immunity does have a quantitative effect of shirting the thresholds for switches from increased to decreased prevalence, i.e., an equation having a positive or negative sign. Specifically, competition with recovered individuals causes: (i) an increase in the threshold for a sufficiently large sink in order to get decreased prevalence with increased competence of the second host, (ii) a decrease in the threshold for a sufficiently large source in order to get increased prevalence with increased interspecific competitive ability of the second host, and (iii) an increase in the threshold for a sufficiently large source in order to get increased prevalence with increased interspecific competitive ability of the second host.

#### S3 Supplementary Figure

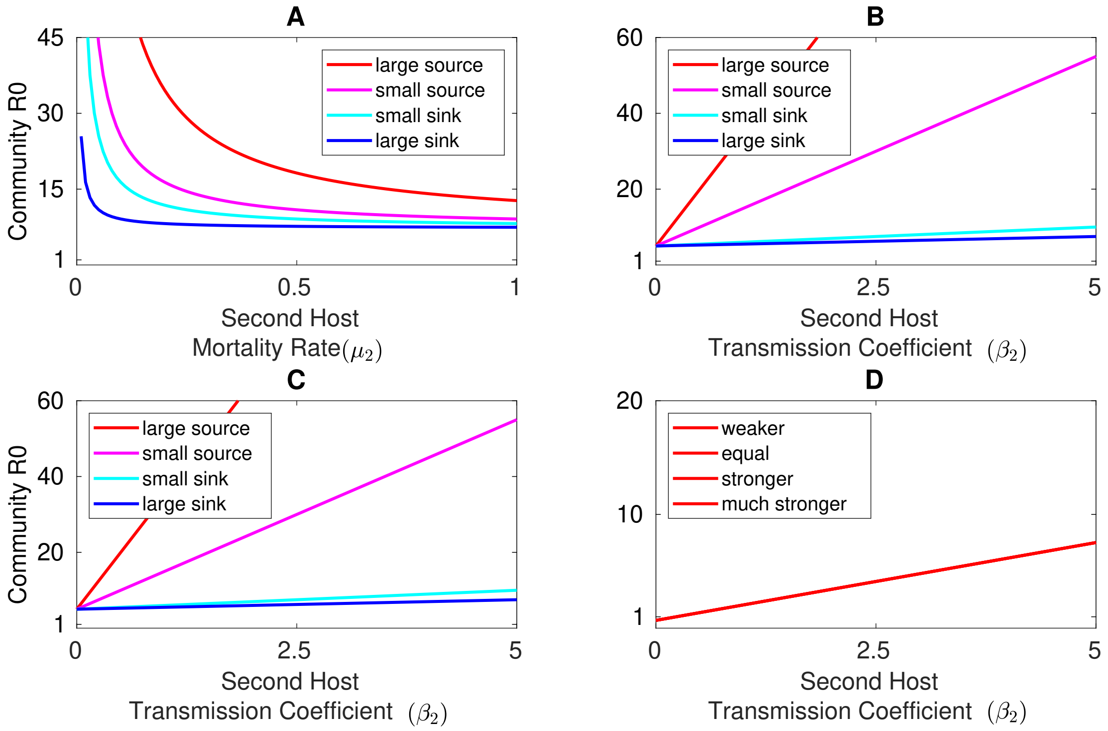

Figure S1: Increased competence of the second host increases the pathogen's community reproductive number, $\mathcal{R}_0$. Values of $\mathcal{R}_0$ were computed numerically using the next generation matrix method (Diekmann et al. 2010). Responses to (A) increased disease-induced mortality in the second host and (B,C) increases in the transmission rate of the second host when infected individuals have (B) equal and (C) higher uptake rates than susceptible individuals and the second host is a (blue) large sink, (cyan) small sink, (magenta) small source, or (red) large source. Panels B and C are identical because the competitive ability of infected individuals does not affect $\mathcal{R}_0$. (D) Responses to increases in the transmission rate of the second host when infected individuals are (blue) weaker, (cyan) equal, (magenta) stronger, or (red) much stronger interspecific competitors than susceptible individuals. The curves in panel D are identical because the value of $\mathcal{R}_0$ is independent of the competitive ability of infected individuals. Parameters are the same as in Figure 2 in the main text; see appendix S1.6 for equations and parameters.

### S4 Figure equations and parameters

For all figures, the model equations are

$$\begin{aligned}
\frac{dS_i}{dt} &= \underbrace{\left[r_i(S_i + c_i I_i)[1 - \alpha_{i1}(S_1 + e_{i1}I_1) - \alpha_{i2}(S_2 + e_{i2}I_2)]\right]}_{\text{growth \& competition}} - \underbrace{\beta_i P S_i}_{\text{infection}} - \underbrace{m_i S_i}_{\text{mortality}} \\
\frac{dI_i}{dt} &= \underbrace{\beta_i S_i P}_{\text{infection}} \underbrace{-(m_i + \mu_i)I_i}_{\text{mortality}} \\
\frac{dP}{dt} &= \underbrace{\chi_1 I_1 + \chi_2 I_2}_{\text{propagule excretion}} \underbrace{-(u_{S_1}S_1 + u_{I_1}I_1 + u_{S_2}S_2 + u_{I_2}I_2)P}_{\text{propagule uptake}} \underbrace{-\delta P}_{\text{degradation}} .
\end{aligned} \tag{S83}$$

where $r_i$ and $c_i r_i$ (with $c_i \leq 1$) are the maximum reproduction rates of susceptible and infected individuals of species $i$, $\alpha_{ij}$ is the per capita competitive effect of host $j$ on host $i$, and $e_{ij}$ determines whether infected individuals of host $j$ have weaker ($e_{ij} < 1$), equal ($e_{ij} = 1$), or stronger ($e_{ij} > 1$) competitive effects on host $i$ than susceptible individuals of host $j$; all other parameters are defined as in model (1) of the main text.

For Figure 1, the infectious propagule equation is

$$\frac{dP}{dt} = \underbrace{\chi_1 I_1 + \chi_2 I_2}_{\text{propagule excretion}} \underbrace{-f(q)(u_{S_1}S_1 - u_{I_1}I_1 - u_{S_2}S_2 - u_{I_2}I_2)P}_{\text{propagule uptake}} \underbrace{-q\delta P}_{\text{degradation}} \tag{S84}$$

where $f(q) = 1 + \frac{\delta}{\hat{U}} - \frac{\delta}{\hat{U}}q$ is a change of parameters defined on $0 \leq q \leq 1 + \hat{U}/\delta$ that transforms the environmental transmission model from a form that behaves like a frequency-dependent direct transmission model ($q = 0$) to a form that behaves like a density-dependent direct transmission model ($q = 1 + \hat{U}/\delta$), while leaving the densities at the single-species equilibrium for the focal host unchanged; see appendix S1.5.4 for details.

**Figure 1B,C:**

High, Weak (dashed cyan): $r_1 = 3$, $r_2 = 3$, $c_1 = 0$, $c_2 = 0$, $\alpha_{11} = 1$, $\alpha_{22} = 0.1$, $a_{ij} = 1$, $\beta_1 = 1.5$, $\beta_2 = 1.6$, $m_1 = m_2 = 0$, $\mu_1 = 0.1$, $\mu_2 = 0.1$, $\chi_1 = 1$, $\chi_2 = 3$, $u_{S_1} = 8$, $u_{I_1} = 1$, $u_{S_2} = 1$, $u_{I_2} = 3$, and $\delta = 3$. $\alpha_{21} = 0$ and $\alpha_{12} = 0$ in panel B and $\alpha_{21} = 0.9$ and $\alpha_{12} = 0.001$ in panel C. $\hat{U} = 4.44$ in the change of parameters function.

Low, Weak (dashed magenta): $r_1 = 2$, $r_2 = 2$, $\alpha_{11} = 1$, $\alpha_{22} = 0.1$, $a_{ij} = 1$, $\beta_1 = 2$, $\beta_2 = 1/3$, $m_1 = m_2 = 0$, $\mu_1 = 0.1$, $\mu_2 = 0.1$, $\chi_1 = 2$, $\chi_2 = 2$, $u_i = 3$ for $X \in \{S, I\}$, and $\delta = 1$. $\alpha_{21} = 0$ and $\alpha_{12} = 0$ in panel B and $\alpha_{21} = 0.9$ and $\alpha_{12} = 0.05$ in panel C. $\hat{U} = 1.85$ in the change of parameters function.

High, Strong (solid cyan): $r_1 = 0.1$, $r_2 = 0.1$, $c_1 = 0$, $c_2 = 0$, $\alpha_{11} = 1$, $\alpha_{22} = 10$, $a_{ij} = 1$, $\beta_1 = 4$, $\beta_2 = 4$, $m_1 = m_2 = 0$, $\mu_1 = 0.1$, $\mu_2 = 0.1$, $\chi_1 = 2$, $\chi_2 = 4$, $u_i = 3$ for $X \in \{S, I\}$, and $\delta = 20$. $\alpha_{21} = 0$ and $\alpha_{12} = 0$ in panel A and $\alpha_{21} = 0.5$ and $\alpha_{12} = 9$ in panel B. $\hat{U} = 1.26$ in the change of parameters function.

Low, Strong (solid magenta): $r_1 = 2$, $r_2 = 2$, $c_1 = 0$, $c_2 = 0$, $\alpha_{11} = 1$, $\alpha_{22} = 10$, $a_{ij} = 1$, $\beta_1 = 1$, $\beta_2 = 1/3$, $m_1 = m_2 = 0$, $\mu_1 = 0.1$, $\mu_2 = 0.1$, $\chi_1 = 2$, $\chi_2 = 2$, $u_{X_i} = 3$ for $X \in \{S, I\}$, and $\delta = 10$. $\alpha_{21} = 0$ and $\alpha_{12} = 0$ in panel A and $\alpha_{21} = 0.4$ and $\alpha_{12} = 4$ in panel B. $\hat{U} = 2.92$

in the change of parameters function.

**Figure 2A:** $r_1 = 1$, $r_2 = 1$, $c_1 = 0.1$, $c_2 = 0.1$, $\alpha_{11} = 1$, $\alpha_{21} = 0$, $\alpha_{12} = 0$, $\alpha_{22} = 1$, $a_{ij} = 1$, $\beta_1 = 1$, $\beta_2 = 1$, $m_1 = m_2 = 0$, $\mu_1 = 0.25$, $\chi_1 = 2$, $u_{X_i} = 1$ for $X \in \{S, I\}$, and $\delta = 0.1$. The values of $\chi_2$ are (blue) $\chi_2 = 0.2$, (cyan) $\chi_2 = 1$, (magenta) $\chi_2 = 2$, and (red) $\chi_2 = 6$.
**Figure 2B:** $r_1 = 2$, $r_2 = 2$, $c_1 = 0$, $c_2 = 0$, $\alpha_{11} = 1$, $\alpha_{21} = 0$, $\alpha_{12} = 0$, $\alpha_{22} = 1$, $a_{ij} = 1$, $\beta_1 = 0.5$, $m_1 = m_2 = 0$, $\mu_1 = 0.1$, $\mu_2 = 0.1$, $\chi_1 = 2$, $u_{X_i} = 3$ for $X \in \{S, I\}$, and $\delta = 1$. The values of $\chi_2$ are (blue) $\chi_2 = 0.1$, (cyan) $\chi_2 = 0.2$, (magenta) $\chi_2 = 2$, and (red) $\chi_2 = 6$.
**Figure 2C:** $r_1 = 2$, $r_2 = 2$, $c_1 = 0$, $c_2 = 0$, $\alpha_{11} = 1$, $\alpha_{21} = 0$, $\alpha_{12} = 0$, $\alpha_{22} = 1$, $a_{ij} = 1$, $\beta_1 = 0.5$, $m_1 = m_2 = 0$, $\mu_1 = 0.1$, $\mu_2 = 0.1$, $\chi_1 = 2$, $u_{X_i} = 6$ for $X \in \{S, I\}$, and $\delta = 1$. The values of $\chi_2$ are (blue) $\chi_2 = 0.1$, (cyan) $\chi_2 = 0.2$, (magenta) $\chi_2 = 2$, and (red) $\chi_2 = 6$.

**Figure 2D:** $r_1 = 2$, $r_2 = 2$, $c_1 = 0$, $c_2 = 0$, $\alpha_{11} = 1$, $\alpha_{21} = 0.8$, $\alpha_{12} = 0.8$, $\alpha_{22} = 1$, $a_{11} = 1$, $a_{22} = 1$, $\beta_1 = 0.5$, $m_1 = m_2 = 0$, $\mu_1 = 0.1$, $\mu_2 = 0.1$, $\chi_1 = 2$, $\chi_2 = 2$, $u_{X_i} = 2$ for $X \in \{S, I\}$, and $\delta = 7$. The values of $a_{12}$ and $a_{21}$ are (blue) $a_{12} = a_{21} = 0.5$, (cyan) $a_{12} = a_{21} = 1$, (magenta) $a_{12} = a_{21} = 1.5$, and (red) $a_{12} = a_{21} = 2.5$.

**Figure 4A:** $r_1 = 2$, $r_2 = 2$, $c_1 = 0$, $c_2 = 0$, $\alpha_{11} = 1$, $\alpha_{21} = 0$, $\alpha_{12} = 0$, $a_{ij} = 1$, $\beta_1 = 0.5$, $\beta_2 = 0.5$, $m_1 = m_2 = 0$, $\mu_1 = 0.1$, $\mu_2 = 0.1$, $\chi_1 = 2$, $u_{X_i} = 2$ for $X \in \{S, I\}$, and $\delta = 1$. The values of $\chi_2$ are (blue) $\chi_2 = 0.5$, (cyan) $\chi_2 = 1.5$, and (red) $\chi_2 = 3$.
**Figure 4B:** Same as Figure 3A except $\alpha_{21} = 0.5$ and $\alpha_{12} = 0.5$.
**Figure 4C:** $r_1 = 2$, $r_2 = 2$, $c_1 = 0$, $c_2 = 0$, $\alpha_{11} = 1$, $\alpha_{21} = 0.5$, $\alpha_{22} = 1$, $a_{ij} = 1$, $\beta_1 = 0.5$, $\beta_2 = 0.5$, $m_1 = m_2 = 0$, $\mu_1 = 0.1$, $\mu_2 = 0.1$, $\chi_1 = 2$, $u_{X_i} = 2$ for $X \in \{S, I\}$, and $\delta = 1$. The values of $\chi_2$ are (blue) $\chi_2 = 1$, (cyan) $\chi_2 = 2$, (magenta) $\chi_2 = 4$, and (red) $\chi_2 = 3$.
